# Dual contributions of cerebellar-thalamic networks to learning and offline consolidation of a complex motor task

**DOI:** 10.1101/2020.08.27.270330

**Authors:** Andres P Varani, Romain W Sala, Caroline Mailhes-Hamon, Jimena L Frontera, Clément Léna, Daniela Popa

## Abstract

The contribution of cerebellum to motor learning is often considered to be limited to adaptation, a short-timescale tuning of reflexes and previous learned skills. Yet, the cerebellum is reciprocally connected to two main players of motor learning, the motor cortex and the basal ganglia, via the ventral and midline thalamus respectively. Here, we evaluated the contribution of cerebellar neurons projecting to these thalamic nuclei in a skilled locomotion task in mice. In the cerebellar nuclei, we found task-specific neuronal activities during the task, and lasting changes after the task suggesting an offline processing of task-related information. Using pathway-specific inhibition, we found that dentate neurons projecting to the midline thalamus contribute to learning and retrieval, while interposed neurons projecting to the ventral thalamus contribute to the offline consolidation of savings. Our results thus show that two parallel cerebello-thalamic pathways perform distinct computations operating on distinct timescales in motor learning.

## INTRODUCTION

Learning to execute and automatize certain actions is essential for survival and animals have indeed the ability to learn complex patterns of movement with great accuracy to improve the outcomes of their actions(Krakauer et al., 2019). The acquisition of a motor skill is often divided into at least two components (Krakauer et al., 2019; Seidler, 2010): 1) sequence learning, which is needed when series of distinct actions are required, and 2) adaptation which corresponds to the ability to adapt a previous competence, and typically takes place when motor actions yield unexpected sensory outcomes. The neurobiological substrate of motor skills involves neurons distributed in the cortex, basal ganglia and cerebellum, which each implement a distinct learning algorithm (Doya, 1999). Supervised learning takes place in the cerebellum (Raymond and Medina, 2018), and is central for the adaptation of skills such as oculomotor movements (Herzfeld et al., 2018; Nguyen-Vu et al., 2013; Yang and Lisberger, 2014), reaching (Hewitt et al., 2015), locomotion (Darmohray et al., 2019; Morton and Bastian, 2006), as well as conditioned reflexes (Clopath et al., 2014; Longley and Yeo, 2014). The cerebellum is thought to form associations between actions and predicted sensory outcome at short-time scale (typically under one second), which are seen as internal models (Ito, 2008). The involvement of the cerebellum in learning of complex actions involving sequences of movements is far less well understood and still controversial (Baetens et al., 2020; Bernard and Seidler, 2013; Krakauer et al., 2019; Seidler et al., 2002).

Motor skills are generally progressively acquired ((e.g. Karni et al., 1998)). Several phases of motor learning, with distinct behavioral and anatomo-functional hallmarks have been described: in the early phase (Acquisition), fast improvements of performance take place, but they are susceptible to interferences; following a transitional Consolidation phase, the behavior reaches a Maintenance phase, where it becomes less variable, automatic, resistant to interferences, and may rely on different sets of brain structures compared to the initial training ((e.g. Brashers-Krug et al., 1996; Korman et al., 2003; Muellbacher et al., 2002)); some of the consolidation occurs offline during rest, which may be sufficient to change the recruitment of brain regions in the task execution (Shadmehr and Holcomb, 1997). Motor memories may also persist in the form of savings, which facilitate re-learning of the task at a later point in time (Huang et al., 2011; Mauk et al., 2014). Overall, learning is a process which is distributed in time and space.

Understanding the contribution of the cerebellum to learning thus requires taking into account its integration in brain-scale circuits including the cortex and basal ganglia (Caligiore et al., 2017). In the mammalian brain, both cerebellum and basal ganglia receive the majority of their afferences from the cerebral cortex and send projections back to the cortex via anatomically and functionally segregated channels, which are relayed by mostly non-overlapping thalamic regions (Bostan et al., 2013; Hintzen et al., 2018; Proville et al., 2014). Furthermore, anatomical and functional reciprocal di-synaptic connections have been demonstrated between the basal ganglia and the cerebellum (Bostan and Strick, 2010; Carta et al., 2019). The projections from the cerebellum to motor cortex and the striatum are relayed through distinct thalamic regions, respectively the ventral thalamus and intralaminar thalamus (Chen et al., 2014; Steriade, 1995), suggesting distinct contributions of these diencephalic projections of the cerebellum.

In the present study, we hypothesized that the cerebellum may contribute to some phases of learning in a complex motor task via its projections to the motor cortex and/or the basal ganglia. We thus focused on the dentate and interposed cerebellar nuclei and their projections to the centrolateral (intralaminar) thalamus and ventral anterior lateral complex (motor thalamus), which respectively relay their activity to the striatum and the motor cortex (Chen et al., 2014; Gornati et al., 2018; Proville et al., 2014). We first looked for task-related activities in the cerebellum and thalamus using chronic *in vivo* extracellular recordings in the cerebellar nuclei, intralaminar and ventral thalamus, along the learning of the motor task. Second, we examined the contribution of cerebellar nuclei and cerebello-thalamic pathways to learning using chemogenetic disruptions either during or after the learning sessions.

## RESULTS

In the present study, we used the paradigm of the accelerating rotarod, where the animals walk on an accelerating rotating horizontal rod; the animals must develop locomotion skills to avoid falling from the rod. To examine the involvement of the cerebellum in the accelerating rotarod task along the different phases of learning, we first evaluated the neuronal activity in the cerebello-cortical and cerebello-striatal pathways which primarily involve the interposed nucleus and ventral anterior lateral thalamus (VAL), as well as the dentate nucleus and the centrolateral thalamus (CL), respectively. For this purpose, we implanted C57/Bl6J mice (n=16) with microelectrode arrays that were designed to target either the cerebellar or the thalamic nuclei. Then, mice were trained for seven consecutive days on an accelerating rotarod with seven trials a day, each daily session preceded and followed by 10 minutes of free locomotion in an open-field (Fig 1A,B). The animals showed significant learning during the first day as evidenced by an increase in the latency to fall from the rotating rod between the first and last trial (Fig 1A, bottom, Suppl. Table 1). During the second day, there was still a significant increase between the first and last trial of the session (Fig 1A, bottom, Suppl. Table 1) and also a global increase compared with the first day. The improvement in the later days was more gradual, comparison between first and last trial did not reach significance; the asymptotic values of latency to fall were reached on the 4^th^ day and maintained for the 3 following days. Based on these observations, we defined three phases of learning: Acquisition on day 1, Consolidation on days 2, 3, 4 and Maintenance on days 5, 6, 7 (Durieux et al., 2012; Yin et al., 2009).

**Figure 1.**
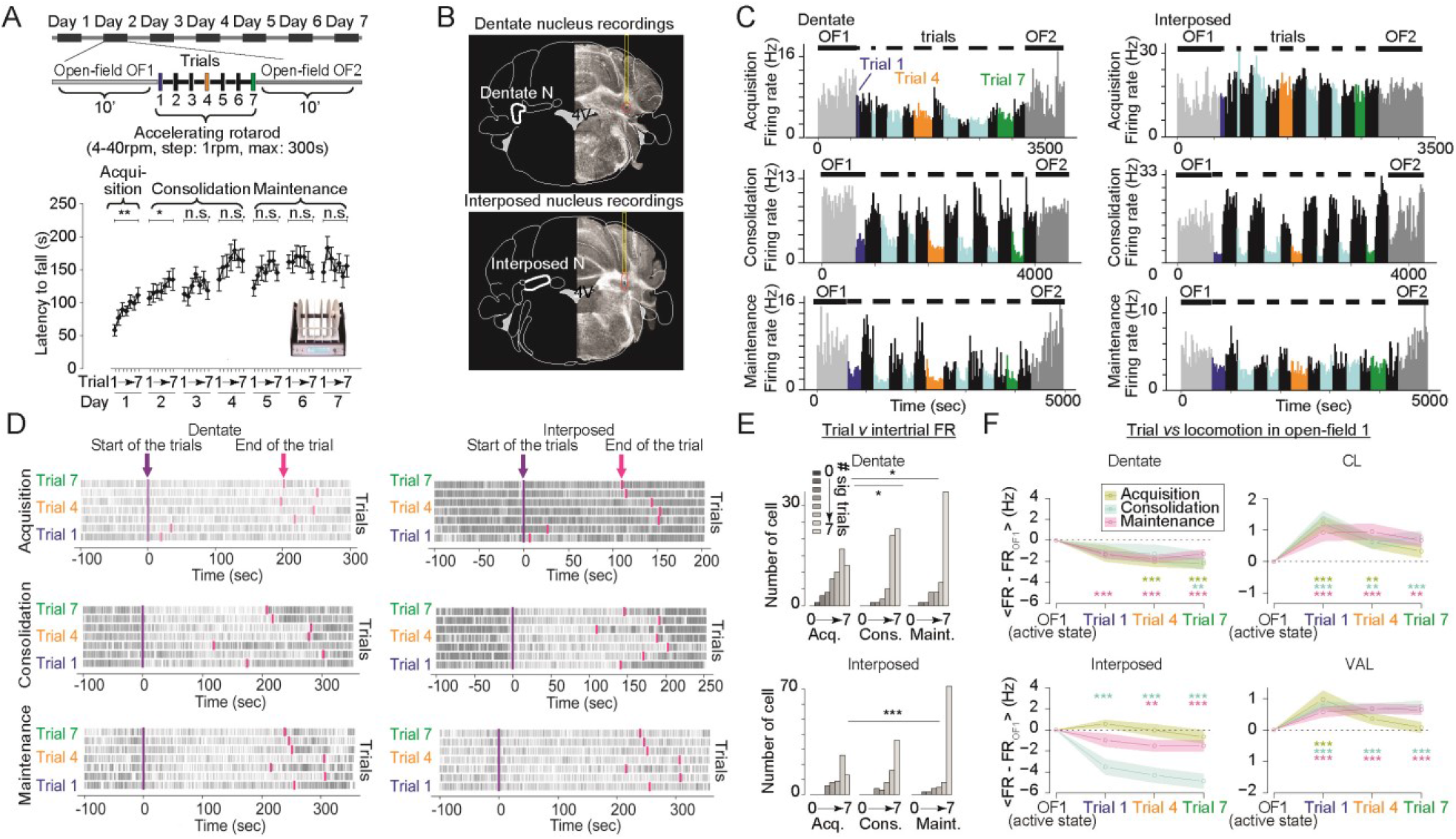
The engagement of the cerebellar nuclei during the accelerating rotarod task changes along the learning protocol. a) Scheme of the accelerating rotarod protocol (top) and latency to fall (bottom) during the accelerating rotarod across trials along days in mice recorded over the 7 days (**p<0.01, *p<0.05 t-test comparing trial 1 vs trial 7). b) Example of electrode placements in the Dentate (top) and Interposed nuclei (bottom). c) Firing rate histograms showing the evolution of activity in representative cells for Dentate (left) and Interposed (right) nuclei during Acquisition (top), Consolidation (middle) and Maintenance (bottom). Rotarod trials are shown in colors, while the open-Field before (OF1) and after (OF2) the rotarod session are shown in grey, and the intertrial periods are shown in black. d) Raster plots showing the activity of representative cells for dentate (left) and interposed (right) during trials for acquisition, consolidation and maintenance. e) Distribution of numbers of trials in which cells showed a significiant modulation of firing rate between trial and inter-trial periods during acquisition (Acq), consolidation (Cons) and maintenance (Maint) phase for Dentate (top) and Interposed (bottom) (*p<0.05, ***p<0.001 Mann-Whitney test, Holm-Sidak corrected for multiple comparison). f) Evolution of the average firing rate (mean +/− SEM), normalized by subtracting the average firing rate during the active part of the open-field session before the first rotarod trial, during Acquisition, Consolidation and Maintenance for Dentate (top left), Interposed (bottom left) nuclei, centrolateral thalamus (CL, top right), ventrolateral thalamus (VAL, bottom right) (*p<0.05, **p<0.01, ***p<0.001 Dunett Posthoc test).

We first examined to which extent the firing rate of the cerebellar nuclei (Dentate and Interposed) was modified during the running sessions on the rotarod. In most of the cells and for most of the phases of learning, the rate varied between the open-field sessions, the rotarod trials and the inter-trial episodes (Fig 1C, D). We then determined how the firing rate was changing in the rotarod apparatus depending on whether the mice were running or immobile, by comparing the distribution of instantaneous rate during each trial and the immediately preceding inter-trial during which the animal rested at the bottom of the apparatus for 300s (Fig 1E). The cells in the two cerebellar nuclei exhibited a significant difference in the distribution of instantaneous firing rate for at least one of the trials (Fig 1E, Suppl. Table 2). Moreover, we found that the number of trials with significant change in the distribution of instantaneous firing rate of the cells increased from the Acquisition phase to the later phases of learning (Fig 1E, Suppl. Table 2). This indicates that in the rotarod apparatus, the firing of Dentate and Interposed neurons was not affected only during the early phase of learning but instead showed changes along the execution of the rotarod task.

To test if the changes of firing rate reported above reflect state differences between locomotion and immobility in the rotarod apparatus or alternatively result from an engagement of the cells in the task, we compared the firing rate during the trials (1^st^, 4^th^ or 7^th^ trials) with periods of locomotion in the preceding open-field (Fig 1F). We observed that the firing rate in the Dentate nucleus was decreased in all phases, particularly in late trials (Fig 1F top left, Suppl. Table 3), while the reduction of firing in the Interposed nucleus was only limited to the last trials of Consolidation and Maintenance phase (Fig 1F bottom left, Suppl. Table 3). In a separate set of mice, we then examined the evolution of cell firing in the CL and VAL thalamus. We observed a ubiquitous increase of firing rate in CL with the exception of the last trial in the Acquisition phase (Fig 1F top right, Suppl. Table 3). Similarly the VAL thalamus showed increases of firing rate during Consolidation and Maintenance phases in all the trials, but in the Acquisition phase the increase was only observed for the first trial (Fig 1F bottom right, Suppl. Table 3). Thus the main modulation of firing rate in rotarod locomotion (compared to open-field) was a reduction in the cerebellar nuclei which developed as the learning progressed, while increased firing rates were observed for most of the CL and VAL neurons. However, the change in firing rate while walking in the open-field vs rotarod might reflect a difference in the involvement of the cells these two conditions, or more trivially it might reflect differences between the locomotion speed in the rotarod and the open-field.

To test the link between the locomotion speed and the cells’ firing, we performed a linear regression of the average firing rate as a function of the locomotion speed in the rotarod and in the open-field (Fig 2). Cells in the Dentate and Interposed nuclei exhibited negative correlation with the locomotion speed on the rotarod (Fig 2A,B, Suppl. Table 4). In contrast, on the open-field, much weaker correlations with the locomotion speed were found in the Dentate (Fig 2C, top) and Interposed (Fig 2C, bottom) cells, with significantly more negative slopes on the rotarod compared to the open-field in all learning phases (Fig 2C, Suppl. Table 5). Contrarily to the preeminence of negative correlation observed in the cerebellar nuclei in the rotarod, both positive and negative correlations were found in CL and VAL neurons (Fig Sup1A). When examining the slope of the correlations in cells with significant correlations with rotarod speed (Fig 2D), we found slightly more negative slopes in the Maintenance phase in the Dentate cells (Fig 2D, top left, Suppl. Table 6) and slightly more negative slopes in the Acquisition phase compared to the later phases in Interposed cells (Fig 2D, bottom left, Suppl. Table 6). Stronger negative slopes in the rotarod were also found in the CL thalamus during Consolidation and Maintenance phases (Fig 2D, top right, Suppl. Table 6), whereas slopes in the VAL thalamus remained stable along learning (Fig 2D, bottom right, Suppl. Table 6). We found monotonous relationships between statistically significant slopes and their associated Pearson’s correlation coefficient for Dentate (Fig 2E, top, Suppl. Table 7) and Interposed (Fig 2E, bottom, Suppl. Table 7) cells during all the phases-with the exception of Acquisition phase for the Dentate nucleus-indicating that the stronger modulations of firing rate by the locomotion speed tend to be more consistent. Similarly, we also observed monotonous relationships for CL and VAL thalamus (Fig Sup1B, Suppl. Table 7). In order to analyze the evolution of the modulation by the speed between structures, we then compared their distributions of Pearson’s correlation coefficients (which are dimensionless quantities) in all phases. We observed that Dentate-CL overlap index (ƞ) showed more similar Pearson’s correlation coefficients during Consolidation and Maintenance than in Acquisition (Fig 2F, top, Suppl. Table 8). In contrast, Interposed-VAL overlap index (ƞ) showed similar Pearson’s correlation coefficients along all phases (Fig 2F, bottom, Suppl. Table 8).

**Fig 2.**
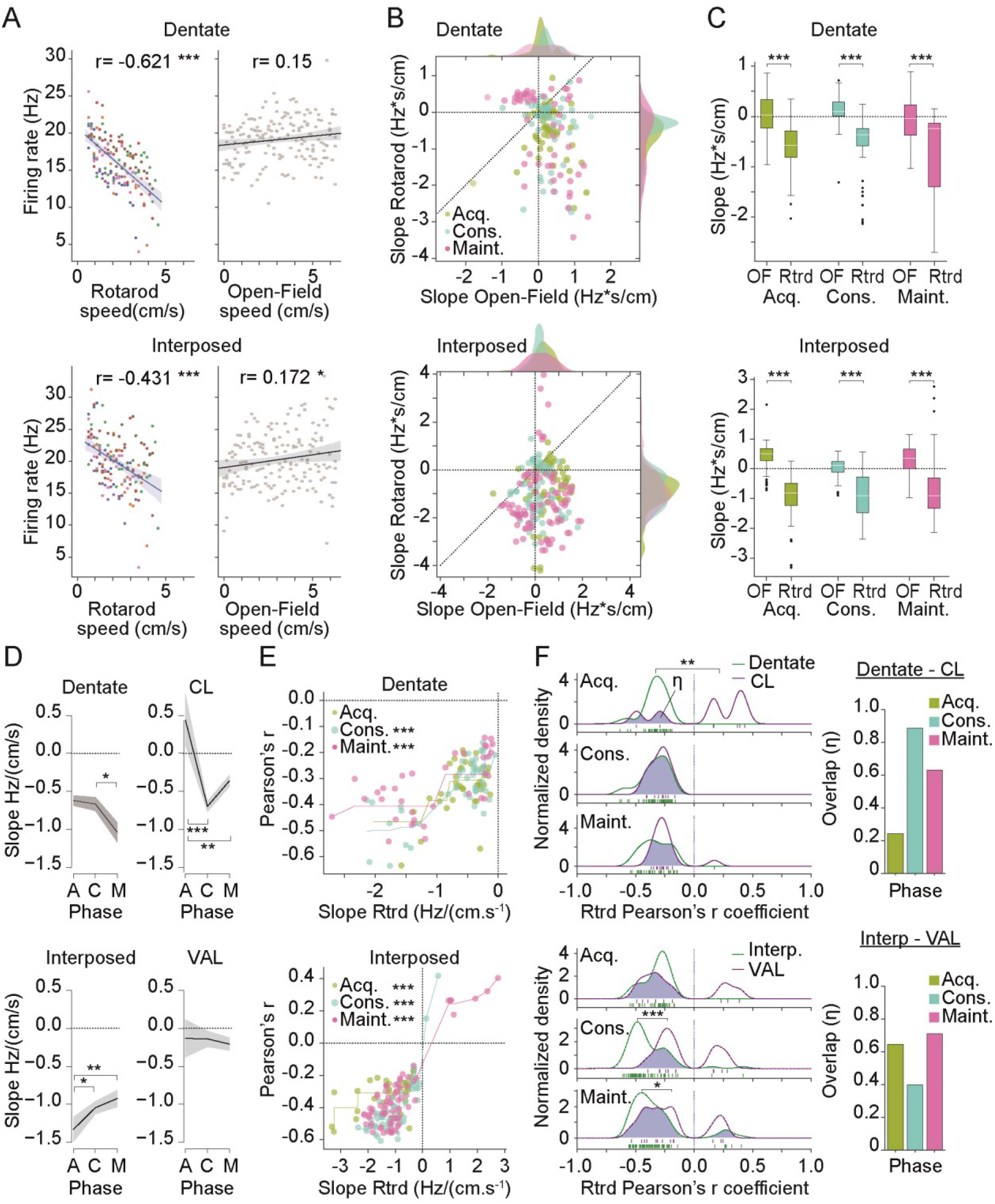
Dentate and Interposed nuclei display a context-dependent sensitivity to speed, with a negative correlation between cerebellar activity and rotarod speed. a) Scatterplot showing the average firing rate per speed bins during the different trials of rotarod (left) versus open-field (right) session, for Dentate (top) and Interposed (bottom) nucleus (linear regression lines are shown with a 95% confidence interval) (*p<0.05, ***p<0.001 Pearson correlation). b) Scatter plot showing the slope of linear regression explaining the firing rate by the speed, in the open-field versus on the rotarod for each neuron in Dentate (top) and Interposed (bottom) nucleus during Acquisition (Acq.), Consolidation (Cons.) and Maintenance (Maint.). The diagonal dotted line represents equality between the slope in the open-Field and rotarod. Marginal axes show the histograms of the distributions of slopes, smoothed using a Gaussian kernel density estimate (ς=0.05). c) Boxplots displaying the distributions of linear regression slopes of neurons from Dentate (top) and Interposed (bottom) during open-field (OF) and rotarod (Rtrd) (***p<0.001 Wilcoxon test). d) Significant linear regression slopes for rotarod (mean +/− SEM) for Dentate (top left), centrolateral thalamus (CL, top right), Interposed (bottom left) and ventral anterior lateral thalamus (VAL, bottom right) during Acquisition (A), Consolidation (C) and Maintenance (M) (*p<0.05, **p<0.01, ***p<0.001 Tukey Posthoc test). e) Scatter plot showing the correspondence between the slope of linear regression on rotarod versus the associated Pearson correlation coefficient for each neuron for Dentate (top) and Interposed (bottom) nucleus during Acquisition (Acq.), Consolidation (Cons.) and Maintenance (Maint.). The lines represent the isotonic regression of the Pearson’s r by the slope on rotarod (***p<0.001 Spearman Rank test). f) Density of Pearson’s r coefficients for Dentate and CL, (top) and Interposed and VAL, (bottom) for Acquisition, Consolidation and Maintenance phase, the overlapping area is represented in blue and is associated to the overlapping index η (*p<0.05, **p<0.01, ***p<0.001 Mann Whitney test); a barplot of the value of η is shown on the right.

Overall, these results indicate the presence of a number of cells in the cerebellar nuclei modulated during the rotarod task. This modulation consists mostly in a reduction in firing rate proportional to the locomotion speed on the rotarod, which is not observed in the open-field locomotion, suggesting a specific engagement of these cells in the task execution. Moreover, this modulation is observed in all phases of learning. In contrast, the modulation of units in the CL thalamus exhibit stronger changes in the course of learning, suggesting changes in the encoding of the task in the motor circuits; the observations in the VAL thalamus suggest an earlier engagement of this structure since the modulation was stable along the learning phases. In addition to these observations, we found that the cells in the Dentate and CL thalamus exhibited more similar modulations by the speed in late phases (Consolidation and Maintenance), while the similarity of modulation by speed in Interposed and VAL thalamus cells remained more stable across all phases. This suggests that Dentate-CL and Interposed-VAL pathways have a differential engagement during motor learning, the former exhibiting rather an increased engagement in the course of learning.

### Neuronal activity during rest is modified during and after learning

Motor skill learning undergoes consolidation between episodes of learning, and we therefore examined the activity of the motor circuits during periods of immobility before, between and after rotarod trials (Fig 3). For open-field sessions, a reduction of the mean firing rate was found in all phases of learning in the Interposed nucleus and in the Maintenance phase in the Dentate, while little if any change was observed in the thalamus (Fig 3A, Suppl. Table 9). Moreover, we observed that this reduction was more important during Consolidation and Maintenance for Interposed cells (Suppl. Table 10), whereas it remained stable during all the phases for Dentate cells (Suppl. Table 10). In a smaller set of animals, we recorded the firing rate in the open-field quiet periods and in the resting period between rotarod trials (Inter-trials). In these cases, we observed different patterns of discharge during rest (Fig 3B, Suppl. Table 11): in the cerebellar nuclei the firing rate during the quiet period between trials and after learning in the open-field decreased in the Consolidation and Maintenance phases compared to the firing in the open-field before the learning sessions of the day (Suppl. Table 11). In the VAL thalamus the firing rate increased during the inter-trial resting periods and returned progressively to the baseline in the open-field after learning, while in CL thalamus this evolution was only observed in Acquisition and Maintenance phases (Suppl. Table 11). Overall, these results indicate the presence of lasting changes in the cerebellum “offline” firing during quiet periods during and following the learning session, while thalamic neurons only exhibited changes in firing at rest only at shorter timescale, between trials.

**Fig 3.**
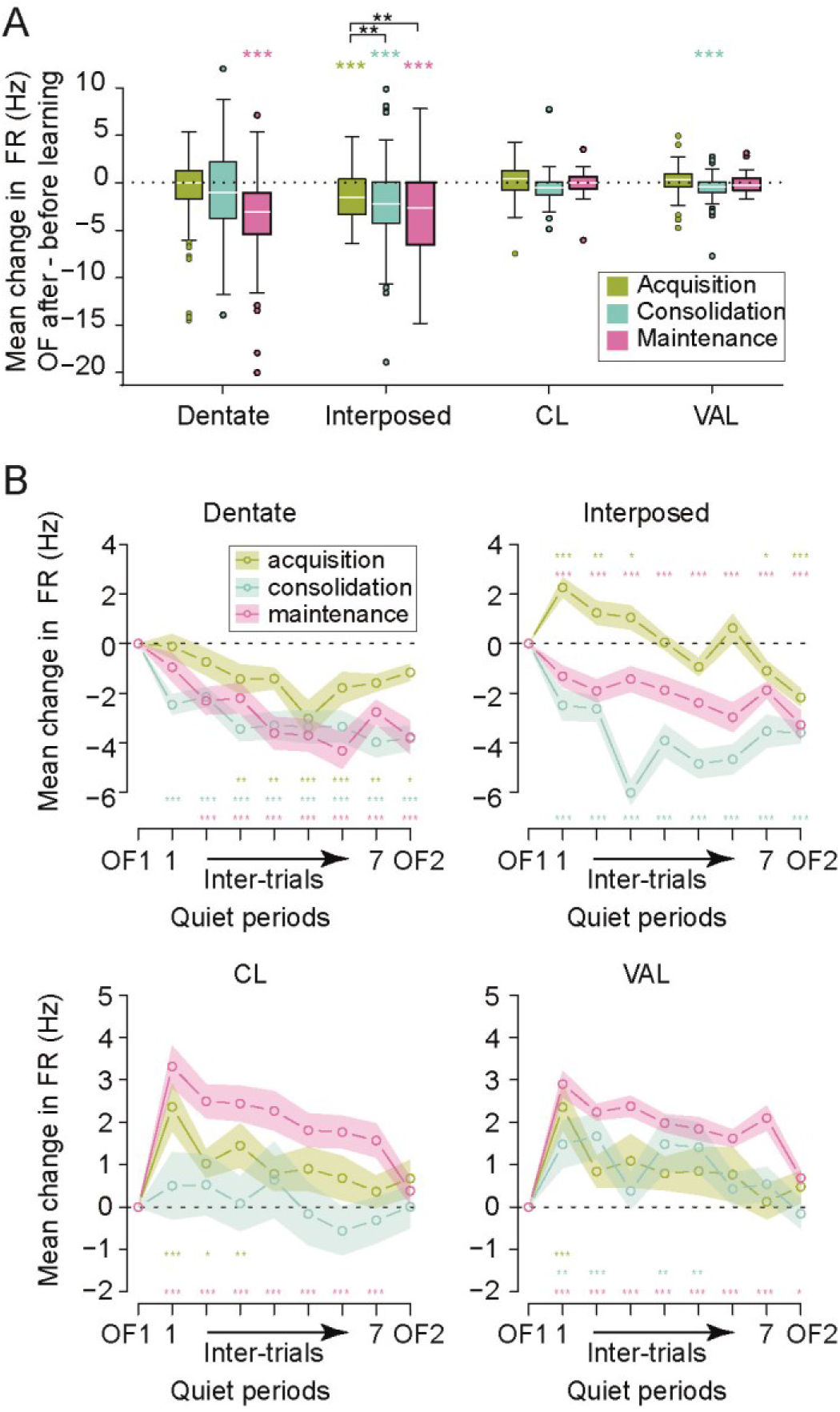
The quiet state activity in the cerebellar nuclei and in the thalamus changes during the learning protocol. a) Boxplots showing the distribution of change in mean firing rate between open-field (OF) before and after rotarod, during acquisition, consolidation and maintenance for Dentate, Interposed, centrolateral thalamus (CL) and ventral anterior lateral thalamus (VAL) (***p<0.001 Tukey Posthoc test). b) Evolution of the mean quiet state firing rate (mean +/− SEM) in open-field sessions and Inter-trial periods during Acquisition, Consolidation and Maintenance for Dentate (top left), Interposed (top right), CL (bottom left) and VAL (bottom right). Mean firing rate was normalized by subtracting the open-field before (OF1) (*p<0.05, **p<0.01, ***p<0.001 Dunett Posthoc test).

### Partial inhibition of the cerebellar nuclei using hM4D(Gi)-DREADD does not impact basic motor abilities

While the analysis above shows that many neurons exhibit some modulation of firing during and after the accelerated rotarod, the contribution of this activity to learning and behavior remains unknown. To evaluate this contribution, we used a chemogenetic approach (Roth, 2016) using the inhibitory DREADD (hM4D-Gi) activated by the synthetic drug Clozapine-N-Oxide (CNO).

In order to validate this approach, mice were injected with AAV5-hSyn-hM4D(Gi)-mCherry (DREADD) or AAV5-hSyn-EGFP (Sham) and implanted with microelectrode arrays in the cerebellar nuclei (Fig 4A-H). Post-hoc histology confirmed the position of the electrodes (Fig 4E) and showed that the expression of AAV5-hSyn-hM4D(Gi)-mCherry was confined to the cerebellar nuclei and restricted to the neuronal membrane, with 78% of the cells expressing hM4D(Gi)-DREADD in the cerebellar nuclei (Fig 4B-D). A week after surgery, the neuronal activity was recorded in the cerebellar nuclei in an open-field arena before and after accelerated rotarod sessions. Neither saline (SAL) in DREADD mice nor CNO injection (1 mg/kg) in Sham mice affected the firing rate, while the same dose of CNO induced a significant decrease in the firing rate of DREADD mice during Acquisition (day 1), Consolidation (day 4) and Maintenance (day 7) phases (Fig 4G and H, Suppl. Table 12).

**Fig 4.**
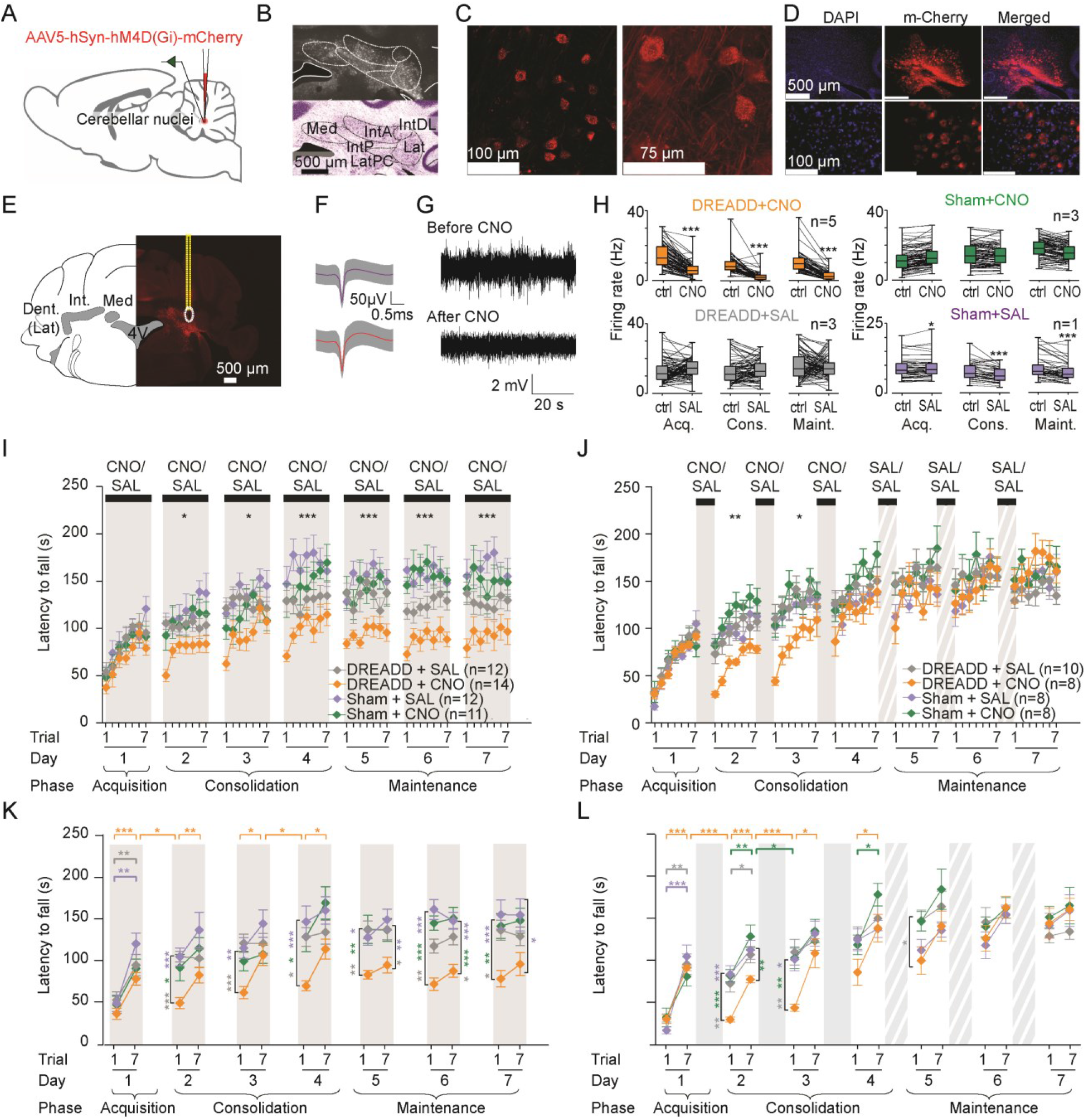
Cerebellar inactivation impairs the performance in the Consolidation and Maintenance but not Acquisition phases of motor learning task. a) Scheme showing electrode position and viruses injections. b) Coronal section of the cerebellum showing hM4Di-DREADD expression in the three cerebellar nuclei. c) Representative confocal image showing hM4Di-DREADD expression on neuronal membranes. d) Image showing DAPI positive neurons expressing hM4Di-DREADD. e) Coronal section of the cerebellum showing electrode placement (yellow dotted line)(left) and cells expressing hM4Di-DREADD nearby the electrode (right). f) Examples of spike shapes obtained from spike sorting in cerebellar nuclei (mean +/− SD). g) Examples of high-pass filtered traces of cerebellar nuclei recordings before and after CNO injection. h) Boxplots showing the mean firing rate of neurons recorded in DREADD and non-DREADD injected mice after CNO or SAL injection during Acquisition (Acq.), Consolidation (Cons.) and Maintenance (Maint). Cerebellar nuclei firing rate was reduced after 1 mg/kg of CNO injection in DREADD injected mice (***p<0.001 paired t-test). Small variations were observed in Sham+SAL group (*p<0.05, ***p<0.001 paired t-test). i) Daily of injections of CNO before trial 1 in DREADD injected mice induced impairment in motor learning during Consolidation and Maintenance but not Acquisition phase (*p<0.05, ***p<0.001; repeated measure ANOVA Group effect). j) CNO injections after Acquisition phase (30 min after trial 7) induced impairment in motor learning during Consolidation phase (*p<0.05, **p<0.01; repeated measure ANOVA Group effect). k,l) Latencies to fall in trial 1 and 7 were compared for each day (horizontal comparison lines trial 1 vs trial 7; *p<0.05, **p<0.01, ***p<0.001 paired t-test). Latencies to fall in the trial 7 of a day and trial 1 of the next day were compared (horizontal comparison lines trial 7 vs trial 1; *p<0.05, ***p<0.001 paired t-test). Comparisons between groups for trials 1 and trials 7 were also represented (vertical comparison lines, respectively trial 1 difference to controls and trial 7 difference to controls; *p<0.05, **p<0.01, ***p<0.001 t-test). Data represents mean ± S.E.M. *n* indicates the number of mice.

We then examined whether the reduction of cerebellar nuclei firing impacted on the spontaneous motor activity, strength and motor coordination. Analysis of the open-field locomotor activity revealed that velocity was not altered by CNO or SAL injection in DREADD or Sham mice (Fig Sup2A, Suppl. Table 26). In addition, no significant differences were observed between the experimental groups in the fixed speed rotarod (Fig Sup2B, Suppl. Table 27), footprint patterns (Fig Sup2C, Suppl. Table 28), grid test (Fig Sup2D, Suppl. Table 28), horizontal bar (Fig Sup2E, Suppl. Table 28) and vertical pole (Fig Sup2F, Suppl. Table 28) indicating that fatigue, coordination and strength are not affected by the reduction of cerebellar nuclei firing induced by 1mg/kg CNO. These results thus indicate that the reduction of the cerebellar nuclei firing induced by the CNO had a limited impact on the basic motor abilities of the mice.

### Cerebellar inactivation by hM4D(Gi)-DREADD during or after task impact on motor learning

To test the effect of a reduction of cerebellar output on motor learning, we first examined the impact of cerebellar nuclei inhibition by injecting CNO (1 mg/kg) each day before the first trial of an accelerated rotarod session (Fig 4I,K, Suppl. Table 13,14,15). This treatment did not affect significantly the learning on the first day, but reduced the performance on the following days compared to the control groups (Sham+SAL, Sham+CNO: Sham mice which received either SAL or CNO, and DREADD+SAL: DREADD mice which received SAL, Suppl. Table 13). During the Consolidation phase, the performance of the initial trial, but not the last ones, of each day was reduced (Suppl. Table 15), indicating that the DREADD+CNO animals could compensate within each day the low performance of the first trials. These poor performances were associated with a significant loss of performance between the last trial of one day and the following day (Suppl. Table 14), indicating a defect in the consolidation of motor learning in the DREADD+CNO mice. During the Maintenance phase, the performance of the DREADD+CNO group remained lower than the control groups both for the first and last trials of the task, and no further improvement of the performance of this group was observed during this phase (Suppl. Table 15). These results show that the inhibition of the cerebellar nuclei impacts on learning consolidation in the Consolidation phase, and on the execution of the task when reaching the Maintenance phase.

The action of CNO administration lasts longer than the learning trials; therefore the results above do not differentiate the action of cerebellar inhibition during and after the learning trials. We therefore injected another set of mice with CNO after the training sessions (Fig 4J,L, Suppl. Table 13,14,15). Cerebellar nuclei inhibition after learning in the Acquisition and Consolidation phases reduced the performance on the first trials of the next day, but did not prevent learning within days (Suppl. Table 14), indicating a defect in the offline consolidation, which could be overcome by training during the day. The performance on the last day of the Consolidation phase of DREADD+CNO mice was not different from the mice of the control groups (Suppl. Table 13). These results indicate that the cerebellum participates to the offline consolidation of the accelerated rotarod learning.

### Selective cerebello-thalamic pathways differentially impact motor learning

The cerebellar nuclei project to a wide array of targets; to examine whether cerebellar neurons with distinct targets differentially contribute to the rotarod learning and execution, an AAV5-hSyn-DIO-hM4D(Gi)-mCherry virus expressing an inhibitory DREADD conditioned to the presence of Cre-recombinase was injected into the Dentate and Interposed cerebellar nuclei, while a retrograde CAV-2 virus expressing the Cre recombinase was injected either in the CL or in the VAL, which respectively relay cerebellar activity to the striatum and cerebral cortex. We could then express the inhibitory DREADD either in the Dentate neurons projecting to the CL (Dentate-CL, Fig 5A) or in Interposed neurons projecting to the VAL thalamus (Interposed-VAL, Fig 5D). Since we did not observe an effect of CNO in Sham mice, we only compared DREADD-injected animals receiving either CNO or SAL.

**Fig 5.**
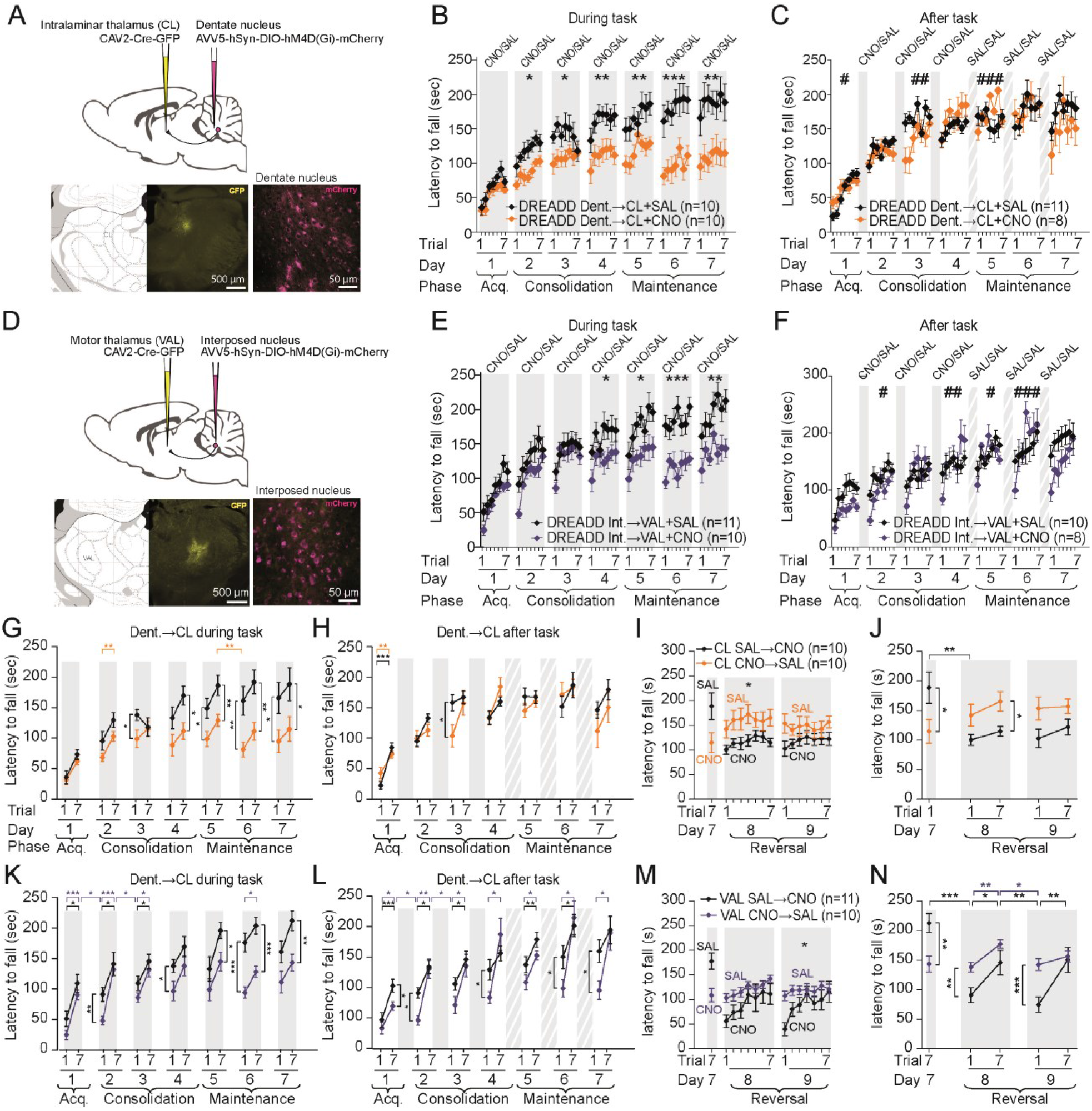
Dentate-centrolateral thalamus and Interposed-ventral anterior lateral thalamus pathway inactivation impairs consolidation and maintenance but not acquisition phase of motor learning task. Schemes showing AVV5-hSyn-DIO-hM4D(Gi)-mCherry and CAV2-Cre-GFP virus injections in Dentate and centrolateral thalamus (CL) (a) or Interposed and ventral anterior lateral thalamus (VAL) (d), respectively. Example of Dentate-CL (a, bottom) and Interposed-VAL (d, bottom) neurons expressing GFP (left, 500μm) and m-Cherry (right, 50 μm). b) Daily of injections of CNO before trial 1 in Dentate-CL pathway injected mice induced global impairments during Maintenance but not Acquisition and Consolidation phase (*p<0.05, **p<0.01, ***p<0.001; repeated measure ANOVA Group effect). c) CNO injections after Acquisition phase (30 minutes after trial 7) in Dentate-CL pathway did not induce alterations during Consolidation or Maintenance phase (#p<0.05, ##p<0.01, ###p<0.001; repeated measure ANOVA Group effect). e) Daily injections of CNO before trial 1 in Interposed-VAL pathway injected mice induced early impairments during Consolidation and Maintenance but not Acquisition phase (*p<0.05, **p<0.01, ***p<0.001; repeated measure ANOVA Group effect). f) CNO injections after Acquisition phase (30 minutes after trial 7) in Interposed-VAL pathway induced early impairments during Consolidation and Maintenance phase (#p<0.05, ##p<0.01, ###p<0.001; repeated measure ANOVA Group effect). g,h,k,l) Latencies to fall in trial 1 and 7 were compared for each day (horizontal comparison lines trial 1 vs trial 7; *p<0.05, **p<0.01, ***p<0.001 paired t-test). Latencies to fall in the trial 7 of a day and trial 1 of the next day were compared (horizontal comparison lines trial 7 vs trial 1; *p<0.05, ***p<0.001 paired t-test). Comparisons between groups for trials 1 and trials 7 were also represented (vertical comparison lines, difference with controls trial 1 and trial 7; *p<0.05, **p<0.01, ***p<0.001 t-test). i,m) Dentate-CL and Interposed-VAL pathway inhibition during Maintenance after a proper Consolidation phase of motor learning task showed early impairments (*p<0.05; repeated measure ANOVA Group effect) (*p<0.05, **p<0.01 and ***p<0.001). j,n) Latencies to fall in trial 1 and 7 were compared for each day (horizontal comparison lines difference trial 1 vs next trial 7; *p<0.05, **p<0.01, ***p<0.001 paired t-test). Latencies to fall in the trial 7 of a day and trial 1 of the next day were compared (horizontal comparison lines trial 7 vs next trial 1; *p<0.05, ***p<0.001 paired t-test). Comparisons between groups for trials 1 and trials 7 were also represented (vertical comparison lines, difference from control, trial 1 and trial 7; *p<0.05, **p<0.01, ***p<0.001 t-test). Dent. : Dentate cerebellar nucleus, Int.: interposed cerebellar nucleus. Data represents mean ± S.E.M, *n* indicates the number of mice.

To examine the impact of the inhibition of spontaneous locomotion, motor coordination and strength, we first examined locomotor activity in open-field experiments (Fig Sup3A, Suppl. Table 29). Analysis of the open-field locomotor activity revealed that velocity was generally not systematically by CNO or SAL injection in DREADD or Sham mice (Fig Sup3A, Suppl. Table 29); significant differences in the first open-field (OF1) for Interposed-VAL+CNO and Dentate-CL+CNO indicating respectively slightly higher and lower speed in day 1 and 4, compared to control groups. In addition, no significant differences were observed between Dentate-CL+CNO in the fixed speed rotarod test (Fig Sup3B, Suppl. Table 30). We found a decrease in the latency to fall for 15 and 25 r.p.m. in the Interposed-VAL+CNO group. No significant differences were observed between the experimental groups for footprint patterns (Fig Sup3C, Suppl. Table 31), grid test (Fig Sup3D, Suppl. Table 31), horizontal bar (Fig Sup3E, Suppl. Table 31) and vertical pole (Fig Sup3F, Suppl. Table 31) indicating that coordination and strength are not affected by the inhibition of cerebellar-thalamic pathways induced by 1mg/kg CNO.

We then examined how the rotarod learning was impaired by inhibiting the cerebello-thalamic pathways during and after the task (Fig 5A-N), as for full cerebellar nuclei inhibition (see above). Inhibition of the Dentate-CL pathway during the task (Fig 5B,G, Suppl. Table 16,17,18) produced a progressive departure from the performance of the control group, yielding to a strong reduction of performance in the Maintenance phase. In contrast, when the inhibition took place after the task (Fig 5C,H, Suppl. Table 16,17,18), the performances remained similar to the control group (although incidentally, significant Group×Trial interaction or posthoc difference between trial 1 was observed on some days).

In contrast, the pattern induced by the inhibition of the Interposed-VAL pathway differed sensibly. First, when inhibition took place during the task (Fig 5E,K, Suppl. Table 16,17,18), there was a strong loss of performance from the last trial of one day to the first trial of the following day. This overnight loss was compensated by a fast relearning up to similar levels as the control mice during the Consolidation phase. In the Maintenance phase, the learning saturated at lower level than control mice but similar to the Maintenance phase of the Dentate-CL group. Second, when inhibition took place after the task (Fig 5F,L, Suppl. Table 16,17,18), a similar overnight decrease of performance was observed during the Consolidation phase. These results are consistent with a contribution of the offline activity (as observed for full cerebellar nuclei inhibition, Fig 4L Suppl. Table 14,15) to the overnight consolidation of the motor learning.

We then shifted the treatment of all mice to SAL in the Maintenance phase (Fig 5I,J,M,N, Suppl. Table 19,20,21). Interestingly, the mice previously treated with CNO continued to exhibit strongly reduced performances on the first trial of each day, but the performances then raised to the control level within each day (associated with significant within-day learning during the Maintenance phase, Fig 5I,M, Suppl. Table 19). This indicates that the mice still failed to consolidate new learning in the Maintenance phase, even in the absence of cerebellar inhibition.

Mice which learned the task while either Dentate-CL or Interposed-VAL neurons were inhibited both showed reduced performances compared to controls. To examine to which extent the reduced performance were associated to a deficit of execution versus a reduced learning, we switched the treatment between the two groups (Fig 5I,J,M,N, Suppl. Table 19,20,21): mice that received SAL during 7 previous days were then administered with CNO and *vice versa*. In both groups, mice that received CNO during learning did not exhibit a sudden improvement of performance upon replacement by SAL. In the case of the Dentate-CL (Fig 5I,J, Suppl. Table 19,20,21), the performances only a mildly improved during the two days of reverse treatment. In the case of Interposed-VAL (Fig 5M,N, Suppl. Table 19,20,21), the mice continued to show a pattern of overnight loss of performance and relearning, which yielded similar final values at the end of each day as under CNO. Reciprocally, in both groups, mice that learned under SAL and received CNO exhibited a sudden drop in performance in the first day of reverse treatment. In the case of the Dentate-CL (Fig 5I,J), no improvement within days was then observed, while the interposed-VAL (Fig 5M,N, Suppl. Table 19,20,21) mice exhibited the same alternation of within-day increases and overnight decrease of performances, as observed in the CNO-treated interposed-VAL mice before the reversal of treatment. These results show that the impact of cerebellar-thalamic inhibition during learning cannot be readily reversed by alleviating the inhibition. Cerebellar-thalamic inhibition after learning reduces the performance and prevents re-learning, but with a different pattern in the two pathways, the Interposed-VAL showing within-day improvement and overnight decline in performances absent in the Dentate-CL pathway.

### Learning is inversely correlated to initial performance in control animals

The measure of performance reflects the sum of the learning over all past sessions. To gain more insight into the nature of impairments induced by cerebellar inhibitions, we then reanalyzed the data by focusing on the change of performance across days and nights during the learning (Fig 6). The latency to fall of a single trial is a “noisy” measure of the degree of learning; to obtain a more reliable measure of learning, we simplified the data by performing a linear regression on the performance of each day where the slope indicates the learning rate of the day, and the endpoints provide estimates of the initial and final performances on each day (Fig 6A). An initial inspection of average change in latency during days (Fig 6B, warm colors, Suppl. Table 22,23) or change overnight (Fig 6B, cold colors, Suppl. Table 22,23) showed various patterns across the groups (all control groups were pooled here). Interestingly, within-day learning was still significant in the Maintenance phase –when performance does not increase anymore– but was counterbalanced by similar overnight loss of performance. In most cases, the amplitudes of within-day learning and overnight loss were not significantly different from the respective control groups, preventing thus to interpret the nature of difference in learning between groups. We then examined how the performance changes at each phase of learning evolved as a function of another.

**Fig 6.**
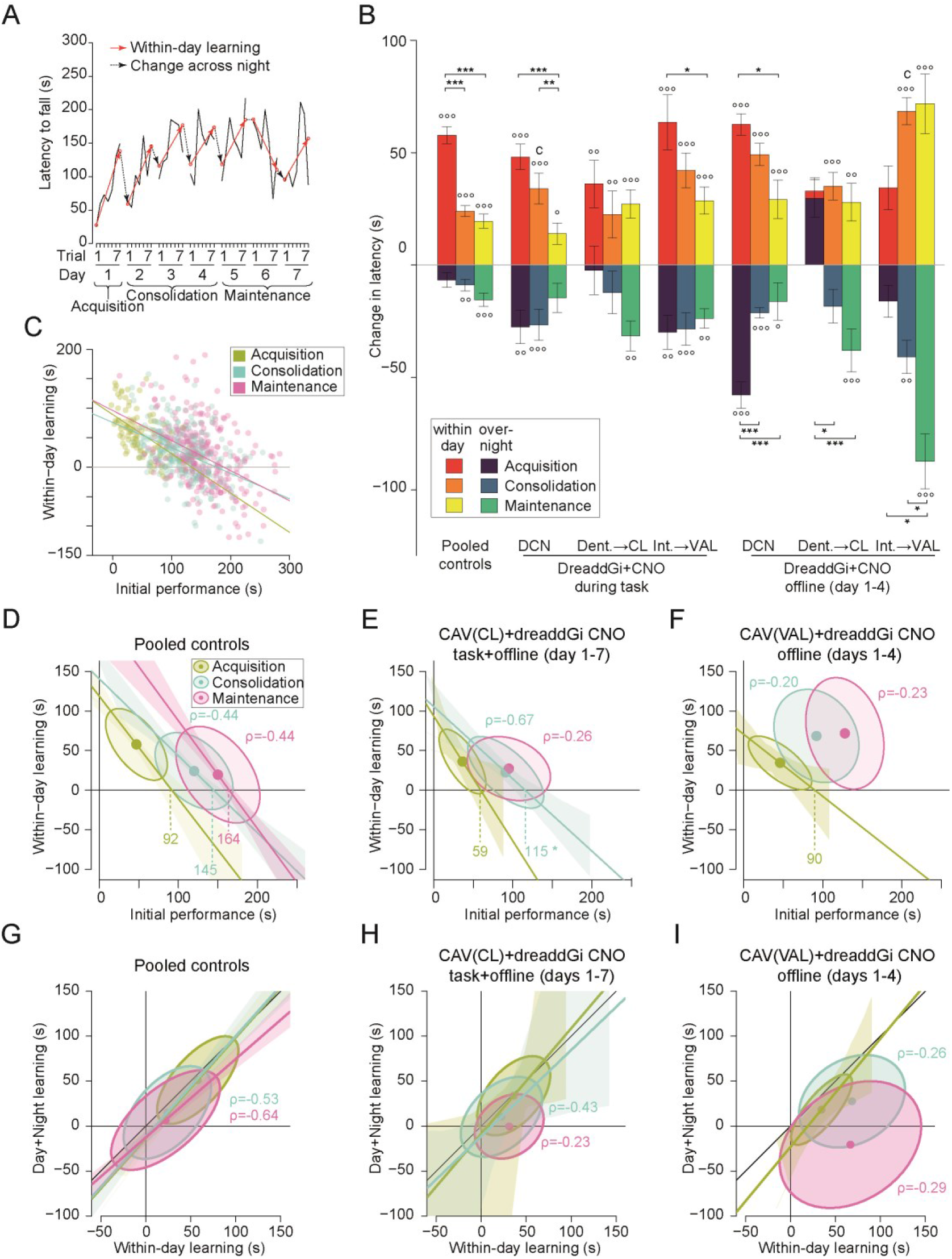
Learning profile is altered by the chemogenetic modulation of Dentate-centrolateral thalamus and Interposed-ventral anterior lateral thalamus pathway. a) Example of evolution of latencies to fall for a control mouse during the accelerated rotarod protocol, showing a high variability along session. Linear regressions estimated values of trial 1 and 7 during each day are shown using red hollow dots, within-day learning (red arrows) and overnight change (dotted black arrows). b) Bar plot showing change in latency associated to within-day learning and overnight change for all experimental groups in all phases (mean +/− SEM). Difference from 0: ° p<0.05, °° p<0.01 °°° p<0.001; Tukeys’ difference between phases: * p<0.05, ** p<0.01, *** p<0.001. Difference from control group: *c* p<0.05. c) Scatterplots showing the within-day learning as a function of the initial performances in pooled control group in all phases. Ordinary least square linear regressions outcomes are shown for each phase. Bivariate plot showing the within-day learning as a function of the initial performance (d,e,f) and day+night learning as a function of within-day learning (g,h,i) for Pooled controls (left), Dent.-CL task+offline (middle) and Int.-VAL offline (right). The ellipse contains 50% of a bivariate normal distribution fitted to the points and the dot indicates the center of the distribution. Deming linear regression outcomes are represented for each phase (d,e,f,g,h,i). The intercepts on the initial performance axis are represented for each phase (D,E,F). ρ indicates the value of the Pearson’s coefficient for each phase.

In the control group, we unexpectedly found an inverse correlation between the initial performance of each day and the amount of learning: animals with strong initial performance on a given day showed weaker improvements than animals with poor initial performances; this was true for all phases of learning (Fig 6C). The relationship between initial performance and within-day learning evolved across the phases (Fig 6C,D Suppl. Table 24). Notably, the intercept of the linear regression line on the x-axis revealed the existence of “fixed points”, which correspond to the initial performance which is not expected to be followed by an increase or decrease of performance following the training of the day. For the control group, such fixed points were found for the 3 phases of learning and exhibited a progressive increase in value from the Acquisition to the Maintenance phases (92s, 145s and 164s for the three phases, see Suppl. Table 24). Thus, in control mice, the amount of learning on each day depends on the initial performance of the day and is a decreasing function of the initial performance up to a fixed point where no learning is expected, which varies with the phases of learning.

The learning not only depends on the process taking place during the multiple trials of a day, but also on the ability of the animal to maintain the same level of performance the next day by consolidating the learning. We then examined how the increase of initial performance between two successive days was correlated with the gain of performance within the first of these two days. We found that these values were strongly correlated in the control group with a slope close to 1 at all phases of learning (Fig 6G, Suppl. Table 25); interestingly, the correlation remained high in the Maintenance phase even if the average performance of the population did not increase. This indicates that for each mouse, the improvement of performance during the task (or loss in the Maintenance phase) is fully retrieved the next day (Fig 7A top left).

**Fig 7.**
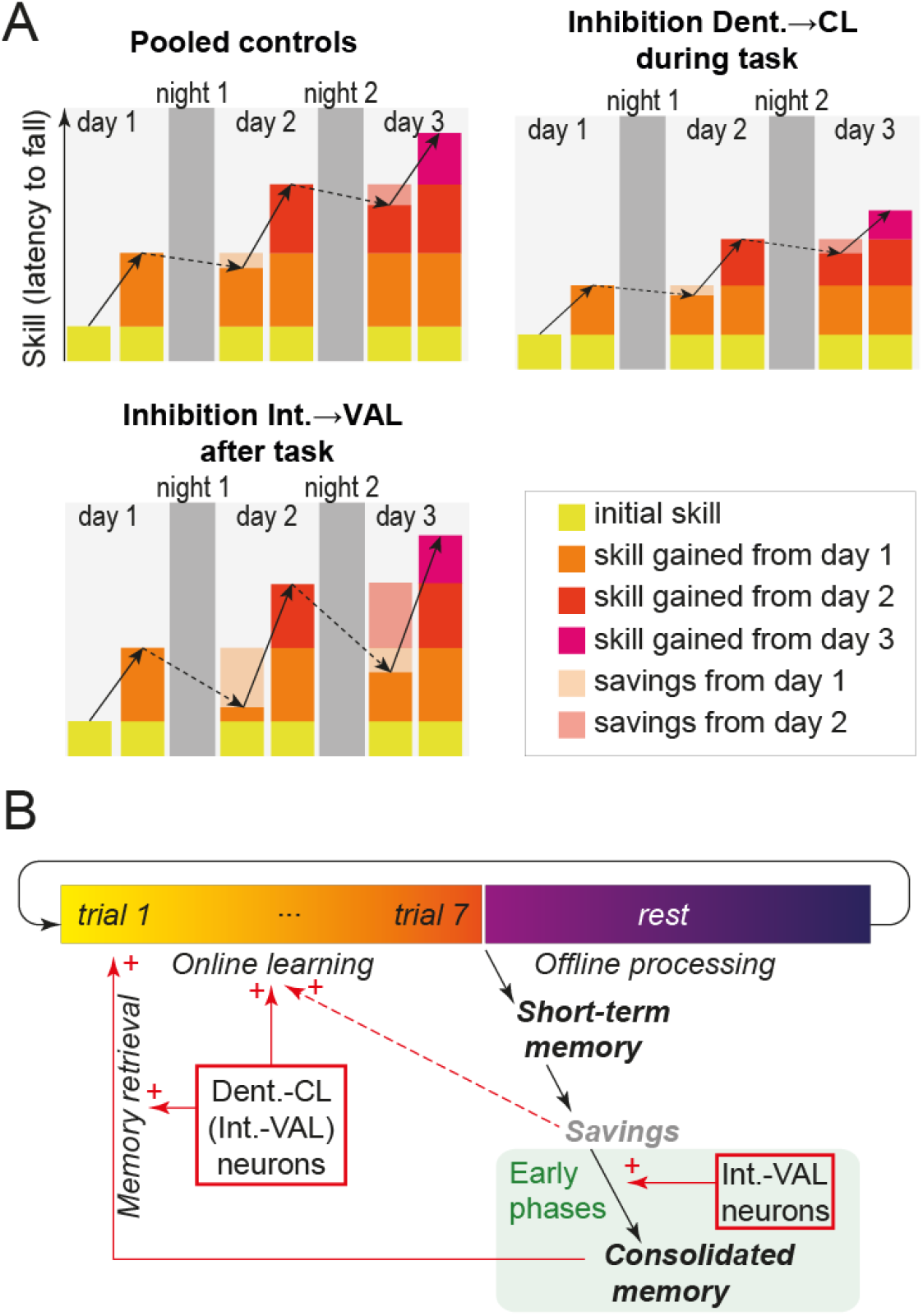
Summary of the behavioral findings. a) Schematic representation of the within-day learning (plain arrow) and overnight loss (dotted arrow) for Pooled controls, Dentate-centrolateral thalamus+CNO administered during the task (Dent.-CL+CNO) and Interposed-ventral anterior lateral thalamus+CNO administred after the task(Int.-VAL+CNO). In the former group, the intensity of learning is reduced but consolidation is intact, while in the latter, the consolidation is reduced but a latent trace remains in the form of savings that are promptly relearned the next day. b) Summary of the controls exerted by Dentate-CL and Interposed-VAL neurons on the skill learning: the Dentate-CL neurons contribute to online-learning and retrieval from learned skill, while the Interposed-VAL neurons contribute to consolidate offline the recent learning into a form of consolidated, readily-available, memory; in the absence of such consolidation, a trace of learning still remains in the form of savings, which will accelerate the online learning the next day; a defect in consolidation in the earlier phases cannot be rescued in the late phase.

In groups with full cerebellar nuclei inhibition, the learning patterns shall result from the combination of multiple output pathways, so we focused our attention on groups with discrete phenotypes: 1) the Dentate-CL group with inhibition by CNO during the task which shows reduced learning (Fig 5B,G) that contrasts with the lack of effect when inhibition by CNO takes place after the task (Fig 5C,H) indicating that the main effect occurs during learning; 2) the Interposed-VAL group with inhibition by CNO after the task which shows an effect on overnight change in performance (Fig 5F,L). We did not try to further analyze the effect of CNO inhibition during the task of Interposed-VAL mice, since their performance likely result from a compound effect of online inhibition –which resemble the effect found in Dentate-CL mice, but is difficult to interpret because of the reduced performance in the fixed-speed rotarod–, and of offline inhibition because the action of CNO extends beyond the duration of the task.

### Differential impact on learning of Dentate-CL and Interposed-VAL neurons inhibition

The inhibition of the Dentate-CL neurons during the task preserved the inverse correlation between the initial performance and the learning intensity in the Acquisition and Consolidation phases, but the intensity of learning was lower as attested by the lower values of the fixed points (59 s and 115 s respectively, Fig 6E, Suppl. Table 24). This indicates that for a given initial performance, the amount of learning was lower in the CNO-treated group than in the corresponding SAL-treated control group (with similar slopes but non-overlapping confidence intervals for intercept of x axis in CNO vs SAL groups in Consolidation phase, Suppl. Table 24). In the Maintenance phase, the initial performances remained at a similar value as in the Consolidation phase, indicating that no further long-lasting learning occurred. In contrast with the control group, the correlation between initial performance and within-day learning was not significant anymore (Fig 6E, Suppl. Table 24). Moreover, while the learning was preserved overnight in the Acquisition and Consolidation phases, the correlation between within-day and between-days learning was lost in the Maintenance phase (Fig 6H, Suppl. Table 25). Thus, the inhibition of the Dentate-CL reduced the intensity of learning and prevented the consolidation in the late stage of learning (Fig 7A, top right).

The inhibition of the Interposed-VAL neurons after the task in the Acquisition and Consolidation phases was associated with a striking pattern of increased intensity of within-day learning and increased overnight forgetting in the Consolidation and Maintenance phases (Fig 6B, Suppl. Table 22). However, this came with a de-correlation of initial performance of each day and learning intensity (Fig 6F, Suppl. Table 24), except in the Acquisition phase, when the CNO has not yet been administered. Despite the large within-day learning, the initial performance of each day only marginally improved consistent with a limited consolidation. Indeed, the amount of learning over each day + following night was not correlated anymore with the learning during the day (Fig 6I, Suppl. Table 25) in the Consolidation and in the Maintenance phase. Interestingly, the increase in learning and reduced day+night learning observed in the Consolidation persisted in the Maintenance phase, once the CNO has been replaced by SAL, indicating that the deficit in learning could not be recovered afterwards (Fig 5F,L). These results suggest that despite a poor consolidation, the mice increased their performance on each day by regaining the learning from previous days and adding some more learning on top, a feature distinctive of the existence of savings (Fig 7A bottom left, see Discussion).

Overall these results indicate that the Dentate-CL and Interposed-VAL cerebellar neurons are involved in different functions and that the cerebellum controls multiple processes during and after learning sessions of accelerated rotarod.

## DISCUSSION

In this study, we observed task-specific discharge in the cerebellar nuclei consistent with an involvement of the intermediate and lateral parts of the cerebellum. We also found that the transient, mild chemogenetic modulation which reduces but does not suppress the cerebellar nuclei activity, preserves the motor abilities but disrupts motor learning. Moreover, we could distinguish two contributions of the cerebellum to learning; one is carried by neurons projecting toward the midline thalamus and possibly to the motor thalamus, and is needed for learning and recall. The other is carried by neurons projecting toward the motor thalamus and is required to perform an offline consolidation of a latent memory trace into a consolidated, readily available, motor skill (Fig 7). Thus, our results suggest that learning a complex motor task involves the coupling of the cerebellum with the basal ganglia, which is needed during the learning and execution, and the coupling of the cerebellum with the cortex which plays a critical role in the offline consolidation of the task.

### A role for the cerebellum in a multi-nodal network

The accelerating rotarod task has proven to be a powerful motor learning paradigm to study the multiple scales of the mechanisms of motor learning in the rodents ((e.g. Costa et al., 2004; Rothwell et al., 2014; Yang et al., 2009)). It is also suitable to study the multiple time scales of motor learning: when repeated over multiple days, distinct phases of learning, with different rate of performance improvement and organization of locomotion strategies, can be distinguished (Buitrago et al., 2004) and selectively disrupted (Hirata et al., 2016). The basal ganglia and motor cortex are recruited and required to complete the task (Cao et al., 2015; Costa et al., 2004; Kida et al., 2016). Moreover, the areas involved in the basal ganglia evolves along the phases of learning (Cao et al., 2015; Durieux et al., 2012; Yin et al., 2009). Consistent with the growing evidence of the dynamical interplay between the cerebellum, cortex and basal ganglia (Caligiore et al., 2017), the involvement of the connections between the cerebellum and the other structures in motor learning is thus expected.

The impact of cerebellar defects or manipulations on rotarod learning have been examined in too many studies to be listed exhaustively here, but the reported effects range from ataxia and disruption of the ability to run on a rod (Sausbier et al., 2004), to normal learning (Galliano et al., 2013), defects in learning (Groszer et al., 2008), defects in consolidation (Sano et al., 2018) and even increase in learning (Iscru et al., 2009). However, the cerebellum is critical for interlimb coordination (Machado et al., 2015), and many studies lack proper motor controls to test the ability to walk on a rotating rod: the improvement in performance may be limited by problems of running on the rod rather than problems of learning. Moreover studies often involve genetic mutations which leave room from variable compensations along development and adult life, as exemplified by the diversity of the motor phenotype of mice with degeneration of Purkinje cell (Porras-Garcia et al., 2013). Finally the multiphasic nature of rotarod learning is often overlooked. Yet, the targeted suppression of parallel-fiber to Purkinje cell synaptic long-term depression in the cerebellar cortex disrupts rotarod learning after the acquisition phase without altering any other motor ability (Galliano et al., 2013). Consistently, Thyrotropin-releasing hormone (TRH) knock-out mice do not express long-term depression at parallel fiber-Purkinje cell synapses and exhibit impaired performance in late phase of the rotarod learning, while the administration of TRH in the knock-out mice both restores long-term depression and accelerated rotarod learning(Watanave et al., 2018). More generally, studies in mutant mice suggest that cerebellar plasticity is required for adapting skilled locomotion(Vinueza Veloz et al., 2015). This raises the possibility that cerebellar plasticity is involved in accelerated rotarod learning.

### A specific impact on learning of DN-CL neurons

In our study, we found that the chemo-genetic inhibition of Dentate-CL neurons during the task reduces the performances of the mice in the late phases of learning. This effect unlikely results from basic motor deficits: we found that the chemogenetic modulation did not significantly alter 1) limb motor coordination in footprint analysis, 2) strength in the grid test, 3) speed in spontaneous locomotion in the open-field test, 4) locomotion speed and balance required to complete the horizontal bar test and 5) body-limb coordination and balance required in the vertical pole test. Since all these motor parameters may be affected by cerebellar lesions, this suggests that Dentate-CL neurons are not necessary to maintain those functions, which might thus be relayed by other cerebellar nuclei neurons; indeed focal lesions in the intermediate cerebellum (thus projecting to the Interposed nuclei) has been reported to induce ataxia without altering rotarod learning (Stroobants et al., 2013). Alternatively the effect of the partial inhibition induced by CNO (typically ~50-80% reduction in firing rate) may be compensated at other levels in the motor system to ensure normal performances in these tasks; indeed, the selective ablation of Dentate-CL neurons has been reported to yield locomotor deficits in the initial performances on the accelerated rotarod (Sakayori et al., 2019), which contrasts with the lack of significant deficit in the Acquisition phase following Dentate-CL (partial) inhibition in our study. A possible explanation could be that our intervention selectively disrupted the advanced patterns of locomotion only needed at the higher speeds of the rotarod. However, the highest speeds reached on the rotarod correspond to the average locomotion speed in the open-field. Moreover, in our conditions, the inhibition of the Dentate-CL neurons did not produce significant deficits in the fixed speed rotarod; CNO-treated animals ran in average for about two minutes at 20 r.p.m. while they fell in average after the same amount of time on the accelerating rotarod, corresponding to a rotarod speed below 20 r.p.m. at the time of the fall. This rules out a contribution of weakness or fatigue to the latency to fall in the accelerated rotarod, the CNO-treated animals being able to run on the fixed speed rotarod more than twice the distance, at a higher speed, than the distance they run on the accelerated rotarod before falling. Overall, this indicates that the inhibition of the Dentate-CL neurons selectively affects accelerated-rotarod learning and not the elementary motor abilities needed in the task.

We observed in control mice and in mice with inhibition of Dentate-CL neurons that the daily gain in latency depended from the initial latency of the day; for a given initial performance, CNO-treated animals in the Consolidation phase showed lower increase in latency to fall indicating weaker learning rate. This effect did not reach significance in the Acquisition phase, possibly because the DN-CL network is more engaged in later phases of learning, as suggested by our electrophysiological experiments. Moreover, the administration of CNO in animals that learned under Saline SAL induced a sudden drop in performance, revealing a deficit in the execution of the learned task. This indicates that the Dentate-CL neurons both contribute to learning and retrieval of motor skills.

The inhibition of Interposed-VAL neurons during the task also yield lower levels of performance in the Maintenance stage, suggesting that these neurons contribute also to learning and retrieval of motor skills, although the mild defect in fixed speed rotarod could indicate the presence of an execution problem. Interestingly, both Dentate and Interposed nuclei contain some neurons with collaterals in both thalamic structures(Aumann and Horne, 1996; Sakayori et al., 2019), suggesting that the effect on learning could be mediated in part by a combined action on the learning process of the striatum (via the CL thalamus) and cortex (via the VAL thalamus); however, consistent with (Sakayori et al., 2019), the manipulations of cerebellar neurons retrogradely targeted either from the CL or from the VAL produce different effects.

### Contribution of VAL-projecting cerebellar neurons to offline consolidation

While in the control mice, the final performance at the end of a session could be reproduced at the beginning of the next session, this link was lost when Interposed-VAL neurons were inhibited after the task (‘offline’), suggesting an impairment of consolidation. However, in this group of mice, the daily gain of performance increased across days instead of decreasing; this larger gain of performance reveals the presence of savings. Therefore, if the inhibition of Interposed-VAL neurons alters the consolidation, a latent trace of the learning remains and allows for a faster relearning on the next day.

The effect of CNO peaks in less than an hour and lasts for several hours afterwards (Alexander et al., 2009); therefore consolidation is substantially disrupted by altering the cerebellar activity in the few hours that follow the learning session. We found evidence for reduced tonic activity during rest in the open-field immediately after the learning sessions in the Interposed nucleus in all phases of learning; this was also observed during the rest between trials in a smaller subset of animals (in which the other cerebellar and thalamic nuclei also exhibited significant changes in firing rate). These results suggest that the consolidation starts readily when the animals are resting. This falls in line with a number of evidence indicating that cerebellar-dependent learning is consolidated by the passage of time, even in the awake state (Cohen et al., 2005; Doyon et al., 2009; Muellbacher et al., 2002; Nagai et al., 2017; Shadmehr and Holcomb, 1997), although very few studies in the Human have performed offline cerebellar stimulations (Samaei et al., 2017).

In the case of rotarod, it has been noted that sleep is not required for the overnight preservation of performance (Nagai et al., 2017); however sleep may still be required for the change of cortical (Cao et al., 2015; Li et al., 2017) or striatal neuronal substrate of the accelerated rotarod skill (Yin et al., 2009). Therefore, while the offline activity of cerebellar nuclei could be more specifically associated in converting savings into readily available skills, multiple processes of memory consolidation would co-exist and operate at different timescales.

The existence of multiple timescales for consolidation has already been described in Human physiology where the movements could be consolidated without sleep while consolidation of goals (Cohen et al., 2005) or sequences (Doyon et al., 2009) would require sleep. It is indeed difficult, as for most real-life skills, to classify the accelerating rotarod as a pure adaptive learning or sequence learning: on one hand, the shape of the rod and its rotation induce a change in the correspondence between steps and subsequent body posture and thus require some locomotor adaptation. On the other hand, the acceleration of the rod introduces sequential aspects: 1) the same step executed a few seconds or tens of seconds apart in a trial result in different consequences on body position, and 2) asymptotic performances involve to use successively multiple types of gait as the trials progress (Buitrago et al., 2004). Following offline inhibition of Interposed-VAL neurons, which aspect would be maintained and which would be lost? Faster relearning has been proposed to reflect an improved performance at selecting successful strategies (Morehead et al., 2015; Ruitenberg et al., 2018). An attractive possibility could be that novel sensory-motor correspondences encountered on the rotarod would remained learned, possibly leaving a memory trace within the cerebellum, but these elementary ‘strategies’ would not be properly temporally ordered into a sequence over the duration of a trial (2 minutes); the next day learning session would benefit from the existence of these fragments of skill, but learning would still be required to order them properly. A similar idea has indeed been proposed for the contribution of cerebellum to sequence learning (Spencer and Ivry, 2009). Alternatively, recent studies revealed mechanisms which could serve sequence learning (Khilkevich et al., 2018; Ohmae and Medina, 2015) and would be affected by the offline inhibition of Interposed-VAL neurons via their feedback collaterals to the cerebellar cortex (Houck and Person, 2015). However, our study does not allow us to conclude on the nature of savings remaining after the offline inhibition of Interposed-VAL neurons.

### An internal model in the output of the cerebellum?

Improving the rotarod performance requires the mice to match their locomotion speed to the accelerating speed of the rod. In the cerebellar nuclei, we observed that neurons exhibited substantial modulation as a function of speed on the rotarod, with a negative slope in the majority of cases, but this modulation was very different from the modulation of speed in the open-field, which rather exhibited a positive slope if any. This suggests that the purpose of the cerebellar activity in the rotarod is not to specify the locomotion speed. In mice, the speed of locomotion and gait are transmitted to the motoneurons by neurons in the midbrain locomotor region, which receives sparse inputs from the forebrain except from the basal ganglia, and little if any inputs from the cerebellar nuclei (Caggiano et al., 2018). Therefore, in the accelerated rotarod task, the profile of locomotion speed determined in a cerebello-cortico-basal ganglia network is more likely propagated to the midbrain locomotor region via the output nuclei of the basal ganglia (Rueda-Orozco and Robbe, 2015).

It is well established that a number of neurons in the Interposed nucleus exhibit a modulation along the stride (Armstrong and Edgley, 1984; Sarnaik and Raman, 2018), and a transient optogenetic manipulation of this activity results in an alteration of the gait (Sarnaik and Raman, 2018). Indeed, the optogenetic activation of genetically-defined Interposed nucleus neurons projecting mainly to the ventral lateral thalamus and the red nucleus produced higher strides (Low et al., 2018). Stride-related signals are also found in the Dentate nucleus, which is more clearly recruited in response to perturbations (Schwartz et al., 1987) or during skilled locomotion such as on a ladder (Marple-Horvat and Criado, 1999); this is consistent with the selective impact of lateral cerebellum lesion to obstacle stepping while preserving overground locomotion (Aoki et al., 2013). The changes of firing rate in the Dentate and Interposed units reported in our study thus likely reflect the integrated representation of the stride (Sauerbrei et al., 2015). An alternate –and compatible-hypothesis, is that the cerebellar output reflects the expected consequences of actions, representing an internal model of locomotion on the rod. In the rotarod, the faster the rod turns, the less individual steps produce the expected forward progress; the decrease of nuclear cell firing as the speed increase could indeed reflect the estimated “undershooting” of the outcome of the steps.

In conclusion, our results provide clear evidence for the existence of online contributions of the cerebello-thalamic pathways to the formation and retrieval of motor memories distributed in a cerebello-striato-cortical network. They also show a contribution to the offline consolidation of savings by a distinct pathway. Thus, our work highlights the importance of studying the contribution to learning of single nodes in the motor network from an integrated perspective (Caligiore et al., 2017; Krakauer et al., 2019).

---

**Fig Sup1.**
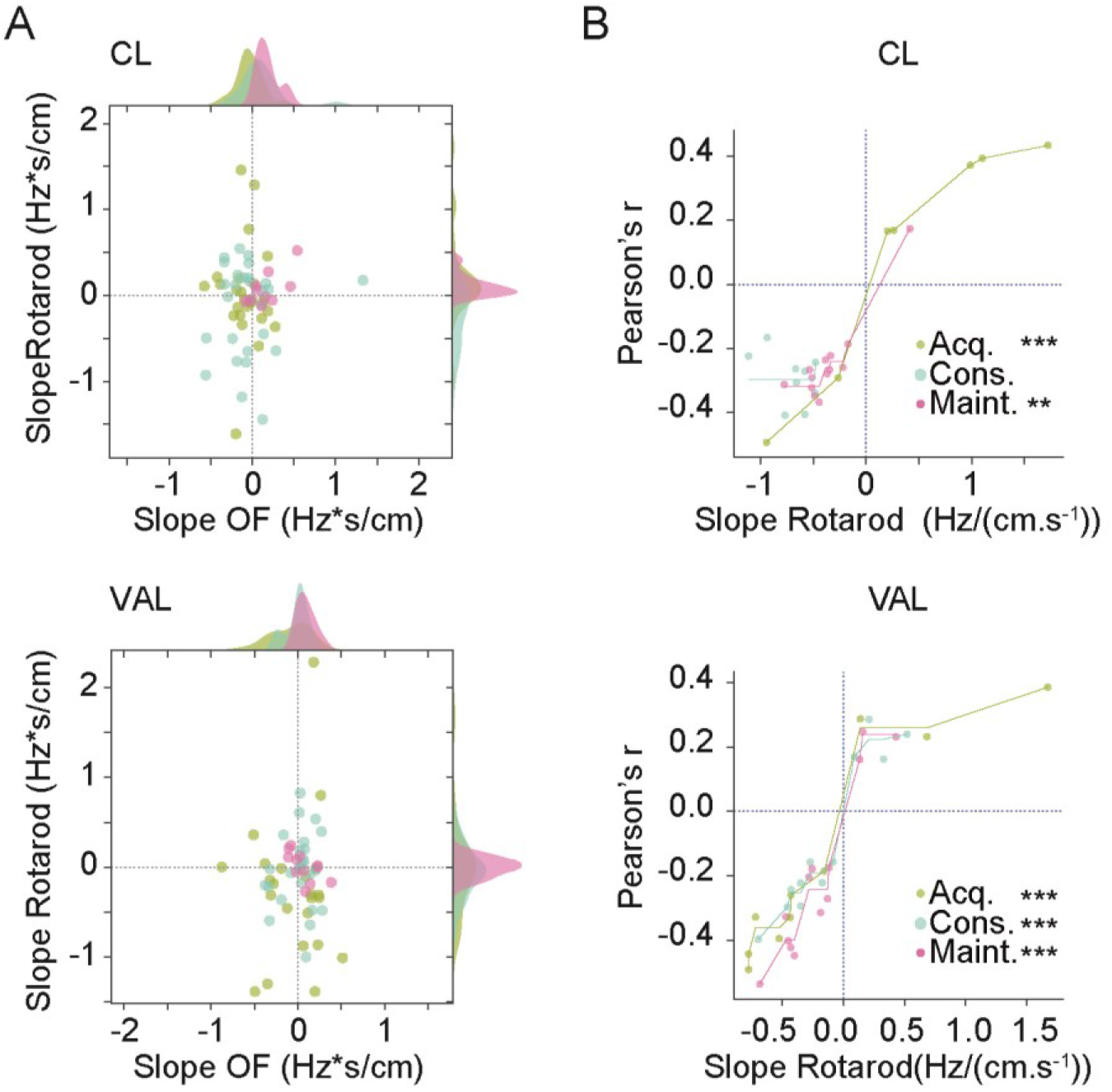
Centrolateral and ventral anterior lateral thalamus display a sensitivity to rotarod speed. a) Scatter plot showing the slope of linear regression explaining the firing rate by the speed, in the open-field versus the rotarod for each neuron in centrolateral thalamus (CL, top) and ventral anterior lateral thalamus (VAL, bottom) during Acquisition (Acq.), Consolidation (Cons.) and Maintenance (Maint.). Marginal axes show the histograms of the distributions of slopes, smoothed using a Gaussian kernel density estimate (ς=0.05). b) Scatter plot showing the correspondence of slope of linear regression on rotarod versus the associated Pearson correlation coefficient for each neuron for CL (top) and VAL (bottom), during Acquisition (Acq.), Consolidation (Cons.) and Maintenance (Maint.). The lines represent the isotonic regression of the Pearson’s r by the slope on rotarod (**p<0.01, ***p<0.001 Spearman Rank test).

**Fig Sup2.**
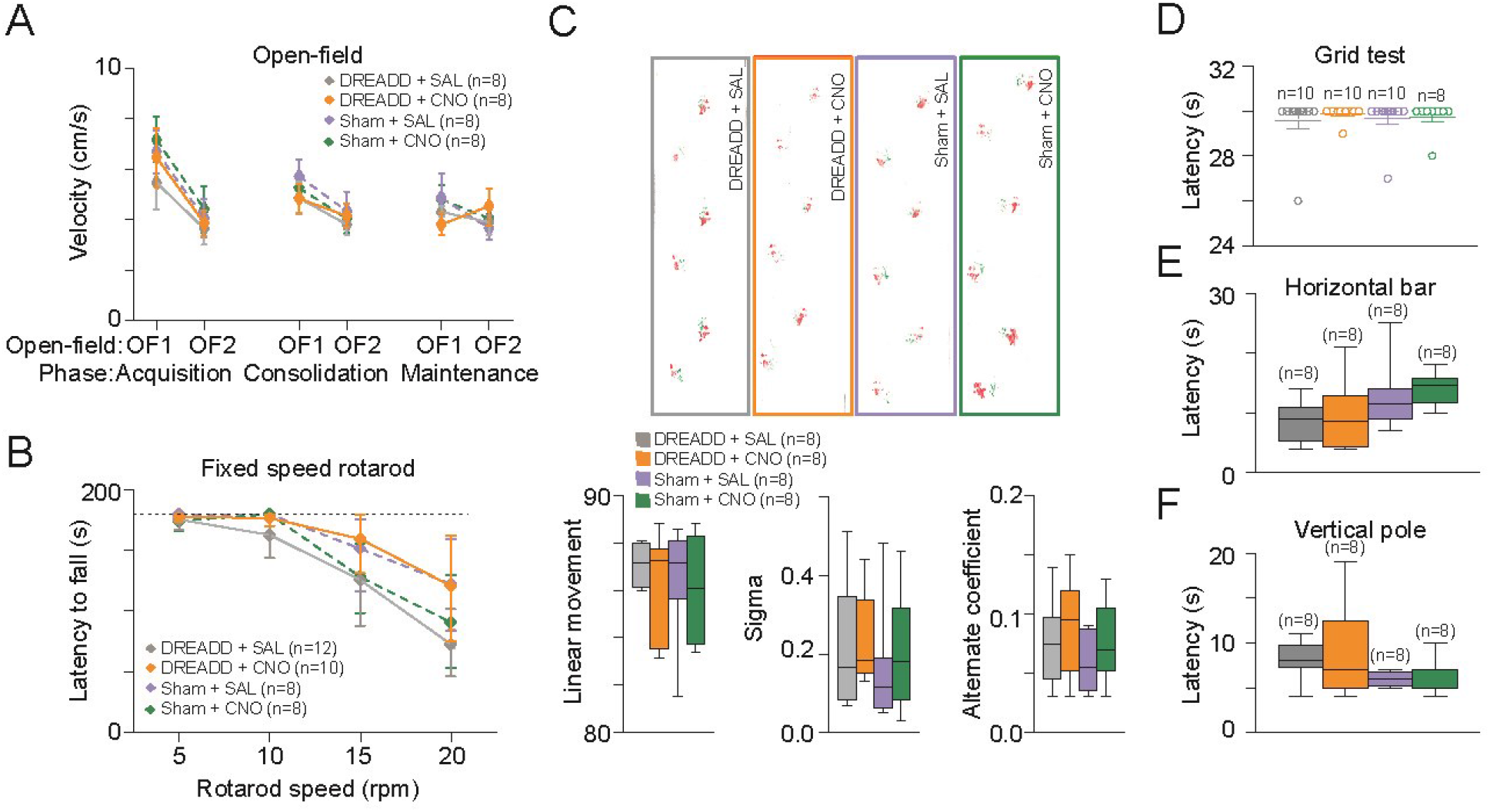
Cerebellar nuclei inhibition did not affect execution and fatigue, locomotion, motor coordination, balance and strength. a) Locomotor activity (Velocity) in DREADD and non-DREADD (Sham) injected mice after CNO or SAL injection during open-field sessions before (OF1) and after (OF2) rotarod for Acquisition, Consolidation and Maintenance (**p<0.01 t-test OF1 vs OF2). b) Latency to fall during fixed speed rotarod (5, 10, 15, 20, 25 r.p.m.) for all experimental groups. One way repeated measure ANOVA was performed on averaged values for all the speed steps in each experimental group followed by a Tukey Posthoc pairwise comparison. c) Footprint patterns were quantitatively assessed for 3 parameters as shown on representative footprint patterns (top) for all experimental groups. Three parameters are represented graphically: linear movement (bottom left), sigma (bottom middle) and alternation coefficient (bottom right). d) Latency reflecting the time before falling from the grid. 30 seconds of cut-off of was established as the maximum latency (dotted line on figure). e) Latency to cross the horizontal bar (balance beam test) for all experimental groups. f) Latency to reach home cage in vertical pole test for all experimental groups. *p<0.05 One Way ANOVA followed by a PostHoc Tukey test. *n* indicates the number of mice.

**Fig Sup3.**
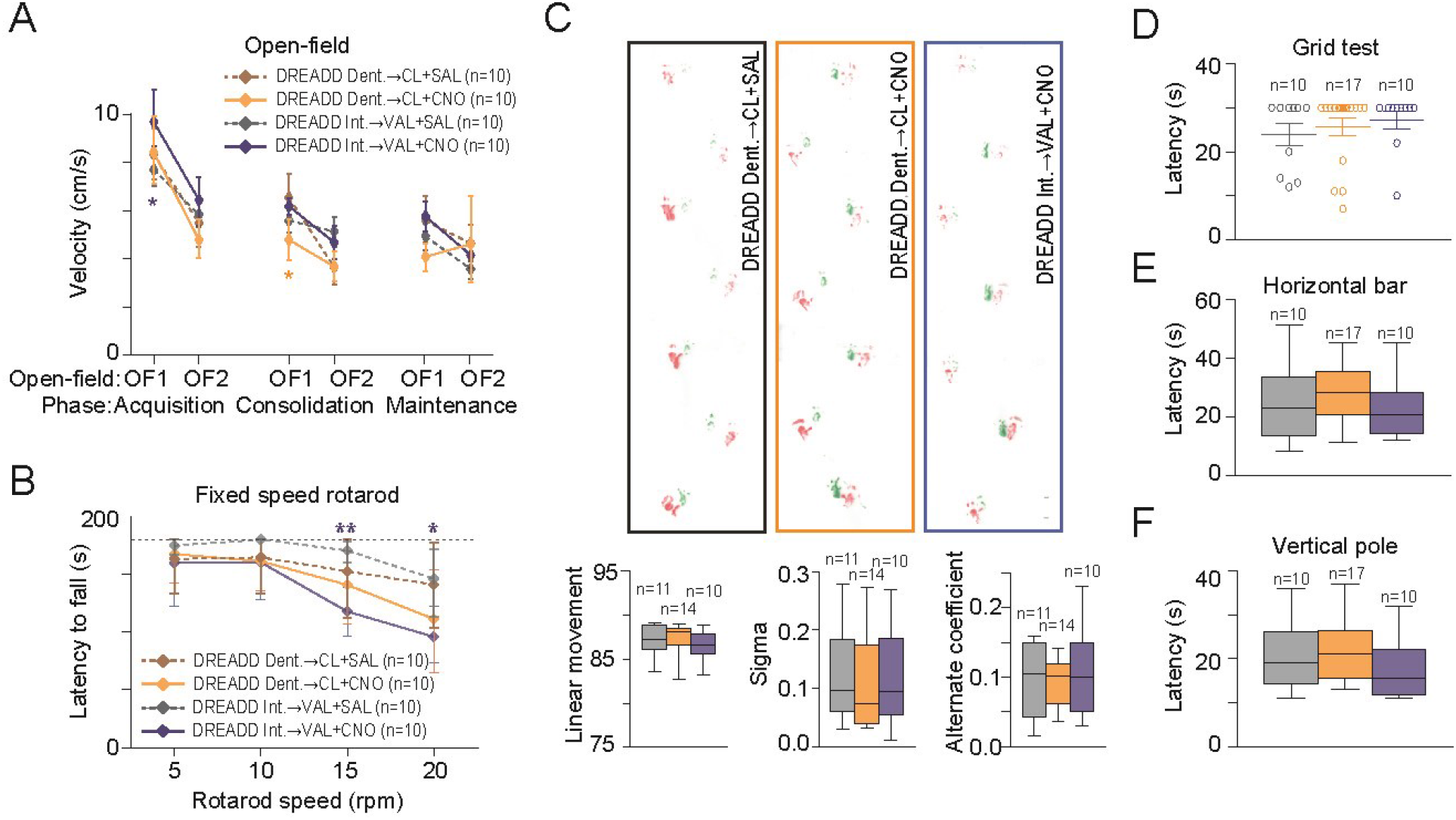
Dentate-centrolateral and Interposed-ventral anterior lateral inhibition did not affect execution and fatigue, locomotion, motor coordination, balance and strength. a) Locomotor activty (Velocity) in DREADD injected mice after CNO or SAL injection during open-fields sessions before (OF1) and after (OF2) rotarod for Acquisition, Consolidation and Maintenance (**p<0.01 t-test OF1 vs OF2). b) Latency to fall during fixed speed rotarod (5, 10, 15, 20 r.p.m.) for all experimental groups. One way repeated measure ANOVA was performed on averaged values for all the speed steps in each experimental group followed by a Tukey Posthoc pairwise comparison. c) Footprint patterns were quantitatively assessed for 3 parameters as shown on representative footprint patterns (top) for all experimental groups. Three parameters are represented graphically: linear movement (bottom left), sigma (bottom middle and alternation coefficient (bottom right). d) Latency reflecting the time before falling from the grid. 30 seconds of cut-off of was established as the maximum latency (dotted line on figure). e) Latency to cross the horizontal bar (balance beam test) for all experimental groups. f) Latency to reach home cage in vertical pole test for all experimental groups. *p<0.05 One Way ANOVA followed by a PostHoc Tukey test. DCN, deep cerebellar nuclei; CL, centrolateral thalamus; VAL, ventral anterior lateral thalamus. *n* indicates the number of mice.

## AUTHOR CONTRIBUTIONS

Conceptualization, A.P.V., D.P. and C.L.; Methodology, A.P.V. and D.P.; Software, R.W.S. and C.L.; Formal Analysis, A.P.V., R.W.S. and C.L.; Investigation, A.P.V., R.W.S., C.M.H and J.L.F.; Data Curation, A.P.V., R.W.S. and C.L.; Writing –Original Draft, A.P.V., C.L. and D.P.; Writing –Review & Editing, D.P. and C.L.; Visualization, A.P.V., R.W.S., J.L.F and C.L.; Supervision, D.P. and C.L.; Funding Acquisition, D.P. and C.L.

## ACKNOWLEDGMENTS

This work was supported by Fondation pour la Recherche Medicale (FRM, DPP20151033983) to D.P. and Agence Nationale de Recherche to D.P. (ANR-16-CE37-0003-02 Amedyst, ANR-19-CE37-0007-01 Multimod, Labex Memolife) and to C.L. (ANR-17-CE37-0009 Mopla, ANR-17-CE16-0019 Synpredict) and by the Institut National de la Santé et de la Recherche Médicale (France).

## STAR METHODS

### RESOURCE AVAILABILITY

#### Lead contact

Further information and requests for resources should be directed to and will be fulfilled by the Lead Contact, Daniela Popa.

#### Material availability

This study did not generate new unique reagents.

#### Data and Code availability

The data and source code generated during this study will be made available by the corresponding author upon a reasonable request.

### EXPERIMENTAL MODEL AND SUBJECT DETAILS

Adult male C57BL/6J mice (Charles River, France, IMSR Cat# JAX:000664, RRID:IMSR_JAX:000664), 8 weeks of age and 24 ± 0.4 g of weight at the beginning of the experiment were used in the study. Mice were fed with a chow diet and housed in a 22 °C animal facility with a 12-hr light/dark cycle (light phase 7am–7pm). The animals had free access to food and water. All animal procedures were performed in accordance with the recommendations contained in the European Community Council Directives.

### METHOD DETAILS

#### 1. Behavioral experiments

##### 1.1. Accelerated rotarod task

The rotarod apparatus (mouse rotarod, Ugo Basile) consisted of a plastic roller with small grooves running along its turning axis (Bearzatto et al., 2005). One week after injections, mice were trained with seven trials per day during seven consecutive days. This training protocol was chosen in order to distinguish the three different phases (Acquisition, Consolidation and Maintenance). During each trial, animals were placed on the rod rotating at a constant speed (4 r.p.m.), then the rod started to accelerate continuously from 4 to 40 r.p.m. over 300 s. The latency to fall off the rotarod was recorded. Animals that stayed on the rod for 300 s were removed from the rotarod and recorded as 300 s. Mice that clung to the rod for two complete revolutions were removed from the rod and time was recorded. Between each trial, mice were placed in their home cage for a 5-minutes interval.

##### 1.2. Open-field activity

Mice were placed in a circle arena made of plexiglas with 38 cm diameter and 15 cm height (Noldus, Netherlands) and video recorded from above. Each mouse was placed in the open-field for a period of 10 minutes before and after the accelerated rotarod task with the experimenter out of its view. The position of center of gravity of mice was tracked using an algorithm programmed in Python 3.5 and the OpenCV 4 library. Each frame obtained from the open fields videos were analyzed according to the following process: open-field area was selected and extracted in order to be transformed into a grayscale image. Then, a binary threshold was applied on this grayscale image to differentiate the mouse from the white background. To reduce the noise induced by the recording cable or by particles potentially present in the Open-field, a bilateral filter and a Gaussian blur were sequentially applied, since those components are supposed to have a higher spatial frequency compared to the mouse. Finally, the OpenCV implementation of Canny algorithm was applied to detect the contours of the mouse, the position of the mouse was computed as mouse’s center of mass. The trajectory of the center of mass were interpolated in x and y using scipy’s Univariate Spline function (with smoothing factor s=0.2 x length of the data), allowing the extraction of a smoothed trajectory of the mouse. The distance traveled by the mouse between two consecutive frames was calculated as the variation of position of the mouse multiplied by a scale factor, to allow the conversion from pixel unit to centimeters. The total distance traveled was obtained by summing the previously calculated distances over the course of the entire open-field session. The speed was computed as the variation of position of center points on two consecutive frames divided by the time between these frames (the inverse of the number of frames per seconds). This speed was then averaged by creating sliding windows of 1 second. After each session, fecal boli were removed and the floor was wiped clean with a damp cloth and dried after the passing of each mouse. Active and quiet state was determined by using a bi-threshold method in which two consecutive thresholded and filtered frames were subtracted one from each other. In order to have a proper recognition of both active and quiet state, lower and upper threshold of the changed pixels were arbitrarily set to 0.04% and 0.12%, respectively. The both thresholds were based on the video acquisition conditions (camera resolution, focal distance of the objective and distance from the objective) and the size of the animal detected. For each time point, every percentage of changed pixels below the lower or above the upper threshold was considered as quiet or active state, and percentages placed between both thresholds were considered as a continuity of the previous state.

##### 1.3. Horizontal bar

Motor coordination and balance were estimated with the balance beam test which consists of a linear horizontal bar extended between two supports (length: 90 cm, diameter: 1.5 cm, height: 40 cm from a padded surface). The mouse is placed in one of the sides of the bar and released when all four paws gripped it. The mouse must cross the bar from one side to other and latencies before falling are measured in a single trial session with a 3-minutes cut-off period.

##### 1.4. Vertical pole

Motor coordination was estimated with the vertical pole test. The vertical pole (51 cm in length and 1.5 cm in diameter) was wrapped with white masking tape to provide a firm grip. Mice were placed heads up near the top of the pole and released when all four paws gripped the pole. The bottom section of the pole was fixated to its home-cage with the bedding present but without littermates. When placed on the pole, animals naturally tilt downward and climb down the length of the pole to reach their home cage. The time taken before going down to the home-cage with all four paws was recorded. A 20 seconds habituation was performed before placing the mice at the top of the pole. The test was given in a single trial session with a 3-minutes cut-off period.

##### 1.5. Footprint patterns

Motor coordination was also evaluated by analysing gait patterns. Mouse footprints were used to estimate foot opening angle and hindbase width, which reflects the extent of muscle loosening. The mice crossed an illuminated alley, 70 cm in length, 8 cm in width, and 16 cm in height, before entering a dark box at the end. Their hindpaws were coated with nontoxic water-soluble ink and the alley floor was covered with sheets of white paper. To obtain clearly visible footprints, at least 3 trials were conducted. The footprints were then scanned and examined with the Dvrtk software (Jean-Luc Vonesch, IGBMC). The stride length was measured with hindbase width formed by the distance between the right and left hindpaws.

##### 1.6. Grid test

The grid test is performed to measure the strength of the animal. It consists of placing the animal on a grid which tilts from a horizontal position of 0° to 180°. The animal is registered by the side and the time it drops is measured. The time limit for this experiment is 30 seconds. In those cases where the mice climbed up to the top of grid, a maximum latency of 30 seconds was applied.

##### 1.7. Fixed speed rotarod

Motor coordination, postural stability and fatigue were estimated with the rotorod (mouse rotarod, Ugo Basile). Facing away from the experimenter׳s view, the mice placed on top of the plastic roller were tested at constant speeds (5, 10, 15 and and 20 r.p.m). Latencies before falling were measured for up to 3 minutes in a single trial session.

#### 2. Chronic in vivo extracellular recordings

Recordings were performed in awake behaving control mice during the open-field as well as the accelerated rotarod sessions in the day 1, 4 ad 7. Recordings and analysis were performed using an acquisition system with 32 channels (sampling rate 25kHz; Tucker Davis Technology System 3) as described in (de Solages et al., 2008; Popa et al., 2013). Cells activity in the Intralaminar thalamus (centrolateral, CL), motor thalamus (ventral anterior lateral, VAL) and cerebellar nuclei (Dentate and Interposed) was recorded by using bundles of electrodes consisting in nichrome wire (0.005 inches diameter, Kanthal RO-800) folded and twisted into six to eight electrode bundles. To protect these bundles and ensure a good electrodes placement, they were then placed through metal tubing (8-10mm length, 0.16-0.18mm inner diameter, Coopers Needle Works Limited, UK) attached to an electrode interface board (EIB-16 or EIB-32; Neuralynx) by Loctite universal glue. Different configurations were used in order to record simultaneously, CL (from bregma: AP −1.70 mm, ML ±0.75 mm, DV −3.0 mm), VAL (from bregma: AP −1.4 mm, ML ±1.0 mm, DV −3.5 mm) and/or the cerebellar nuclei (from Bregma: Interposed: −6.0 AP, +/−1.5 ML, −2.1 depth from dura; Dentate: −6.2 AP, +/−2.3 ML, −2.4 depth from dura). Microwires of each bundle were connected to the EIB with gold pins (Neuralynx). The entire EIB and its connections were secured in place by dental cement for protection purpose. Electrodes were cut to the desired length (extending 0.5mm below tube tip). The impedance of every bundle was measured and gold-plated electrochemically to lowered and set microwire’s impedance to 200– 500 kΩ. Mice were anesthetized with isoflurane and placed in the stereotaxic apparatus, then skull and dura were removed above CL, VAL and cerebellar nuclei recording site (see section 2.5. for a detailed description of the surgical procedure). Electrodes bundles were lowered into the brain, the ground was placed on the cerebellar cortex and the assembly was secured with dental cement. One week after the surgery, we started to record cellular activity in CL, VAL and cerebellar nuclei in freely moving mice placed in the open-field and the accelerated rotarod sessions. Mice were habituated to the recording cable for 2–3 d before starting the recording. A custom-made pulley system balanced the weight and torque of the wires during running and allowed the wires to accompany the mouse during the accelerated rotarod task. Signal was acquired by headstage and amplifier from TDT (RZ2, RV2, Tucker-Davis Technologies, USA) and analyzed with Matlab and Python 3.5. The spike sorting was performed with Matlab (Mathworks, Natick, MA, USA) scripts based on *k*-means clustering on PCA of the spike waveforms (Paz et al., 2006). At the end of experiments, the placement of the electrodes (CL, VAL and cerebellar nuclei) was verified. For each neuron, the spike times were used to compute the raster plots and firing rate histograms across trial and along phases. Mean firing rate was calculated for rotarod trials, resting periods and open-field sessions. The algorithm described in 2.2.3 was used to separate from the open-field sessions the active or quiet state in order to determine their corresponding mean firing rate. In order to assess the effect of the task on the neurons, a Mann-Whitney test (with α=0.01) was used to compare instantaneous firing rate during each trial *vs* the resting period before and after the trial. For each trial, two distributions were created: one corresponding to the firing rates during the given trial, and the other to the firing rate during the resting period before and after the trial. In the case of trial 1, only the resting period after the trial was taken into account, given the fact that trial 1 is preceded by an open-field session.

##### 2.1. Correlation between neuronal activity and locomotor speed

Linear regression of firing rate (Hz) by locomotor speed (cm/s) was used to compute the intercept (Hz), slope (Hz*s/cm), R squared and Pearson’s correlation coefficient. The slope reflects the strength of the modulation of the neuronal activity by the locomotor activity and the Pearson’s correlation coefficient reflects the consistency of this modulation. For each neuron, linear regressions were computed in two conditions: during trials, and open-Field sessions. For rotarod trials, speed was estimated as 2πrv/60, where r is the radius of the rotarod axis in centimeters (1,5cm) and v is the rotation speed in r.p.m., allowing us to estimate the value of speed in cm/s. Mean firing rate was computed for each speed steps on the rotarod (8s bins). In order to assess the relationship between neuronal activity and speed while freely moving in the open-field, the speed was computed as explained in 2.2.3, resulting in a set of speeds expressed by the mouse during the open-field session. Those speeds were then interpolated in time considering the previous element in the set, allowing a continuous function that express speed in cm/s as a function of the time. To allow a valid comparison of the relationship between neuronal activity and speed on the rotarod and while freely moving in an open-field, the previously mentioned speed function was then discretized to match equivalent speed step on the rotarod (speed steps of 1 +/− 0.5 r.p.m.). The mean firing rate associated to each speed step in the open-field was then calculated as the number of spikes detected while being in the speed step, divided by the cumulative time spent in this speed step. The linear models were fitted using the *linregress* function from the *scipy* Python package. The modulation of activity by the speed along phases was compared for the different structures. Pearson’s correlation distributions for Acquisition, Consolidation and Maintenance phase were compared by using an overlapping index (non parametric measure of the effect size associated to the difference of the distributions between the compared structures). A continuous function was calculated from both distributions and smoothed by using a kernel density estimation (bandwidth = 0.05). Those functions were normalized in order to have their AUC equal to 1. The overlapping index was estimated by calculating the integral of the minimum of those functions over the interval of definition (Pastore and Calcagnì, 2019). This provided the overlapping index, a continuous value defined between 0 and 1 where the former would mean no overlap between the distributions and the latter would mean equal distributions.

#### 3. Chemogenetic

##### 3.1. Cerebellar outputs inactivation

We used evolved G-protein-coupled muscarinic receptors (hM4Di) that are selectively activated by the pharmacologically inert drug Clozapine-N-Oxide (CNO) (Alexander et al., 2009). In our study, non-cre and cre dependent version of the hM4Di receptor packaged into an AAV were used in order to facilitate the stereotaxic-based delivery and regionally restricted the expression of hM4Di. As demonstrated previously (Anaclet et al., 2018; Anaclet et al., 2014; Anaclet et al., 2015; Pedersen et al., 2017; Venner et al., 2016). hM4Di receptor and ligand are biologically inert in the absence of ligand. Moreover, at the administered dose of 1 mg/kg, CNO injection induces a maximum effect during the 1–3 h postinjection period (Anaclet et al., 2018; Anaclet et al., 2014) which enables us to confirm that during the whole duration of our protocols the CNO was still effective. We are therefore confident that the findings described in our study result from specific inhibition of the targeted neuronal population and not from a nonspecific effect of CNO or its metabolite clozapine (Gomez et al., 2017).

In order to globally inactivate the cerebellar outputs, stereotaxic surgeries were used to inject DREADD viral constructs bilaterally into the Dentate, Interposed and Fastigial nucleus. Mice were anesthetized with isoflurane for induction (3% in closed chamber during 4-5 minutes) and placed in the Kopf stereotaxic apparatus (model 942; PHYMEP, Paris, France) with mouse adapter (926-B, Kopf), and isoflurane vaporizer. Anesthesia was subsequently maintained at 1– 2% isoflurane. A longitudinal skin incision and removal of pericranial connective tissue exposed the bregma and lambda sutures of the skull. The coordinates for the Dentate nucleus injections were: 6.2 mm posterior to bregma, +/−2.3 mm lateral to the midline and −2.4 mm from dura while the Interposed injections were placed anteroposterior (AP) −6.0 mm, mediolateral (ML) = +/−1.5 mm in respect to bregma and dorsoventral (DV) −2.1 mm depth from dura. Finally, the Fastigial injections were placed −6.0 AP, +/−0.75 ML in respect to bregma and −2.1 depth from dura. Small holes were drilled into the skull and DREADD (AAV5-hSyn-hM4D(Gi)-mCherry, University of North Carolina Viral Core, 7.4 × 10^12^ vg per ml, 0.2 μl) or control (AAV5-hSyn-EGFP, UPenn Vector Core, the same concentration and amount) virus were delivered bilaterally via quartz micropipettes (QF 100-50-7.5, Sutter Instrument, Novato, USA) connected to an infusion pump (Legato 130 single syringe, 788130-KDS, KD Scientific, PHYMEP, Paris, France) at a speed of 100 nl/minutes. The micropipette was left in place for an additional 5 minutes to allow viral dispersion and prevent backflow of the viral solution into the injection syringe. The scalp wound was closed with surgical sutures, and the mouse was kept in a warm environment until resuming normal activity. All animals were given analgesic and fluids before and after the surgery.

In a separate set of mice, non-DREADD or DREADD (Dentate, Fastigial and Interposed) bundles of electrodes were implanted into the cerebellar nuclei, as described above. Both non-DREADD or DREADD injections and electrodes implantation were performed the same day. This experiment was performed in order to evaluate and validate that hM4D(Gi) receptors decrease the activity within the three cerebellar nuclei. Surgery, virus injections (AAV5-hSyn-hM4D(Gi)-mCherry or AAV5-hSyn-EGFP), coordinates (Fastigial: −6.0 AP, +/−0.75 ML, −2.1 depth from dura; Interposed: −6.0 AP, +/−1.5 ML, −2.1 depth from dura; Dentate: −6.2 AP, +/−2.3 ML, −2.4 depth from dura), chronic *in vivo* extracellular recordings and analysis were performed as we previously described above. One week following stereotaxic surgery to allow for virus expression, recordings at open-field were performed before and after CNO or saline (SAL) injection at the day 1, 4 and 7 of the accelerated rotarod task protocol. Mice were recorded for a 10 minutes baseline period followed by intraperitoneal injections of CNO 1mg/kg or SAL which were performed in a random sequence using a crossover design. After CNO or SAL injection, mice were recorded during 30 minutes before and 15 minutes after the accelerated rotarod task protocol.

##### 3.2. Cerebellar-thalamic outputs inactivation

In order to inhibit specifically cerebellar outputs to the centrolateral (CL) and/or ventral anterior lateral (VAL) thalamus we applied a chemogenetic pathway-specific approach (Boender et al., 2014). The technique comprises the combined use of a CRE-recombinase expressing canine adenovirus-2 (CAV-2) and an adeno-associated virus (AAV-hSyn-DIO-hM4D(Gi)-mCherry) that contains the floxed inverted sequence of the DREADD hM4D(Gi)-mCherry. It entails the infusion of these two viral vectors into two sites that are connected through direct neuronal projections and represent a neuronal pathway. AAV-hSyn-DIO-hM4D(Gi)-mCherry is infused in the site where the cell bodies are located, while CAV-2 is infused in the area that is innervated by the corresponding axons. After infection of axonal terminals, CAV-2 is transported towards the cell bodies and expresses CRE-recombinase (Kremer et al., 2000; Hnasko et al., 2006). AAV-hSyn-DIO-hM4D(Gi)-mCherry contains the floxed inverted sequence of hM4D(Gi)-mCherry, which is reoriented in the presence of CRE, prompting the expression of hM4D(Gi)-mCherry. This ensures that hM4D(Gi)-mCherry is not expressed in all AAV-hSyn-DIO-hM4D(Gi)-mCherry infected neurons, but exclusively in those that are also infected with CAV-2. Using the same procedures described above, 0.4 μl of the retrograde canine adeno-associated cre virus (CAV-2-cre, titter ≥ 2.5 × 10^8^) (Plateforme de Vectorologie de Montpellier, Montpellier, France) was bilaterally injected in the CL (from bregma: AP −1.70 mm, ML ±0.75 mm, DV −3.0 mm) and VAL (from bregma: AP −1.4 mm, ML ±1.0 mm, DV −3.5 mm). In addition, 0.2 μl of AAV-hSyn-DIO-hM4D(Gi)-mCherry (UNC Vector Core, Chapel Hill, NC, USA) was bilaterally injected one week later into the cerebellar nuclei, focusing on the Dentate (from bregma: AP −6.2 mm, ML ±2.3 mm, DV −2.4 mm) and Interposed (from bregma: AP −6.0 mm, ML ±1.5 mm, DV −2.1 mm) nucleus. Based on anatomical and functional evidences (Hintzen et ., 2018; Chen et al., 2014; Teune et al., 2000; Sakai 2000), in a group of mice we decided to inhibit those neurons that project from Dentate to CL and in another group we targeted those neurons that project from Interposed to VAL. All the stereotactic coordinates were determined based on The Mouse Brain Atlas (Paxinos and Franklin, 2004).

#### 4. Behavioral experiments design

Behavioral tests were performed one week following stereotaxic surgery to allow for virus expression. Balance beam, vertical pole, footprint patterns, grid test and fixed speed rotarod experiments were performed 30 minutes after CNO (1 mg/kg, ip) or SAL injections. Two different strategies were used for the accelerating rotarod motor learning task experiments: 1) CNO (1 mg/kg, ip) or SAL was injected every day 30 minutes before the 1st trial of the accelerated rotarod task. Four days later to ensure a proper CNO washout, mice were retested by receiving 7 trials for two consecutive daily sessions. Drug-free mice received CNO (1mg/kg) or SAL 30 minutes before the first trial in both days. The treatments were inverted meaning that those animals that received CNO during the preceding 7 days in this case were injected with SAL and the other way around. 2) CNO (1 mg/kg, ip) was injected 30 minutes after last trial at the day 1, 2 and 3; subsequently mice received SAL 30 minutes after last trial at the day 4, 5 and 6 of the accelerated rotarod task.

The DREADD ligand Clozapine-N-Oxide (CNO, TOCRIS, Bristol, UK) was dissolved in SAL (0.9% sodium chloride) and injected intraperitoneally at 1mg/kg.

#### 5. Histology

Mice were anesthetized with ketamine/xylazine (100 and 10 mg/kg, i.p., respectively) and rapidly perfused with ice-cold 4% paraformaldehyde in phosphate buffered SAL (PBS). The brains were carefully removed, postfixed in 4% paraformaldehyde for 24 h at 4 °C, cryoprotected in 20% sucrose in PBS. The whole brain was cut into 40-μm-thick coronal sections on a cryostat (Thermo Scientific HM 560; Waltham, MA, USA). The sections were mounted on glass slides sealed with Mowiol mounting medium (Mowiol^®^ 4-88; Sigma-Aldrich, France). Verification of virus injection site and DREADDs expression was assessed using a wide-field epifluorescent microscope (BX-43, Olympus, Waltham, MA, USA) using a mouse stereotaxic atlas (Paxinos and Franklin, 2004). We only kept mice showing a well targeted viral expression. Representative images of virus expression were acquired a Zeiss 800 Laser Scanning Confocal Microscope (×20 objective, NA 0.8) (Carl Zeiss, Jena, Germany). Images were cropped and annotated using Zeiss Zen 2 Blue Edition software (for example images, see Fig. 5A, D).

### QUANTIFICATION AND STATISTICAL ANALYSIS

Latency to fall in rotarod was analyzed by paired t-test for T1 vs T7 for each day (mean ± SEM). Distribution of numbers of trials in which cells were considered task related were compared using Mann-Whitney tests, Holm-Sidak corrected for multiple comparison between days. Evolution of the average firing rate (mean +/− SEM),) was normalized by subtracting the average firing rate during the active part of the open-field session before the first rotarod trial for each day (one-way ANOVA repeated measured followed by Dunett Posthoc test). Linear regression between speed (open-field and rotarod) and neuronal activity were performed and we extracted the Pearson’s correlation coefficient, slope and its associated p-value. Comparisons between slops in open-field vs rotarod were performed using Wilcoxon test. One-way ANOVA repeated measure followed by Tukey Posthoc test was used to compare values of significant linear regression slopes for rotarod (mean +/− SEM) between phases. Monotonous relationship between significant linear regression slopes for rotarod and their associated Pearson’s correlation coefficient were assessed using Spearman Rank test for each brain structure and phases. Distributions of significant Pearson’s correlation coefficient were compared between brain structures using Mann Whitney test for each phase, followed by the computation of the overlapping index η (used as a unbiased non parametric estimate of the effect size). Mean change in quiet state firing rate between open-field after and before were compared using one-way ANOVA followed by Tukey Posthoc test for each phase, while the evolution of mean quiet state (open-field sessions and Inter-trial periods for each phase) was analyzed using one-way ANOVA followed by Dunett Posthoc test. Mean firing rate was normalized by subtracting the open-field before (OF1). Modulation of firing rate for chemogenetic experiments were analyzed using one-way ANOVA followed by Tukey Posthoc test for each phase. Latency to fall (mean ± S.E.M) in rotarod for chemogenetic experiments were analyzed using one-way ANOVA repeated measure followed by two types for Posthoc tests: paired t-test for repeated measure comparison and independent t-test for cross group comparisons. Locomotor activity (velocity) in open-field (mean ± S.E.M) was analyzed using two-way ANOVA repeated measure (treatment×moment) followed by t-test Posthoc (comparisons between treatments for each open-field session). Fixed speed rotarod (mean ± S.E.M) was analyzed using two-way ANOVA repeated measure (treatment×speed) followed by t-test Posthoc (comparisons between treatments for each speed steps). Footprint patterns parameters, horizontal bar, vertical pole and grid test were analyzed using one-way ANOVA. Data are represented as boxplots (median quartiles and interquartile range plus outliers).

For the figure 6, to get a robust estimate of the initial and final performance of each day, we performed a linear regression on the latency to fall of each day and each animal; the within-day and overnight loss was estimated from the start- and end-points of each regression segment. To estimate the interdependence of initial performance of the day, within day learning and inter-day learning, we used Deming linear regression, assuming equal variance of the noise of the measured quantity on the x- and y-axis. Confidence intervals were obtained by bootstrap.

