## Supplementary material for "Dual contributions of cerebellar-thalamic networks to learning and offline consolidation of a complex motor task"

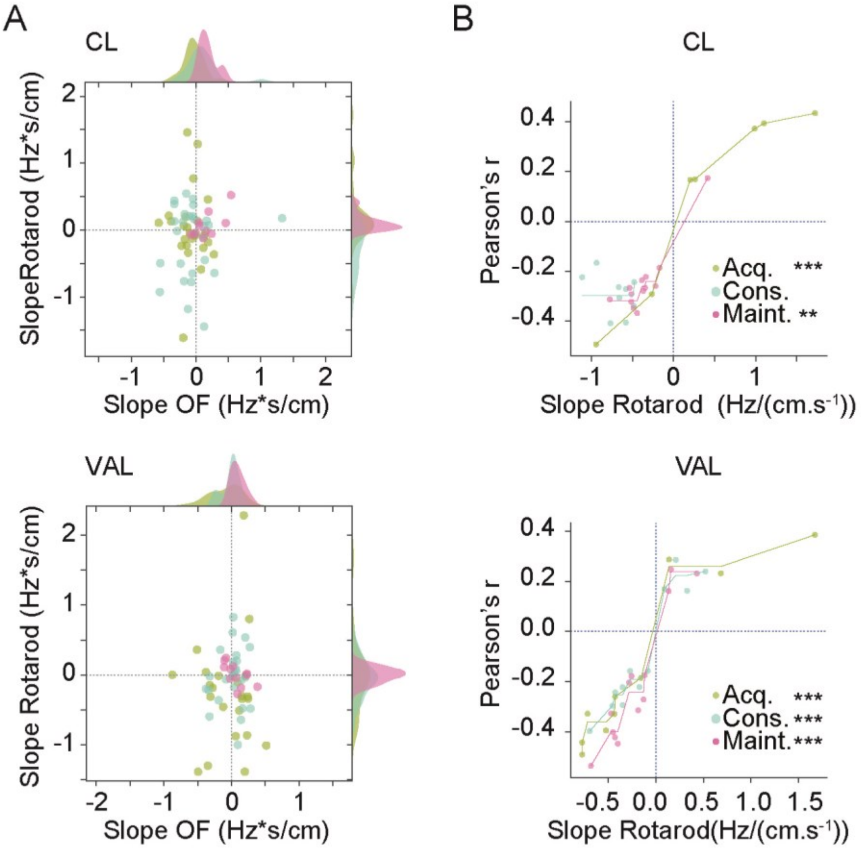


**Fig Sup1. Centrolateral and ventral anterior lateral thalamus display a sensitivity to rotarod speed.** a) Scatter plot showing the slope of linear regression explaining the firing rate by the speed, in the open-field versus the rotarod for each neuron in centrolateral thalamus (CL, top) and ventral anterior lateral thalamus (VAL, bottom) during Acquisition (Acq.), Consolidation (Cons.) and Maintenance (Maint.). Marginal axes show the histograms of the distributions of slopes, smoothed using a Gaussian kernel density estimate ($\sigma$=0.05). b) Scatter plot showing the correspondence of slope of linear regression on rotarod versus the associated Pearson correlation coefficient for each neuron for CL (top) and VAL (bottom), during Acquisition (Acq.), Consolidation (Cons.) and Maintenance (Maint.). The lines represent the isotonic regression of the Pearson's r by the slope on rotarod (**$p<0.01$, ***$p<0.001$ Spearman Rank test).

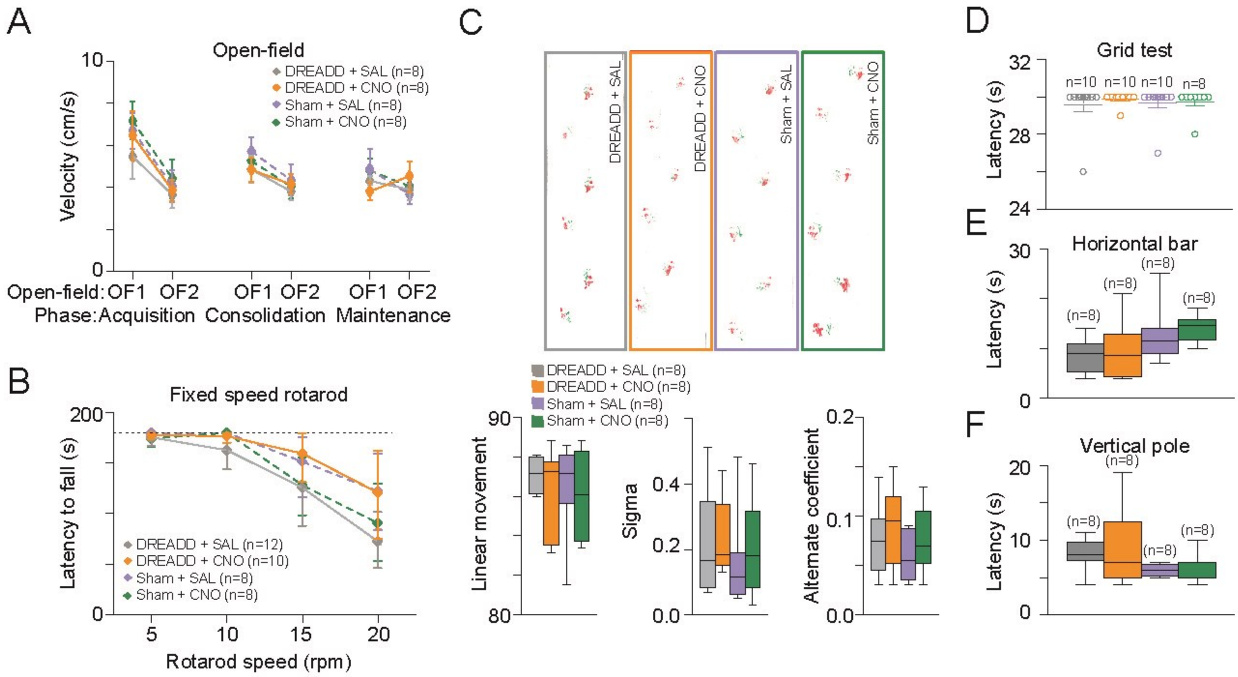


**Fig Sup2. Cerebellar nuclei inhibition did not affect execution and fatigue, locomotion, motor coordination, balance and strength.** a) Locomotor activity (Velocity) in DREADD and non-DREADD (Sham) injected mice after CNO or SAL injection during open-field sessions before (OF1) and after (OF2) rotarod for Acquisition, Consolidation and Maintenance (**p<0.01 t-test OF1 vs OF2). b) Latency to fall during fixed speed rotarod (5, 10, 15, 20, 25 r.p.m.) for all experimental groups. One way repeated measure ANOVA was performed on averaged values for all the speed steps in each experimental group followed by a Tukey Posthoc pairwise comparison. c) Footprint patterns were quantitatively assessed for 3 parameters as shown on representative footprint patterns (top) for all experimental groups. Three parameters are represented graphically: linear movement (bottom left), sigma (bottom middle) and alternation coefficient (bottom right). d) Latency reflecting the time before falling from the grid. 30 seconds of cut-off of was established as the maximum latency (dotted line on figure). e) Latency to cross the horizontal bar (balance beam test) for all experimental groups. f) Latency to reach home cage in vertical pole test for all experimental groups. *p<0.05 One Way ANOVA followed by a PostHoc Tukey test. *n* indicates the number of mice.

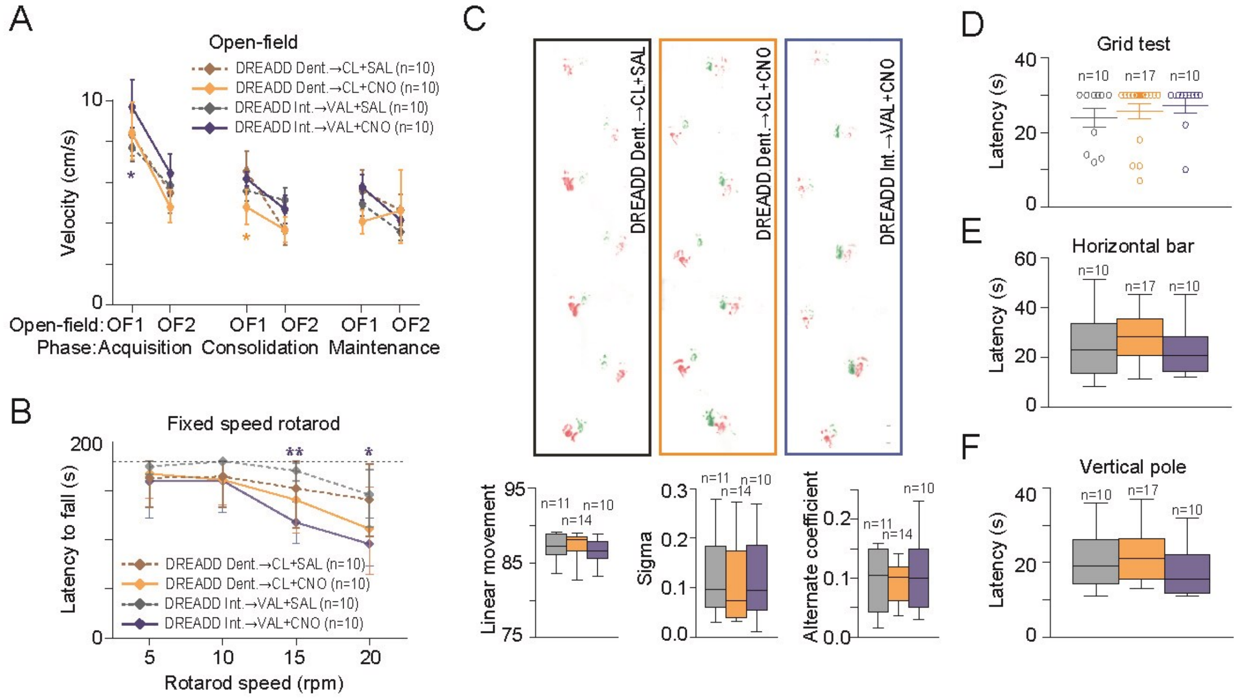


**Fig Sup3. Dentate-centrolateral and Interposed-ventral anterior lateral inhibition did not affect execution and fatigue, locomotion, motor coordination, balance and strength**. a) Locomotor activty (Velocity) in DREADD injected mice after CNO or SAL injection during open-fields sessions before (OF1) and after (OF2) rotarod for Acquisition, Consolidation and Maintenance (**p<0.01 t-test OF1 vs OF2). b) Latency to fall during fixed speed rotarod (5, 10, 15, 20 r.p.m.) for all experimental groups. One way repeated measure ANOVA was performed on averaged values for all the speed steps in each experimental group followed by a Tukey Posthoc pairwise comparison. c) Footprint patterns were quantitatively assessed for 3 parameters as shown on representative footprint patterns (top) for all experimental groups. Three parameters are represented graphically: linear movement (bottom left), sigma (bottom middle and alternation coefficient (bottom right). d) Latency reflecting the time before falling from the grid. 30 seconds of cut-off of was established as the maximum latency (dotted line on figure). e) Latency to cross the horizontal bar (balance beam test) for all experimental groups. f) Latency to reach home cage in vertical pole test for all experimental groups. *p<0.05 One Way ANOVA followed by a PostHoc Tukey test. DCN, deep cerebellar nuclei; CL, centrolateral thalamus; VAL, ventral anterior lateral thalamus. *n* indicates the number of mice.

| Day | Moment 1 | Moment 2 | df | Statistic | CI 2.5% | CI 97.5% | p-value | Cohen's D | Sig. |
|---|---|---|---|---|---|---|---|---|---|
| 1.00 | Trial 1 | Trial 7 | 28.22 | -3.75 | -82.52 | -24.23 | 0.00 | -1.33 | ** |
| 2.00 | Trial 1 | Trial 7 | 25.19 | -2.24 | -68.33 | 11.45 | 0.04 | -0.52 | * |
| 3.00 | Trial 1 | Trial 7 | 28.99 | -0.48 | -39.71 | 29.21 | 0.64 | -0.11 | ns |
| 4.00 | Trial 1 | Trial 7 | 29.97 | -1.67 | -78.88 | 21.38 | 0.12 | -0.41 | ns |
| 5.00 | Trial 1 | Trial 7 | 29.96 | -1.76 | -58.02 | 11.15 | 0.10 | -0.49 | ns |
| 6.00 | Trial 1 | Trial 7 | 29.81 | 1.07 | -15.55 | 45.42 | 0.30 | 0.35 | ns |
| 7.00 | Trial 1 | Trial 7 | 29.76 | -0.50 | -53.62 | 35.49 | 0.62 | -0.15 | ns |

Supplementary Table 1: Statistics for Fig 1 A.

| Structure | Test | Group 1 | Group 2 | Statistic | p-value | Sig. |
|---|---|---|---|---|---|---|
| Dentate | Mann Whitney(Holm Sidak corrected) | Day 1 (n=55) | Day 4 (n=53) | 1111.000 | 0.045 | * |
| Dentate | Mann Whitney(Holm Sidak corrected) | Day 1 (n=55) | Day 7 (n=51) | 1024.500 | 0.028 | * |
| Dentate | Mann Whitney(Holm Sidak corrected) | Day 4 (n=53) | Day 7 (n=51) | 1244.500 | 0.431 | n.s |
| Interposed | Mann Whitney(Holm Sidak corrected) | Day 1 (n=62) | Day 4 (n=67) | 1725.000 | 0.065 | n.s |
| Interposed | Mann Whitney(Holm Sidak corrected) | Day 1 (n=62) | Day 7 (n=93) | 1975.000 | <0.001 | *** |
| Interposed | Mann Whitney(Holm Sidak corrected) | Day 4 (n=67) | Day 7 (n=93) | 2623.500 | 0.052 | n.s |
| CL | Mann Whitney(Holm Sidak corrected) | Day 1 (n=27) | Day 4 (n=27) | 220.500 | 0.030 | * |
| CL | Mann Whitney(Holm Sidak corrected) | Day 1 (n=27) | Day 7 (n=25) | 264.000 | <0.001 | *** |
| CL | Mann Whitney(Holm Sidak corrected) | Day 4 (n=27) | Day 7 (n=25) | 380.500 | 0.400 | n.s |
| VAL | Mann Whitney(Holm Sidak corrected) | Day 1 (n=23) | Day 4 (n=26) | 225.500 | 0.251 | n.s |
| VAL | Mann Whitney(Holm Sidak corrected) | Day 1 (n=23) | Day 7 (n=25) | 354.000 | 0.251 | n.s |
| VAL | Mann Whitney(Holm Sidak corrected) | Day 4 (n=26) | Day 7 (n=25) | 474.000 | ‘0.013 | * |
| Cortex | Mann Whitney(Holm Sidak corrected) | Day 1 (n=55) | Day 4 (n=50) | 225.500 | 0.006 | ** |
| Cortex | Mann Whitney(Holm Sidak corrected) | Day 1 (n=55) | Day 7 (n=49) | 354.000 | 0.140 | n.s |
| Cortex | Mann Whitney(Holm Sidak corrected) | Day 4 (n=50) | Day 7 (n=49) | 474.000 | <0.001 | *** |
| Striatum | Mann Whitney(Holm Sidak corrected) | Day 1 (n=107) | Day 4 (n=88) | 4907.500 | 0.601 | n.s |
| Striatum | Mann Whitney(Holm Sidak corrected) | Day 1 (n=107) | Day 7 (n=116) | 7743.500 | 0.003 | ** |
| Striatum | Mann Whitney(Holm Sidak corrected) | Day 4 (n=88) | Day 7 (n=116) | 6317.000 | 0.006 | ** |

Supplementary Table 2: Statistics for Fig1 E.

| Structure | Phase | Day | ANOVA | p-value | Sig. | n (cells/mouse) | Moment 1 | Moment 2 | Mean Difference | CI 2.5% | CI 97.5% | p-value | Sig. |
|---|---|---|---|---|---|---|---|---|---|---|---|---|---|
| Dentate | Acquisition | 1.00 | F(3,363)=6.854 | <0.001 | *** | 122/9 | Trial 1 | Before | -1.24 | -2.51 | 0.03 | 0.057 | n.s. |
| | | | | | | | Trial 4 | Before | -2.03 | -3.30 | -0.76 | <0.001 | *** |
| | | | | | | | Trial 7 | Before | -2.19 | -3.46 | -0.92 | <0.001 | *** |
| Dentate | Consolidation | 4.00 | F(3,267)=3.572 | 0.015 | * | 90/8 | Trial 1 | Before | -1.00 | -2.41 | 0.40 | 0.224 | n.s. |
| | | | | | | | Trial 4 | Before | -1.27 | -2.67 | 0.14 | 0.088 | n.s. |
| | | | | | | | Trial 7 | Before | -1.93 | -3.33 | -0.52 | 0.003 | ** |
| Dentate | Maintenance | 7.00 | F(3,327)=13 | <0.001 | *** | 110/9 | Trial 1 | Before | -1.34 | -2.05 | -0.64 | <0.001 | *** |
| | | | | | | | Trial 4 | Before | -1.76 | -2.46 | -1.06 | <0.001 | *** |
| | | | | | | | Trial 7 | Before | -1.28 | -1.99 | -0.58 | <0.001 | *** |
| Interposed | Acquisition | 1.00 | F(3,276)=1.995 | 0.115 | n.s | 93/8 | | | | | | | |
| Interposed | Consolidation | 4.00 | F(3,270)=34.88 | <0.001 | *** | 91/7 | Trial 1 | Before | -3.52 | -4.75 | -2.29 | <0.001 | *** |
| | | | | | | | Trial 4 | Before | -4.32 | -5.55 | -3.09 | <0.001 | *** |
| | | | | | | | Trial 7 | Before | -4.88 | -6.12 | -3.65 | <0.001 | *** |
| Interposed | Maintenance | 7.00 | F(3,360)=5.694 | <0.001 | *** | 121/8 | Trial 1 | Before | -0.98 | -1.98 | 0.02 | 0.055 | n.s. |
| | | | | | | | Trial 4 | Before | -1.52 | -2.52 | -0.52 | 0.001 | ** |
| | | | | | | | Trial 7 | Before | -1.53 | -2.52 | -0.53 | <0.001 | *** |
| CL | Acquisition | 1.00 | F(3,207)=14.45 | <0.001 | *** | 70/7 | Trial 1 | Before | 1.20 | 0.75 | 1.64 | <0.001 | *** |
| | | | | | | | Trial 4 | Before | 0.62 | 0.17 | 1.06 | 0.003 | ** |
| | | | | | | | Trial 7 | Before | 0.33 | -0.12 | 0.77 | 0.199 | n.s. |
| CL | Consolidation | 4.00 | F(3,228)=17.25 | <0.001 | *** | 77/7 | Trial 1 | Before | 1.36 | 0.91 | 1.81 | <0.001 | *** |
| | | | | | | | Trial 4 | Before | 0.61 | 0.16 | 1.06 | 0.004 | ** |
| | | | | | | | Trial 7 | Before | 0.76 | 0.31 | 1.20 | <0.001 | *** |
| CL | Maintenance | 7.00 | F(3,195)=9.841 | <0.001 | *** | 66/6 | Trial 1 | Before | 0.91 | 0.44 | 1.37 | <0.001 | *** |
| | | | | | | | Trial 4 | Before | 0.94 | 0.48 | 1.41 | <0.001 | *** |
| | | | | | | | Trial 7 | Before | 0.67 | 0.20 | 1.13 | 0.002 | ** |
| VAL | Acquisition | 1.00 | F(3,255)=9.199 | <0.001 | *** | 86/7 | Trial 1 | Before | 0.96 | 0.48 | 1.44 | <0.001 | *** |
| | | | | | | | Trial 4 | Before | 0.35 | -0.13 | 0.83 | 0.203 | n.s. |
| | | | | | | | Trial 7 | Before | 0.08 | -0.41 | 0.56 | 0.968 | n.s. |
| VAL | Consolidation | 4.00 | F(3,213)=13.2 | <0.001 | *** | 72/7 | Trial 1 | Before | 0.76 | 0.43 | 1.09 | <0.001 | *** |
| | | | | | | | Trial 4 | Before | 0.67 | 0.33 | 1.00 | <0.001 | *** |
| | | | | | | | Trial 7 | Before | 0.74 | 0.41 | 1.07 | <0.001 | *** |
| VAL | Maintenance | 7.00 | F(3,177)=10.89 | <0.001 | *** | 60/6 | Trial 1 | Before | 0.57 | 0.25 | 0.89 | <0.001 | *** |
| | | | | | | | Trial 4 | Before | 0.68 | 0.36 | 1.00 | <0.001 | *** |
| | | | | | | | Trial 7 | Before | 0.64 | 0.32 | 0.95 | <0.001 | *** |

Supplementary Table 3: Statistics for Fig 1 F.

| Structure | Episode | Pearson's r | p-value | Sig. |
|---|---|---|---|---|
| Dentate | Rotarod | -0.62 | <0.001 | *** |
| | Open-Field | 0.15 | 0.054 | n.s. |
| Interposed | Rotarod | -0.43 | <0.001 | *** |
| | Open-Field | 0.17 | 0.027 | * |

Supplementary Table 4: Statistics for Fig 2 A.

| Structure | Phase | Day | Test | Group 1 | Group 2 | Statistic | p-value | Sig. | n(cells/mouse) |
|---|---|---|---|---|---|---|---|---|---|
| Dentate | Acquisition | 1.00 | Wilcoxon | Slopes OF | Slopes Trials | 45.000 | <0.001 | *** | 55/4 |
| Dentate | Consolidation | 4.00 | Wilcoxon | Slopes OF | Slopes Trials | 50.000 | <0.001 | *** | 53/4 |
| Dentate | Maintenance | 7.00 | Wilcoxon | Slopes OF | Slopes Trials | 274.000 | <0.001 | *** | 51/4 |
| Interposed | Acquisition | 1.00 | Wilcoxon | Slopes OF | Slopes Trials | 0.000 | <0.001 | *** | 62/4 |
| Interposed | Consolidation | 4.00 | Wilcoxon | Slopes OF | Slopes Trials | 59.000 | <0.001 | *** | 67/4 |
| Interposed | Maintenance | 7.00 | Wilcoxon | Slopes OF | Slopes Trials | 366.000 | <0.001 | *** | 93/4 |
| CL | Acquisition | 1.00 | Wilcoxon | Slopes OF | Slopes Trials | 92.000 | 0.020 | * | 27/2 |
| CL | Consolidation | 4.00 | Wilcoxon | Slopes OF | Slopes Trials | 62.000 | 0.002 | ** | 43888 |
| CL | Maintenance | 7.00 | Wilcoxon | Slopes OF | Slopes Trials | 13.000 | <0.001 | *** | 25 |
| VAL | Acquisition | 1.00 | Wilcoxon | Slopes OF | Slopes Trials | 128.000 | 0.761 | n.s. | 23 |
| VAL | Consolidation | 4.00 | Wilcoxon | Slopes OF | Slopes Trials | 170.000 | 0.889 | n.s. | 43887 |
| VAL | Maintenance | 7.00 | Wilcoxon | Slopes OF | Slopes Trials | 28.000 | <0.001 | *** | 43886 |

Supplementary Table 5: Statistics for Fig 2 C.

| Structure | ANOVA | p-value | Sig. | n (cells/mouse) | Phase 1 | Phase 2 | Mean Difference | CI 2.5% | CI 97.5% | p-value | Sig. |
|---|---|---|---|---|---|---|---|---|---|---|---|
| Dentate | F(2,112)=4.041 | 0.02 | * | 118/4 | Consolidation | Acquisition | 0.12 | -0.09 | 0.33 | 0.377 | n.s. |
| | | | | | Maintenance | Acquisition | -0.14 | -0.37 | 0.09 | 0.321 | n.s. |
| | | | | | Maintenance | Consolidation | -0.26 | -0.47 | -0.04 | 0.013 | * |
| Interposed | F(2,155)=4.824 | 0.009 | ** | 161/4 | Consolidation | Acquisition | 0.45 | 0.06 | 0.85 | 0.021 | * |
| | | | | | Maintenance | Acquisition | 0.53 | 0.12 | 0.94 | 0.007 | ** |
| | | | | | Maintenance | Consolidation | 0.07 | -0.24 | 0.38 | 0.84 | n.s. |
| CL | F(2,25)=11.07 | <0.001 | *** | 29/2 | Consolidation | Acquisition | -1.14 | -1.71 | -0.56 | <0.001 | *** |
| | | | | | Maintenance | Acquisition | -0.80 | -1.33 | -0.26 | <0.001 | *** |
| | | | | | Maintenance | Consolidation | 0.34 | -0.16 | 0.83 | 0.249 | n.s. |
| VAL | F(2,31)=0.1156 | 0.891 | n.s. | 35/2 | - no posthoc | | | | | | |

Supplementary Table 6: Statistics for Fig 2D.

| Structure | Phase | Day | Test | Group 1 | Group 2 | Rho | p-value | Sig. | n |
|---|---|---|---|---|---|---|---|---|---|
| Dentate | Acquisition | 1.00 | Spearman rank test | Slopes Trials | Pearson r Trials | 0.248 | 0.134 | n.s. | 38/4 |
| Dentate | Consolidation | 4.00 | Spearman rank test | Slopes Trials | Pearson r Trials | 0.585 | <0.001 | *** | 45/4 |
| Dentate | Maintenance | 7.00 | Spearman rank test | Slopes Trials | Pearson r Trials | 0.620 | <0.001 | *** | 35/4 |
| Interposed | Acquisition | 1.00 | Spearman rank test | Slopes Trials | Pearson r Trials | 0.443 | 0.014 | *** | 43951 |
| Interposed | Consolidation | 4.00 | Spearman rank test | Slopes Trials | Pearson r Trials | 0.704 | <0.001 | *** | 57/4 |
| Interposed | Maintenance | 7.00 | Spearman rank test | Slopes Trials | Pearson r Trials | 0.591 | <0.001 | *** | 74/4 |
| CL | Acquisition | 1.00 | Spearman rank test | Slopes Trials | Pearson r Trials | 1.000 | <0.001 | *** | 43868 |
| CL | Consolidation | 4.00 | Spearman rank test | Slopes Trials | Pearson r Trials | -0.300 | 0.423 | n.s. | 43870 |
| CL | Maintenance | 7.00 | Spearman rank test | Slopes Trials | Pearson r Trials | 0.758 | 0.003 | ** | 43874 |
| VAL | Acquisition | 1.00 | Spearman rank test | Slopes Trials | Pearson r Trials | 0.956 | <0.001 | *** | 43871 |
| VAL | Consolidation | 4.00 | Spearman rank test | Slopes Trials | Pearson r Trials | 0.937 | <0.001 | *** | 43873 |
| VAL | Maintenance | 7.00 | Spearman rank test | Slopes Trials | Pearson r Trials | 0.896 | <0.001 | *** | 43874 |

Supplementary Table 7: Statistics for Fig 2 E.

| Phase | Day | Test | Group 1 | Group 2 | Statistic | p-value | Sig. | Eta |
|---|---|---|---|---|---|---|---|---|
| Acquisition | 1.00 | Mann Whitney | Dentate (n=38) | CL (n=7) | 49.000 | 0.001 | ** | 0.243 |
| Consolidation | 4.00 | Mann Whitney | Dentate (n=45) | CL (n=9) | 168.000 | 0.811 | n.s. | 0.885 |
| Maintenance | 7.00 | Mann Whitney | Dentate (n=35) | CL (n=13) | 154.000 | 0.090 | n.s. | 0.632 |
| Acquisition | 1.00 | Mann Whitney | Interposed (n=30) | VAL (n=10) | 124.000 | 0.426 | n.s. | 0.646 |
| Consolidation | 4.00 | Mann Whitney | Interposed (n=57) | VAL (n=12) | 83.000 | <0.001 | *** | 0.398 |
| Maintenance | 7.00 | Mann Whitney | Interposed (n=74) | VAL (n=13) | 308.000 | 0.040 | * | 0.712 |

Supplementary Table 8: Statistics for Fig 2 F.

| Structure | ANOVA | p-value | Sig. | Group 1 | Group 2 | Mean Difference | CI 2.5% | CI 97.5% | p-value | Sig. | n (cells/mouse) |
|---|---|---|---|---|---|---|---|---|---|---|---|
| Dentate | F(2,292)=17.72 | <0.001 | *** | Acquisition Quiet OF2 | Acquisition Quiet OF1 | -2.95 | -6.09 | 0.20 | 0.067 | n.s. | (94/6) |
| | | | | Consolidation Quiet OF2 | Consolidation Quiet OF1 | -1.03 | -4.01 | 1.95 | 0.498 | n.s. | (99/9) |
| | | | | Maintenance Quiet OF2 | Maintenance Quiet OF1 | -3.01 | -4.94 | -1.09 | 0.002 | ** | (110/9) |
| Interposed | F(2,271)=5.476 | 0.005 | ** | Acquisition Quiet OF2 | Acquisition Quiet OF1 | -1.65 | -3.26 | -0.05 | 0.044 | * | (63/6) |
| | | | | Consolidation Quiet OF2 | Consolidation Quiet OF1 | -1.14 | -4.34 | 2.05 | 0.482 | n.s. | (97/8) |
| | | | | Maintenance Quiet OF2 | Maintenance Quiet OF1 | -3.36 | -4.99 | -1.72 | <0.001 | *** | (121/8) |
| CL | F(2,226)=2.866 | 0.059 | n.s. | | | | | | | | |
| VAL | F(2,236)=7.275 | <0.001 | *** | Acquisition Quiet OF2 | Acquisition Quiet OF1 | 0.16 | -0.54 | 0.86 | 0.655 | n.s. | (93/7) |
| | | | | Consolidation Quiet OF2 | Consolidation Quiet OF1 | -0.41 | -1.07 | 0.25 | 0.221 | n.s. | (79/7) |
| | | | | Maintenance Quiet OF2 | Maintenance Quiet OF1 | -0.23 | -0.90 | 0.45 | 0.512 | n.s. | (73/7) |

Supplementary Table 9: Statistics for Fig 3 A.

| Factor | ANOVA | p-value | Sig. | Group 1 : Structure Phase | Group 2 : Structure Phase | Mean Difference | CI 2.5% | CI 97.5% | p-value | Sig. |
|---|---|---|---|---|---|---|---|---|---|---|
| Structure | F(1,580)=9.395 | 0.002 | ** | | | | | | | |
| Phase | F(2,580)=15.404 | <0.001 | *** | | | | | | | |
| Structure:Phase | F(2,580)=11.211 | <0.001 | *** | Dentate Acq. (n=94/6) | Dentate Cons. (n=101/9) | 0.98 | -0.73 | 2.69 | 0.37 | n.s. |
| | | | | Dentate Acq. (n=94/6) | Dentate Maint. (n=110/9) | 0.48 | -1.19 | 2.16 | 0.75 | n.s. |
| | | | | Dentate Cons. (n=101/9) | Dentate Maint. (n=110/9) | -0.49 | -2.14 | 1.15 | 0.74 | n.s. |
| | | | | Interposed Acq. (n=63/6) | Interposed Cons. (n=97/8) | 4.69 | 2.34 | 7.04 | 0.00 | ** |
| | | | | Interposed Acq. (n=63/6) | Interposed Maint. (n=121/8) | 5.28 | 3.02 | 7.54 | 0.00 | ** |
| | | | | Interposed Cons. (n=97/8) | Interposed Maint. (n=121/8) | 0.59 | -1.39 | 2.57 | 0.74 | n.s. |

Supplementary Table 10: Statistics for Fig 3 OF2.

| Structure | Test | Moment 1 | Moment 2 | Acquisition | Consolidation | Maintenance |
|---|---|---|---|---|---|---|
| Dentate | ANOVA | | | F(8,328)=9.077 p<0.001 *** | F(8,416)=21.71 p<0.001 *** | F(8,400)=27.32 p<0.001 *** |
| | n (cells/mouse) | | | 55/4 | 53/4 | 51/4 |
| | Posthoc Dunnett | Intertrial 1 | Quiet OF1 | 0.999 | <0.001 *** | 0.081 |
| | | Intertrial 2 | Quiet OF1 | 0.389 | <0.001 *** | <0.001 *** |
| | | Intertrial 3 | Quiet OF1 | 0.006 ** | <0.001 *** | <0.001 *** |
| | | Intertrial 4 | Quiet OF1 | 0.007 ** | <0.001 *** | <0.001 *** |
| | | Intertrial 5 | Quiet OF1 | <0.001 *** | <0.001 *** | <0.001 *** |
| | | Intertrial 6 | Quiet OF1 | <0.001 *** | <0.001 *** | <0.001 *** |
| | | Intertrial 7 | Quiet OF1 | 0.002 ** | <0.001 *** | <0.001 *** |
| | | Quiet OF2 | Quiet OF1 | 0.044 * | <0.001 *** | <0.001 *** |
| Interposed | ANOVA | | | F(8,296)=28.78 p<0.001 *** | F(8,528)=36.25 p<0.001 *** | F(8,736)=18.55 p<0.001 *** |
| | n (cells/mouse) | | | 62/4 | 67/4 | 93/4 |
| | Posthoc Dunnett | Intertrial 1 | Quiet OF1 | <0.001 *** | <0.001 *** | <0.001 *** |
| | | Intertrial 2 | Quiet OF1 | 0.003 ** | <0.001 *** | <0.001 *** |
| | | Intertrial 3 | Quiet OF1 | 0.022 * | <0.001 *** | <0.001 *** |
| | | Intertrial 4 | Quiet OF1 | 1 | <0.001 *** | <0.001 *** |
| | | Intertrial 5 | Quiet OF1 | 0.054 | <0.001 *** | <0.001 *** |
| | | Intertrial 6 | Quiet OF1 | 0.377 | <0.001 *** | <0.001 *** |
| | | Intertrial 7 | Quiet OF1 | 0.015* | <0.001 *** | <0.001 *** |
| | | Quiet OF2 | Quiet OF1 | <0.001 *** | <0.001 *** | <0.001 *** |
| CL | ANOVA | | | F(8,208)=6.205 p<0.001 *** | F(8,208)=0.8426 p=0.566 | F(8,192)=29.16 p<0.001 *** |
| | n (cells/mouse) | | | 27/2 | 43888 | 25 |
| | Posthoc Dunnett | Intertrial 1 | Quiet OF1 | <0.001 *** | 0.96 | <0.001 *** |
| | | Intertrial 2 | Quiet OF1 | 0.046 * | 0.95 | <0.001 *** |
| | | Intertrial 3 | Quiet OF1 | 0.001 ** | 1.00 | <0.001 *** |
| | | Intertrial 4 | Quiet OF1 | 0.213 | 0.87 | <0.001 *** |
| | | Intertrial 5 | Quiet OF1 | 0.108 | 1.00 | <0.001 *** |
| | | Intertrial 6 | Quiet OF1 | 0.350 | 0.93 | <0.001 *** |
| | | Intertrial 7 | Quiet OF1 | 0.910 | 1.00 | <0.001 *** |
| | | Quiet OF2 | Quiet OF1 | 0.366 | 1.00 | 0.61 |
| VAL | ANOVA | | | F(8,176)=4.58 p<0.001 *** | F(8,200)=6.263 p<0.001 *** | F(8,184)=28.15 p<0.001 *** |
| | n (cells/mouse) | | | 23 | 43887 | 43886 |
| | Posthoc Dunnett | Intertrial 1 | Quiet OF1 | <0.001 *** | 0.001 ** | <0.001 *** |
| | | Intertrial 2 | Quiet OF1 | 0.30 | <0.001 *** | <0.001 *** |
| | | Intertrial 3 | Quiet OF1 | 0.09 | 0.908 | <0.001 *** |
| | | Intertrial 4 | Quiet OF1 | s | 0.001 ** | <0.001 *** |
| | | Intertrial 5 | Quiet OF1 | 0.29 | 0.003 ** | <0.001 *** |
| | | Intertrial 6 | Quiet OF1 | 0.40 | 0.851 | <0.001 *** |
| | | Intertrial 7 | Quiet OF1 | 1.00 | 0.653 | <0.001 *** |
| | | Quiet OF2 | Quiet OF1 | 0.85 | 0.999 | 0.025 * |

Supplementary Table 11: Statistics for Fig 3 B.

| Group | ANOVA | p-value | Sig. | Group 1 | Group 2 | df | Statistic | CI 2.5% | CI 97.5% | p-value | Cohen's D | Sig. |
|---|---|---|---|---|---|---|---|---|---|---|---|---|
| DREADD + CNO | F(1,434)=294.719 | <0.001 | *** | Acq. ctrl | Acq. CNO | 136.61 | 12.28 | 6.66 | 10.04 | <0.001 | 1.51 | *** |
| | | | | Cons. ctrl | Cons. CNO | 98.63 | 10.55 | 6.78 | 10.19 | <0.001 | 1.75 | *** |
| | | | | Maint. ctrl | Maint. CNO | 102.40 | 14.66 | 6.27 | 9.24 | <0.001 | 1.75 | *** |
| DREADD + SAL | F(1,566)=2.009 | 0.157 | n.s. | | | | | | | | | |
| Sham + CNO | F(1,592)=0.014 | 0.905 | n.s. | | | | | | | | | |
| Sham + SAL | F(1,306)=5.665 | 0.018 | * | Acq. ctrl | Acq. SAL | 107.96 | -2.24 | -1.58 | 1.15 | 0.029 | -0.06 | * |
| | | | | Cons. ctrl | Cons. SAL | 90.66 | 5.53 | 0.16 | 3.86 | <0.001 | 0.45 | *** |
| | | | | Maint. ctrl | Maint. SAL | 99.25 | 4.78 | -0.11 | 5.63 | <0.001 | 0.37 | *** |

Supplementary Table 12: Statistics for Fig 4 H (Before-After).

| Group | Time of treatment | Day | ANOVA | p-value | Sig. |
|---|---|---|---|---|---|
| DCN | During task | 1.00 | F(3,45)=0.9452 | 0.426 | n.s. |
| | | 2.00 | F(3,45)=4.021 | 0.013 | * |
| | | 3.00 | F(3,45)=4.212 | 0.01 | * |
| | | 4.00 | F(3,45)=6.601 | <0.001 | *** |
| | | 5.00 | F(3,45)=7.899 | <0.001 | *** |
| | | 6.00 | F(3,45)=13.33 | <0.001 | *** |
| | | 7.00 | F(3,45)=7.741 | <0.001 | *** |
| | After task | 1.00 | F(3,30)=0.414 | 0.744 | n.s. |
| | | 2.00 | F(3,30)=6.215 | 0.002 | ** |
| | | 3.00 | F(3,30)=3.868 | 0.019 | * |
| | | 4.00 | F(3,30)=2.26 | 0.101 | n.s. |
| | | 5.00 | F(3,30)=1.413 | 0.258 | n.s. |
| | | 6.00 | F(3,30)=0.256 | 0.856 | n.s. |
| | | 7.00 | F(3,30)=1.971 | 0.1396 | n.s. |

Supplementary Table 13: Statistics for Fig 4 I J.

| Group | Time of treatment | Comparaison | Moment 1 | Moment 2 | DREADD + CNO | DREADD + SAL | Sham + CNO | Sham + SAL |
|---|---|---|---|---|---|---|---|---|
| DCN | During task | Within-day | Trial 7 Day1 | Trial 1 Day 1 | <0.001 *** | 0.008 ** | 0.19 | 0.002 ** |
| | | | Trial 7 Day2 | Trial 1 Day 2 | 0.007 ** | 1 | 0.19 | 0.464 |
| | | | Trial 7 Day3 | Trial 1 Day 3 | 0.015 * | 1 | 1 | 0.464 |
| | | | Trial 7 Day4 | Trial 1 Day 4 | 0.016 * | 1 | 0.27 | 1 |
| | | | Trial 7 Day5 | Trial 1 Day 5 | 0.255 | 1 | 1 | 0.847 |
| | | | Trial 7 Day6 | Trial 1 Day 6 | 0.196 | 1 | 1 | 1 |
| | | | Trial 7 Day7 | Trial 1 Day 7 | 0.196 | 1 | 1 | 1 |
| | | Next-day | Trial 1 Day2 | Trial 7 Day 1 | 0.031 * | 1 | 1 | 0.9 |
| | | | Trial 1 Day3 | Trial 7 Day 2 | 0.223 | 0.58 | 0.15 | 0.94 |
| | | | Trial 1 Day4 | Trial 7 Day 3 | 0.037 * | 1 | 0.29 | 1 |
| | | | Trial 1 Day5 | Trial 7 Day 4 | 0.071 | 1 | 0.39 | 0.27 |
| | | | Trial 1 Day6 | Trial 7 Day 5 | 0.223 | 0.26 | 1 | 1 |
| | | | Trial 1 Day7 | Trial 7 Day 6 | 0.223 | 1 | 1 | 1 |
| | After task | Within-day | Trial 7 Day1 | Trial 1 Day 1 | <0.001 *** | 0.005 ** | 0.139 | <0.001 *** |
| | | | Trial 7 Day2 | Trial 1 Day 2 | <0.001 *** | 0.031 * | 0.002 ** | 0.349 |
| | | | Trial 7 Day3 | Trial 1 Day 3 | 0.014 * | 0.727 | 0.055 | 0.482 |
| | | | Trial 7 Day4 | Trial 1 Day 4 | 0.027 * | 1 | 0.015 * | 1 |
| | | | Trial 7 Day5 | Trial 1 Day 5 | 0.051 | 1 | 0.605 | 0.798 |
| | | | Trial 7 Day6 | Trial 1 Day 6 | 0.068 | 1 | 0.605 | 0.311 |
| | | | Trial 7 Day7 | Trial 1 Day 7 | 0.483 | 1 | 0.605 | 1 |
| | | Next-day | Trial 1 Day2 | Trial 7 Day 1 | <0.001 *** | 1 | 0.933 | 1 |
| | | | Trial 1 Day3 | Trial 7 Day 2 | <0.001 *** | 1 | 0.043 * | 1 |
| | | | Trial 1 Day4 | Trial 7 Day 3 | 0.305 | 1 | 0.596 | 1 |
| | | | Trial 1 Day5 | Trial 7 Day 4 | -0.076 | 1 | 0.362 | 1 |
| | | | Trial 1 Day6 | Trial 7 Day 5 | 0.463 | 1 | 0.247 | 1 |
| | | | Trial 1 Day7 | Trial 7 Day 6 | 0.405 | 0.26 | 0.633 | 1 |

Supplementary Table 14: Statistics for Fig 4 K L pt1.

| Group | Time of treatment | Comparaison | Day | ANOVA | p-value | Sig. | Group 1 | Group 2 | p-value | Sig. |
|---|---|---|---|---|---|---|---|---|---|---|
| DCN | During task | Trial 1 | 1.00 | F(3,45)=0.6547 | 0.584 | n.s. | | | | |
| | | | 2.00 | F(3,45)=7.449 | <0.001 | *** | Sham + CNO | DREADD + CNO | 0.014 | * |
| | | | | | | | DREADD + SAL | DREADD + CNO | <0.001 | *** |
| | | | | | | | Sham + SAL | DREADD + CNO | <0.001 | *** |
| | | | 3.00 | F(3,45)=6.871 | <0.001 | *** | Sham + CNO | DREADD + CNO | 0.054 | n.s. |
| | | | | | | | DREADD + SAL | DREADD + CNO | <0.001 | *** |
| | | | | | | | Sham + SAL | DREADD + CNO | 0.001 | ** |
| | | | 4.00 | F(3,45)=6.415 | 0.001 | ** | Sham + CNO | DREADD + CNO | 0.013 | * |
| | | | | | | | DREADD + SAL | DREADD + CNO | 0.011 | * |
| | | | | | | | Sham + SAL | DREADD + CNO | <0.001 | *** |
| | | | 5.00 | F(3,45)=6.385 | 0.001 | ** | Sham + CNO | DREADD + CNO | 0.001 | ** |
| | | | | | | | DREADD + SAL | DREADD + CNO | 0.001 | ** |
| | | | | | | | Sham + SAL | DREADD + CNO | 0.012 | * |
| | | | 6.00 | F(3,45)=19.32 | <0.001 | *** | Sham + CNO | DREADD + CNO | <0.001 | *** |
| | | | | | | | DREADD + SAL | DREADD + CNO | 0.002 | ** |
| | | | | | | | Sham + SAL | DREADD + CNO | <0.001 | *** |
| | | | 7.00 | F(3,45)=6.82 | <0.001 | *** | Sham + CNO | DREADD + CNO | 0.005 | ** |
| | | | | | | | DREADD + SAL | DREADD + CNO | 0.01 | * |
| | | | | | | | Sham + SAL | DREADD + CNO | <0.001 | *** |
| | | Trial 7 | 1.00 | F(3,45)=2.609 | 0.063 | nn | | | | |
| | | | 2.00 | F(3,45)=2.668 | 0.058 | n.s. | | | | |
| | | | 3.00 | F(3,45)=1.819 | 0.157 | n.s. | | | | |
| | | | 4.00 | F(3,45)=2.761 | 0.053 | n.s. | | | | |
| | | | 5.00 | F(3,45)=4.429 | 0.008 | ** | Sham + CNO | DREADD + CNO | 0.056 | n.s. |
| | | | | | | | DREADD + SAL | DREADD + CNO | 0.042 | * |
| | | | | | | | Sham + SAL | DREADD + CNO | 0.004 | ** |
| | | | 6.00 | F(3,45)=8.879 | <0.001 | *** | Sham + CNO | DREADD + CNO | <0.001 | *** |
| | | | | | | | DREADD + SAL | DREADD + CNO | 0.016 | * |
| | | | | | | | Sham + SAL | DREADD + CNO | <0.001 | *** |
| | | | 7.00 | F(3,45)=3.304 | 0.028 | * | Sham + CNO | DREADD + CNO | 0.064 | n.s. |
| | | | | | | | DREADD + SAL | DREADD + CNO | 0.371 | n.s. |
| | | | | | | | Sham + SAL | DREADD + CNO | 0.023 | * |
| | After task | Trial 1 | 1.00 | F(3,30)=1.142 | 0.348 | n.s. | | | | |
| | | | 2.00 | F(3,30)=8.619 | <0.001 | *** | Sham + CNO | DREADD + CNO | <0.001 | *** |
| | | | | | | | DREADD + SAL | DREADD + CNO | 0.001 | ** |
| | | | | | | | Sham + SAL | DREADD + CNO | <0.001 | *** |
| | | | 3.00 | F(3,30)=5.689 | 0.003 | ** | Sham + CNO | DREADD + CNO | 0.007 | ** |
| | | | | | | | DREADD + SAL | DREADD + CNO | 0.001 | ** |
| | | | | | | | Sham + SAL | DREADD + CNO | 0.01 | * |
| | | | 4.00 | F(3,30)=1.947 | 0.143 | n.s. | | | | |
| | | | 5.00 | F(3,30)=3.323 | 0.032 | * | Sham + CNO | DREADD + CNO | 0.075 | n.s. |
| | | | | | | | DREADD + SAL | DREADD + CNO | 0.045 | * |
| | | | | | | | Sham + SAL | DREADD + CNO | 0.92 | n.s. |
| | | | 6.00 | F(3,30)=0.6632 | 0.581 | n.s. | | | | |
| | | | 7.00 | F(3,30)=0.5244 | 0.668 | n.s. | | | | |
| | | Trial 7 | 1.00 | F(3,30)=0.7346 | 0.539 | n.s. | | | | |
| | | | 2.00 | F(3,30)=4.102 | 0.014 | * | Sham + CNO | DREADD + CNO | 0.003 | ** |
| | | | | | | | DREADD + SAL | DREADD + CNO | 0.161 | n.s. |
| | | | | | | | Sham + SAL | DREADD + CNO | 0.086 | n.s. |
| | | | 3.00 | F(3,30)=0.6607 | 0.582 | n.s. | | | | |
| | | | 4.00 | F(3,30)=1.748 | 0.178 | n.s. | | | | |
| | | | 5.00 | F(3,30)=1.478 | 0.24 | n.s. | | | | |
| | | | 6.00 | F(3,30)=0.1749 | 0.912 | n.s. | | | | |
| | | | 7.00 | F(3,30)=0.8508 | 0.477 | n.s. | | | | |

Supplementary Table 15: Statistics for Fig 4 K L pt2.

| Group | Time of treatment | Day | ANOVA Treatnent | p-value | Sig. | ANOVA Treatnent:Trial | p-value | Sig. |
|---|---|---|---|---|---|---|---|---|
| Dentate – CL(Gi) | During task | 1.00 | F(1,18)=1.61 | 0.221 | n.s. | F(6,108)=0.5894 | 0.7382 | n.s. |
| | | 2.00 | F(1,18)=7.548 | 0.013 | * | F(6,108)=0.4718 | 0.828 | n.s. |
| | | 3.00 | F(1,18)=5.057 | 0.037 | * | F(6,108)=1.023 | 0.414 | n.s. |
| | | 4.00 | F(1,18)=10.28 | 0.004 | ** | F(6,108)=0.2491 | 0.9587 | n.s. |
| | | 5.00 | F(1,18)=8.577 | 0.008 | ** | F(6,108)=0.9642 | 0.453 | n.s. |
| | | 6.00 | F(1,18)=15.97 | <0.001 | *** | F(6,108)=0.5614 | 0.7602 | n.s. |
| | | 7.00 | F(1,18)=8.386 | 0.009 | ** | F(6,108)=0.5056 | 0.8029 | n.s. |
| | After task | 1.00 | F(1,17)=0.3552 | 0.559 | n.s. | F(6,102)=2.68 | 0.0186 | # |
| | | 2.00 | F(1,17)=0.5855 | 0.454 | n.s. | F(6,102)=1.179 | 0.3235 | n.s. |
| | | 3.00 | F(1,17)=4.418 | 0.051 | n.s. | F(6,102)=3.069 | 0.0084 | ## |
| | | 4.00 | F(1,17)=1.494 | 0.238 | n.s. | F(6,102)=0.773 | 0.5929 | n.s. |
| | | 5.00 | F(1,17)=1.3 | 0.27 | n.s. | F(6,102)=4.524 | <0.001 | ### |
| | | 6.00 | F(1,17)=0.01609 | 0.901 | n.s. | F(6,102)=0.3709 | 0.8959 | n.s. |
| | | 7.00 | F(1,17)=1.585 | 0.225 | n.s. | F(6,102)=1.124 | 0.3539 | n.s. |
| Interposed – VAL(Gi) | During task | 1.00 | F(1,19)=1.914 | 0.1826 | n.s. | F(6,114)=0.9818 | 0.441 | n.s. |
| | | 2.00 | F(1,19)=2.368 | 0.1403 | n.s. | F(6,114)=1.507 | 0.1819 | n.s. |
| | | 3.00 | F(1,19)=0.3523 | 0.5598 | n.s. | F(6,114)=0.7044 | 0.6467 | n.s. |
| | | 4.00 | F(1,19)=4.968 | 0.03809 | * | F(6,114)=0.7656 | 0.5985 | n.s. |
| | | 5.00 | F(1,19)=5.973 | 0.0245 | * | F(6,114)=0.5808 | 0.745 | n.s. |
| | | 6.00 | F(1,19)=17.28 | 0.000536 | *** | F(6,114)=1.797 | 0.1058 | n.s. |
| | | 7.00 | F(1,19)=9.661 | 0.006 | ** | F(6,114)=1.223 | 0.2996 | n.s. |
| | After task | 1.00 | F(1,16)=3.824 | 0.06821 | n.s. | F(6,96)=1.381, | 0.23 | n.s. |
| | | 2.00 | F(1,16)=2.827 | 0.1121 | n.s. | F(6,96)=2.635, | 0.02078 | # |
| | | 3.00 | F(1,16)=0.03395 | 0.8561 | n.s. | F(6,96)=2.015, | 0.07095 | n.s. |
| | | 4.00 | F(1,16)=0.06177 | 0.8069 | n.s. | F(6,96)=3.911, | 0.001543 | ## |
| | | 5.00 | F(1,16)=0.006183 | 0.938 | n.s. | F(6,96)=2.844, | 0.01361 | # |
| | | 6.00 | F(1,16)=0.1843 | 0.6734 | n.s. | F(6,96)=4.661, | <0.001 | ### |
| | | 7.00 | F(1,16)=2.812 | 0.113 | n.s. | F(6,96)=1.507, | 0.1839 | n.s. |

Supplementary Table 16: Statistics for Fig 5 BCEF.

| Group | Time of treatment | Comparaison | Moment 1 | Moment 2 | DREADD + CNO | DREADD + SAL |
|---|---|---|---|---|---|---|
| Dentate– CL(Gi) | During task | Within-day | Trial 7 Day1 | Trial 1 Day 1 | 0.080 | 0.062 |
| | | | Trial 7 Day2 | Trial 1 Day 2 | 0.007 ** | 0.313 |
| | | | Trial 7 Day3 | Trial 1 Day 3 | 0.563 | 0.436 |
| | | | Trial 7 Day4 | Trial 1 Day 4 | 0.563 | 0.139 |
| | | | Trial 7 Day5 | Trial 1 Day 5 | 0.130 | 0.139 |
| | | | Trial 7 Day6 | Trial 1 Day 6 | 0.145 | 0.436 |
| | | | Trial 7 Day7 | Trial 1 Day 7 | 0.563 | 0.436 |
| | | Next-day | Trial 1 Day2 | Trial 7 Day 1 | 0.897 | 1 |
| | | | Trial 1 Day3 | Trial 7 Day 2 | 0.897 | 1 |
| | | | Trial 1 Day4 | Trial 7 Day 3 | 0.897 | 1 |
| | | | Trial 1 Day5 | Trial 7 Day 4 | 0.893 | 1 |
| | | | Trial 1 Day6 | Trial 7 Day 5 | 0.003 ** | 1 |
| | | | Trial 1 Day7 | Trial 7 Day 6 | 0.897 | 1 |
| | After task | Within-day | Trial 7 Day1 | Trial 1 Day 1 | 0.008 ** | <0.001 *** |
| | | | Trial 7 Day2 | Trial 1 Day 2 | 0.985 | 0.075 |
| | | | Trial 7 Day3 | Trial 1 Day 3 | 0.194 | 1.000 |
| | | | Trial 7 Day4 | Trial 1 Day 4 | 0.056 | 0.231 |
| | | | Trial 7 Day5 | Trial 1 Day 5 | 0.985 | 1.000 |
| | | | Trial 7 Day6 | Trial 1 Day 6 | 0.985 | 0.576 |
| | | | Trial 7 Day7 | Trial 1 Day 7 | 0.576 | 0.576 |
| | | Next-day | Trial 1 Day2 | Trial 7 Day 1 | 0.50 | 0.637 |
| | | | Trial 1 Day3 | Trial 7 Day 2 | 1.00 | 0.148 |
| | | | Trial 1 Day4 | Trial 7 Day 3 | 0.80 | 0.069 |
| | | | Trial 1 Day5 | Trial 7 Day 4 | 0.34 | 0.645 |
| | | | Trial 1 Day6 | Trial 7 Day 5 | 1.00 | 0.645 |
| | | | Trial 1 Day7 | Trial 7 Day 6 | 0.15 | 0.457 |
| Interposed – VAL(Gi) | During task | Within-day | Trial 7 Day1 | Trial 1 Day 1 | <0.001 *** | 0.036 * |
| | | | Trial 7 Day2 | Trial 1 Day 2 | <0.001 *** | 0.021 * |
| | | | Trial 7 Day3 | Trial 1 Day 3 | 0.026 * | 0.036 * |
| | | | Trial 7 Day4 | Trial 1 Day 4 | 0.101 | 0.153 |
| | | | Trial 7 Day5 | Trial 1 Day 5 | 0.112 | 0.121 |
| | | | Trial 7 Day6 | Trial 1 Day 6 | 0.026 * | 0.153 |
| | | | Trial 7 Day7 | Trial 1 Day 7 | 0.149 | 0.153 |
| | | Next-day | Trial 1 Day2 | Trial 7 Day 1 | 0.018 * | 0.32 |
| | | | Trial 1 Day3 | Trial 7 Day 2 | 0.018 * | 0.25 |
| | | | Trial 1 Day4 | Trial 7 Day 3 | 0.070 | 0.60 |
| | | | Trial 1 Day5 | Trial 7 Day 4 | 0.053 | 0.27 |
| | | | Trial 1 Day6 | Trial 7 Day 5 | 0.053 | 0.58 |
| | | | Trial 1 Day7 | Trial 7 Day 6 | 0.404 | 0.32 |
| | After task | Within-day | Trial 7 Day1 | Trial 1 Day 1 | 0.028 * | <0.001 *** |
| | | | Trial 7 Day2 | Trial 1 Day 2 | 0.002 ** | 0.036 * |
| | | | Trial 7 Day3 | Trial 1 Day 3 | 0.020 * | 0.037 * |
| | | | Trial 7 Day4 | Trial 1 Day 4 | 0.028 * | 0.132 |
| | | | Trial 7 Day5 | Trial 1 Day 5 | 0.028 * | 0.003 ** |
| | | | Trial 7 Day6 | Trial 1 Day 6 | 0.028 * | 0.037 * |
| | | | Trial 7 Day7 | Trial 1 Day 7 | 0.028 * | 0.132 |
| | | Next-day | Trial 1 Day2 | Trial 7 Day 1 | 0.060 | 0.60 |
| | | | Trial 1 Day3 | Trial 7 Day 2 | 0.025 * | 0.21 |
| | | | Trial 1 Day4 | Trial 7 Day 3 | 0.054 | 0.60 |
| | | | Trial 1 Day5 | Trial 7 Day 4 | 0.060 | 0.60 |
| | | | Trial 1 Day6 | Trial 7 Day 5 | 0.060 | 0.37 |
| | | | Trial 1 Day7 | Trial 7 Day 6 | 0.003 ** | 0.23 |

Supplementary Table 17: Statistics for Fig 5 GHKL pt1.

| Group | Time of treatment | Comparaison | Day | Group 1 | Group 2 | p-value | Sig. |
|---|---|---|---|---|---|---|---|
| Dentate – CL(Gi) | During task | Trial 1 | 1.00 | DREADD + SAL | DREADD + CNO | 0.74 | n.s. |
| | | | 2.00 | DREADD + SAL | DREADD + CNO | 0.13 | n.s. |
| | | | 3.00 | DREADD + SAL | DREADD + CNO | 0.035 | * |
| | | | 4.00 | DREADD + SAL | DREADD + CNO | 0.09 | n.s. |
| | | | 5.00 | DREADD + SAL | DREADD + CNO | 0.026 | * |
| | | | 6.00 | DREADD + SAL | DREADD + CNO | 0.0097 | ** |
| | | | 7.00 | DREADD + SAL | DREADD + CNO | 0.039 | * |
| | | Trial 7 | 1.00 | DREADD + SAL | DREADD + CNO | 0.24 | n.s. |
| | | | 2.00 | DREADD + SAL | DREADD + CNO | 0.089 | n.s. |
| | | | 3.00 | DREADD + SAL | DREADD + CNO | 0.88 | n.s. |
| | | | 4.00 | DREADD + SAL | DREADD + CNO | 0.011 | * |
| | | | 5.00 | DREADD + SAL | DREADD + CNO | 0.0094 | ** |
| | | | 6.00 | DREADD + SAL | DREADD + CNO | 0.0042 | ** |
| | | | 7.00 | DREADD + SAL | DREADD + CNO | 0.041 | * |
| | After task | Trial 1 | 1.00 | DREADD + SAL | DREADD + CNO | 0.093 | n.s. |
| | | | 2.00 | DREADD + SAL | DREADD + CNO | 0.8 | n.s. |
| | | | 3.00 | DREADD + SAL | DREADD + CNO | 0.03 | * |
| | | | 4.00 | DREADD + SAL | DREADD + CNO | 0.94 | n.s. |
| | | | 5.00 | DREADD + SAL | DREADD + CNO | 0.21 | n.s. |
| | | | 6.00 | DREADD + SAL | DREADD + CNO | 0.48 | n.s. |
| | | | 7.00 | DREADD + SAL | DREADD + CNO | 0.29 | n.s. |
| | | Trial 7 | 1.00 | DREADD + SAL | DREADD + CNO | 0.27 | n.s. |
| | | | 2.00 | DREADD + SAL | DREADD + CNO | 0.11 | n.s. |
| | | | 3.00 | DREADD + SAL | DREADD + CNO | 0.67 | n.s. |
| | | | 4.00 | DREADD + SAL | DREADD + CNO | 0.18 | n.s. |
| | | | 5.00 | DREADD + SAL | DREADD + CNO | 0.66 | n.s. |
| | | | 6.00 | DREADD + SAL | DREADD + CNO | 0.89 | n.s. |
| | | | 7.00 | DREADD + SAL | DREADD + CNO | 0.34 | n.s. |
| Interposed – VAL(Gi) | During task | Trial 1 | 1.00 | DREADD + SAL | DREADD + CNO | 0.095 | n.s. |
| | | | 2.00 | DREADD + SAL | DREADD + CNO | 0.0058 | ** |
| | | | 3.00 | DREADD + SAL | DREADD + CNO | 0.12 | n.s. |
| | | | 4.00 | DREADD + SAL | DREADD + CNO | 0.045 | * |
| | | | 5.00 | DREADD + SAL | DREADD + CNO | 0.22 | n.s. |
| | | | 6.00 | DREADD + SAL | DREADD + CNO | 0.00027 | *** |
| | | | 7.00 | DREADD + SAL | DREADD + CNO | 0.063 | n.s. |
| | | Trial 7 | 1.00 | DREADD + SAL | DREADD + CNO | 0.26 | n.s. |
| | | | 2.00 | DREADD + SAL | DREADD + CNO | 0.66 | n.s. |
| | | | 3.00 | DREADD + SAL | DREADD + CNO | 0.43 | n.s. |
| | | | 4.00 | DREADD + SAL | DREADD + CNO | 0.19 | n.s. |
| | | | 5.00 | DREADD + SAL | DREADD + CNO | 0.021 | * |
| | | | 6.00 | DREADD + SAL | DREADD + CNO | 0.00027 | *** |
| | | | 7.00 | DREADD + SAL | DREADD + CNO | 0.0042 | ** |
| | After task | Trial 1 | 1.00 | DREADD + SAL | DREADD + CNO | 0.38 | n.s. |
| | | | 2.00 | DREADD + SAL | DREADD + CNO | 0.0016 | ** |
| | | | 3.00 | DREADD + SAL | DREADD + CNO | 0.067 | n.s. |
| | | | 4.00 | DREADD + SAL | DREADD + CNO | 0.034 | * |
| | | | 5.00 | DREADD + SAL | DREADD + CNO | 0.13 | n.s. |
| | | | 6.00 | DREADD + SAL | DREADD + CNO | 0.048 | * |
| | | | 7.00 | DREADD + SAL | DREADD + CNO | 0.017 | * |
| | | Trial 7 | 1.00 | DREADD + SAL | DREADD + CNO | 0.02 | * |
| | | | 2.00 | DREADD + SAL | DREADD + CNO | 0.88 | n.s. |
| | | | 3.00 | DREADD + SAL | DREADD + CNO | 0.68 | n.s. |
| | | | 4.00 | DREADD + SAL | DREADD + CNO | 0.31 | n.s. |
| | | | 5.00 | DREADD + SAL | DREADD + CNO | 0.087 | n.s. |
| | | | 6.00 | DREADD + SAL | DREADD + CNO | 0.7 | n.s. |
| | | | 7.00 | DREADD + SAL | DREADD + CNO | 0.9 | n.s. |

Supplementary Table 18: Statistics for Fig 5 GHKL pt2.

| Group | Time of treatment | Day | ANOVA Treatnent | p-value | Sig. | ANOVA Treatnent:Trial | p-value | Sig. |
|---|---|---|---|---|---|---|---|---|
| Dentate – CL(Gi) | During task | 8.00 | F(1,18)=6.786 | 0.02 | * | F(6,108)=0.3734 | 0.89 | n.s. |
| | | 9.00 | F(1,18)=3.659 | 0.07 | n.s. | F(6,108)=0.5489 | 0.77 | n.s. |
| Interposed – VAL(Gi) | During task | 8.00 | F(1,19)=3.106 | 0.09 | n.s. | F(6,114)=0.5508 | 0.77 | n.s. |
| | | 9.00 | F(1,19)=4.405 | 0.05 | * | F(6,114)=1.807 | 0.10 | n.s. |

Supplementary Table 19: Statistics for Fig 5 IM.

| Group | Time of treatment | Comparaison | Moment 1 | Moment 2 | SAL→CNO | CNO->SAL |
|---|---|---|---|---|---|---|
| Dentate – CL(Gi) | During task | Within-day | Trial 7 Day8 | Trial 1 Day 8 | 0.59 | 0.99 |
| | | | Trial 7 Day9 | Trial 1 Day 9 | 0.59 | 0.99 |
| | | Next-day | Trial 1 Day8 | Trial 7 Day 7 | 0.03 * | 0.67 |
| | | | Trial 1 Day9 | Trial 7 Day 8 | 0.40 | 0.68 |
| Interposed – VAL(Gi) | During task | Within-day | Trial 7 Day8 | Trial 1 Day 8 | 0.010 * | 0.003 ** |
| | | | Trial 7 Day9 | Trial 1 Day 9 | 0.008 ** | 0.298 |
| | | Next-day | Trial 1 Day8 | Trial 7 Day 7 | <0.001 *** | 0.756 |
| | | | Trial 1 Day9 | Trial 7 Day 8 | 0.005 ** | 0.015 * |

Supplementary Table 20: Statistics for Fig 5 JN pt1.

| Group | Time of treatment | Comparaison | Day | Group 1 | Group 2 | p-value | Sig. |
|---|---|---|---|---|---|---|---|
| Dentate – CL(Gi) | During task | Trial 1 | 7.00 | DREADD + SAL | DREADD + CNO | 0.039 | * |
| | | | 8.00 | DREADD + SAL | DREADD + CNO | 0.061 | n.s. |
| | | | 9.00 | DREADD + SAL | DREADD + CNO | 0.057 | n.s. |
| | | Trial 7 | 7.00 | DREADD + SAL | DREADD + CNO | 0.041 | * |
| | | | 8.00 | DREADD + SAL | DREADD + CNO | 0.017 | * |
| | | | 9.00 | DREADD + SAL | DREADD + CNO | 0.068 | n.s. |
| Interposed – VAL(Gi) | During task | Trial 1 | 7.00 | DREADD + SAL | DREADD + CNO | 0.063 | n.s. |
| | | | 8.00 | DREADD + SAL | DREADD + CNO | 0.006 | ** |
| | | | 9.00 | DREADD + SAL | DREADD + CNO | <0.001 | *** |
| | | Trial 7 | 7.00 | DREADD + SAL | DREADD + CNO | 0.004 | ** |
| | | | 8.00 | DREADD + SAL | DREADD + CNO | 0.19 | n.s. |
| | | | 9.00 | DREADD + SAL | DREADD + CNO | 0.8 | n.s. |

Supplementary Table 21: Statistics for Fig 5 JN pt2.

| Variable | Group | ANOVA | test | Estimate | CI 2.5% | CI 97.5% | p-val | sig. |
|---|---|---|---|---|---|---|---|---|
| Within day learning | pooled controls | Intercept: F(1,616)=187.3, p=0 *** | Acquisition=0 | 57.80 | 47.00 | 68.50 | <0.0001 | *** |
| | | | Consolidation=0 | 24.00 | 17.40 | 30.50 | <0.0001 | *** |
| | | | Maintenance=0 | 19.50 | 12.90 | 26.00 | <0.0001 | *** |
| | | Phase: F(2,616)=29.75, p=4.63e-13 *** | Consolidation-Acquisition=0 | -33.80 | -45.60 | -22.00 | <0.0001 | *** |
| | | | Maintenance-Acquisition=0 | -38.30 | -50.10 | -26.50 | <0.0001 | *** |
| | | | Maintenance-Consolidation=0 | -4.51 | -12.90 | 3.84 | 0.41 | n.s. |
| | DCN dreaddGi+CNO during task | Intercept: F(1,82)=40.42, p=1.095e-08 *** | Acquisition=0 | 48.10 | 28.70 | 67.50 | <0.0001 | *** |
| | | | Consolidation=0 | 33.90 | 21.10 | 46.80 | <0.0001 | *** |
| | | | Maintenance=0 | 13.90 | 1.09 | 26.80 | 0.029 | * |
| | | Phase: F(2,82)=9.788, p=0.0001541 *** | Consolidation-Acquisition=0 | -14.20 | -34.30 | 6.04 | 0.22665 | n.s. |
| | | | Maintenance-Acquisition=0 | -34.20 | -54.30 | -14.00 | 0.00022 | *** |
| | | | Maintenance-Consolidation=0 | -20.00 | -34.30 | -5.72 | 0.00303 | ** |
| | CAV(CL) dreaddGi+CNO during task | Intercept: F(1,58)=20.33, p=3.219e-05 *** | Acquisition=0 | 36.10 | 9.12 | 63.00 | 0.00432 | ** |
| | | | Consolidation=0 | 22.40 | 4.75 | 40.10 | 0.00758 | ** |
| | | | Maintenance=0 | 27.20 | 9.49 | 44.80 | 0.00077 | *** |
| | | Phase: F(2,58)=0.6496, p=0.526 | | | | | | |
| | CAV(VAL) dreaddGi+CNO during task | Intercept: F(1,58)=58.77, p=2.223e-10 *** | Acquisition=0 | 63.50 | 34.50 | 92.50 | <0.0001 | *** |
| | | | Consolidation=0 | 42.00 | 24.60 | 59.50 | <0.0001 | *** |
| | | | Maintenance=0 | 28.70 | 11.20 | 46.10 | 0.00026 | *** |
| | | Phase: F(2,58)=3.339, p=0.0424 * | Consolidation-Acquisition=0 | -21.50 | -53.60 | 10.60 | 0.257 | n.s. |
| | | | Maintenance-Acquisition=0 | -34.80 | -66.90 | -2.76 | 0.029 | * |
| | | | Maintenance-Consolidation=0 | -13.30 | -36.00 | 9.34 | 0.35 | n.s. |
| | DCN dreaddGi+CNO after task | Intercept: F(1,46)=110.3, p=8.338e-14 *** | Acquisition=0 | 62.50 | 36.90 | 88.10 | <0.0001 | *** |
| | | | Consolidation=0 | 49.20 | 34.40 | 63.90 | <0.0001 | *** |
| | | | Maintenance=0 | 29.20 | 14.50 | 44.00 | <0.0001 | *** |
| | | Phase: F(2,46)=4.632, p=0.0147 * | Consolidation-Acquisition=0 | -13.30 | -42.20 | 15.50 | 0.523 | n.s. |
| | | | Maintenance-Acquisition=0 | -33.30 | -62.20 | -4.43 | 0.019 | * |
| | | | Maintenance-Consolidation=0 | -20.00 | -40.40 | 0.45 | 0.057 - | n.s. |
| | CAV(CL) dreaddGi+CNO after task | Intercept: F(1,46)=35.44, p=3.407e-07 *** | Acquisition=0 | 32.90 | -0.77 | 66.60 | 0.0578 - | n.s. |
| | | | Consolidation=0 | 35.10 | 15.70 | 54.60 | <0.0001 | *** |
| | | | Maintenance=0 | 28.00 | 8.52 | 47.40 | 0.0018 | ** |
| | | Phase: F(2,46)=0.1977, p=0.8213 | | | | | | |
| | CAV(VAL) dreaddGi+CNO after task | Intercept: F(1,46)=77, p=2.19e-11 *** | Acquisition=0 | 34.40 | -9.24 | 78.00 | 0.17 | n.s. |
| | | | Consolidation=0 | 68.40 | 42.60 | 94.20 | <0.0001 | *** |
| | | | Maintenance=0 | 71.70 | 45.90 | 97.50 | <0.0001 | *** |
| | | Phase: F(2,46)=1.698, p=0.1944 | | | | | | |
| Overnight change | pooled controls | Intercept: F(1,513)=30.41, p=5.543e-08 *** | Acquisition=0 | -6.67 | -16.10 | 2.79 | 0.25 | n.s. |
| | | | Consolidation=0 | -8.88 | -14.80 | -2.97 | 0.001 | ** |
| | | | Maintenance=0 | -15.40 | -22.40 | -8.45 | <0.0001 | *** |
| | | Phase: F(2,513)=2.46, p=0.08644 | no posthoc test | | | | | |
| | DCN dreaddGi+CNO during task | Intercept: F(1,68)=22.65, p=1.055e-05 *** | Acquisition=0 | -27.50 | -47.80 | -7.18 | 0.0038 | ** |
| | | | Consolidation=0 | -26.60 | -40.20 | -13.00 | <0.0001 | *** |
| | | | Maintenance=0 | -14.60 | -30.20 | 0.97 | 0.0730 | n.s. |
| | | Phase: F(2,68)=1.657, p=0.1983 | | | | | | |
| | CAV(CL) dreaddGi+CNO during task | Intercept: F(1,48)=6.067, p=0.01741 * | Acquisition=0 | -2.34 | -31.60 | 27.00 | 0.9962 | n.s. |
| | | | Consolidation=0 | -12.20 | -31.80 | 7.46 | 0.3495 | n.s. |
| | | | Maintenance=0 | -31.50 | -53.90 | -9.02 | 0.0026 | ** |
| | | Phase: F(2,48)=2.807, p=0.07033 | | | | | | |
| | CAV(VAL) dreaddGi+CNO during task | Intercept: F(1,48)=35.06, p=3.308e-07 *** | Acquisition=0 | -29.90 | -53.70 | -6.00 | 0.0084 | ** |
| | | | Consolidation=0 | -28.40 | -42.90 | -14.00 | <0.0001 | *** |
| | | | Maintenance=0 | -23.70 | -41.00 | -6.43 | 0.0032 | ** |
| | | Phase: F(2,48)=0.1877, p=0.8295 | | | | | | |
| | DCN dreaddGi+CNO after task | Intercept: F(1,38)=50.89, p=1.624e-08 *** | Acquisition=0 | -57.80 | -77.00 | -38.60 | <0.0001 | *** |
| | | | Consolidation=0 | -21.20 | -32.70 | -9.72 | <0.0001 | *** |
| | | | Maintenance=0 | -16.10 | -29.90 | -2.31 | 0.016 | * |
| | | Phase: F(2,38)=10.29, p=0.0002687 *** | Consolidation-Acquisition=0 | 36.60 | 15.40 | 57.80 | 0.00017 | *** |
| | | | Maintenance-Acquisition=0 | 41.70 | 19.20 | 64.20 | <0.0001 | *** |
| | | | Maintenance-Consolidation=0 | 5.09 | -11.70 | 21.90 | 0.75558 | n.s. |
| | CAV(CL) dreaddGi+CNO after task | Intercept: F(1,38)=7.828, p=0.008032 ** | Acquisition=0 | 29.60 | -5.65 | 64.90 | 0.12893 | n.s. |
| | | | Consolidation=0 | -18.30 | -38.60 | 2.09 | 0.09329 | n.s. |
| | | | Maintenance=0 | -38.00 | -63.00 | -13.10 | 0.00082 | *** |
| | | Phase: F(2,38)=7.014, p=0.002555 ** | Consolidation-Acquisition=0 | -47.90 | -87.70 | -8.05 | 0.01346 | * |
| | | | Maintenance-Acquisition=0 | -67.60 | -110.00 | -25.40 | 0.00053 | *** |
| | | | Maintenance-Consolidation=0 | -19.70 | -51.20 | 11.80 | 0.30508 | n.s. |
| | CAV(VAL) dreaddGi+CNO after task | Intercept: F(1,38)=37.07, p=4.309e-07 *** | Acquisition=0 | -16.10 | -66.20 | 34.00 | 0.8271 | n.s. |
| | | | Consolidation=0 | -40.80 | -69.70 | -11.90 | 0.0023 | ** |
| | | | Maintenance=0 | -87.20 | -123.00 | -51.80 | <0.0001 | *** |
| | | Phase: F(2,38)=4.709, p=0.01489 * | Consolidation-Acquisition=0 | -24.70 | -81.30 | 31.90 | 0.561 | n.s. |
| | | | Maintenance-Acquisition=0 | -71.10 | -131.00 | -11.10 | 0.015 | * |
| | | | Maintenance-Consolidation=0 | -46.40 | -91.20 | -1.67 | 0.04 | * |

Supplementary Table 22: Statistics for Fig 6B Within day learning and Overnight change.

| Variable | Group | Group levels | ANOVA | Phase | Estimate | CI 2.5% | CI 97.5% | p-value | sig. |
|---|---|---|---|---|---|---|---|---|---|
| Within-day learning | DCN-dreaddGi treatment during task | SAL, CNO | Group: F(1,24)=6.643, p=0.01652 * | Acquisition | 3.72 | -26.10 | 33.50 | 0.97 | n.s. |
| | | | Group*Stage: F(2,152)=4.509, p=0.01252 * | Consolidation | -36.30 | -68.40 | -4.24 | 0.023 | * |
| | | | | Maintenance | -12.40 | -44.50 | 19.60 | 0.607 | n.s. |
| | CAV(CL)-dreaddGi treatment during task | SAL, CNO | Group: F(1,18)=0.01232, p=0.9128 | | | | | | |
| | | | Group*Stage: F(2,116)=0.4455, p=0.6416 | | | | | | |
| | CAV(VAL)-dreaddGi treatment during task | SAL, CNO | Group: F(1,19)=0.5344, p=0.4737 | | | | | | |
| | | | Group*Stage: F(2,122)=0.5932, p=0.5542 | | | | | | |
| | DCN-dreaddGi treatment after task | SAL, CNO | Group: F(1,16)=5.099, p=0.03826 * | Acquisition | 0.53 | -44.50 | 45.50 | 1 | n.s. |
| | | | Group*Stage: F(2,104)=0.6724, p=0.5127 | Consolidation | -26.40 | -77.80 | 25.00 | 0.41 | n.s. |
| | | | | Maintenance | -17.60 | -69.00 | 33.80 | 0.66 | n.s. |
| | CAV(CL)-dreaddGi treatment after task | SAL, CNO | Group: F(1,17)=0.6256, p=0.4399 | | | | | | |
| | | | Group*Stage: F(2,110)=2.829, p=0.06338 | | | | | | |
| | CAV(VAL)-dreaddGi treatment after task | SAL, CNO | Group: F(1,16)=10.06, p=0.005927 ** | Acquisition | 17.20 | -31.00 | 65.40 | 0.64 | n.s. |
| | | | Group*Stage: F(2,104)=2.77, p=0.0673 | Consolidation | -57.20 | -112.00 | -2.31 | 0.04 | * |
| | | | | Maintenance | -47.80 | -103.00 | 7.14 | 0.097 | n.s. |
| Overnight change | DCN-dreaddGi treatment during task | SAL, CNO | Group: F(1,24)=9.47, p=0.005161 ** | Acquisition | 22.80 | -7.89 | 53.40 | 0.18 | n.s. |
| | | | Group*Stage: F(2,126)=3.956, p=0.02157 * | Consolidation | 12.40 | -20.20 | 44.90 | 0.63 | n.s. |
| | | | | Maintenance | -19.40 | -53.90 | 15.10 | 0.36 | n.s. |
| | CAV(CL)-dreaddGi treatment during task | SAL, CNO | Group: F(1,18)=2.308, p=0.1461 | | | | | | |
| | | | Group*Stage: F(2,96)=0.01152, p=0.9885 | | | | | | |
| | CAV(VAL)-dreaddGi treatment during task | SAL, CNO | Group: F(1,19)=0.1592, p=0.6944 | | | | | | |
| | | | Group*Stage: F(2,101)=0.5409, p=0.5839 | | | | | | |
| | DCN-dreaddGi treatment after task | SAL, CNO | Group: F(1,16)=3.349, p=0.08596 | | | | | | |
| | | | Group*Stage: F(2,86)=2.57, p=0.08239 | | | | | | |
| | CAV(CL)-dreaddGi treatment after task | SAL, CNO | Group: F(1,17)=3.608, p=0.07462 | | | | | | |
| | | | Group*Stage: F(2,91)=1.571, p=0.2133 | | | | | | |
| | CAV(VAL)-dreaddGi treatment after task | SAL, CNO | Group: F(1,16)=10.24, p=0.005587 ** | Acquisition | 0.64 | -53.80 | 55.00 | 1 | n.s. |
| | | | Group*Stage: F(2,86)=2.131, p=0.1249 | Consolidation | 23.70 | -39.10 | 86.50 | 0.61 | n.s. |
| | | | | Maintenance | 57.10 | -9.48 | 124.00 | 0.1 | n.s. |

Supplementary Table 23: Statistics for Fig 6B Saline vs CNO.

| Group | Phase | Pearson's r | Slope [95%CI] | Slope=0 | x Intercept [95 % CI] | y Intercept [95 % CI] |
|---|---|---|---|---|---|---|
| Pooled controls | Acquisition | -0.57 | -1.29 [-1.79;-0.945] | * | 91.6 [ 77; 110] | 119 [ 101; 139] |
| | Consolidation | -0.44 | -0.973 [-1.26;-0.728] | * | 145 [ 137; 155] | 141 [ 112; 174] |
| | Maintenance | -0.44 | -1.36 [-1.74;-1.07] | * | 164 [ 159; 171] | 224 [ 178; 282] |
| CAV(CL)-dreaddGi SAL during task | Acquisition | -0.66 | -1.48 [-2.99;-0.722] | * | 72.6 [47.2; 114] | 107 [75.9; 142] |
| | Consolidation | -0.59 | -1.23 [-1.89;-0.845] | * | 149 [ 139; 162] | 183 [ 134; 269] |
| | Maintenance | -0.38 | -1.53 [-2.8;-0.789] | * | 179 [ 166; 199] | 273 [ 148; 484] |
| CAV(VAL)-deaddGi SAL offline | Acquisition | -0.62 | -1.17 [-2.21;-0.601] | * | 111 [78.7; 148] | 129 [90.7; 187] |
| | Consolidation | -0.32 | -0.669 [-1.53;-0.26] | * | 172 [ 138; 250] | 115 [63.4; 215] |
| | Maintenance | -0.46 | -1.03 [-1.8;-0.546] | * | 199 [ 176; 233] | 206 [ 126; 328] |
| CAV(CL)-dreaddGi CNO during task | Acquisition | -0.68 | -1.58 [-2.33;-0.24] | * | 59.2 [44.8; 191] | 93.5 [47.1; 136] |
| | Consolidation | -0.67 | -0.913 [-1.19;-0.5] | * | 115 [ 102; 137] | 105 [66.8; 133] |
| | Maintenance | -0.26 | -0.403 [-1.2;0.235] | n.s. | 162 [-264; 488] | 65.5 [ 11; 147] |
| CAV(VAL)-dreaddGi CNO offline | Acquisition | -0.69 | -0.785 [-1.39;0.0291] | n.s. | 89.8 [56.7; 170] | 70.5 [27.1;94.3] |
| | Consolidation | -0.20 | -1.55 [-12.2;11.2] | n.s. | 136 [ 2.8; 202] | 211 [-982;1.08e+03] |
| | Maintenance | -0.23 | -3.49 [ -26;13.9] | n.s. | 148 [4.38; 254] | 518 [-1.82e+03;3.63e+03] |

Supplementary Table 24: Statistics for Fig 6DEF (Deming regression).

| Group | Phase | Pearson's r | Slope | Slope=0 | x Intercept | y Intercept |
|---|---|---|---|---|---|---|
| Pooled controls | Acquisition | 0.68 | 1.12 [0.926;1.34] | * | 12.1 [0.658;20.9] | -13.6 [-26.9;-1.23] |
| | Consolidation | 0.53 | 1.12 [0.929;1.36] | * | 10.5 [4.75;15.5] | -11.7 [-19.7;-4.9] |
| | Maintenance | 0.64 | 0.871 [0.747;1.01] | * | 14.4 [7.84;20.5] | -12.6 [-18.9;-6.45] |
| CAV(CL)-dreaddGi SAL during task | Acquisition | 0.50 | 1.47 [0.468;3.21] | * | 10.1 [-104;35.9] | -14.8 [-108;50.8] |
| | Consolidation | 0.60 | 1.11 [0.825;1.75] | * | 0.929 [-12.9;13.2] | -1.03 [-18.2;11.7] |
| | Maintenance | 0.48 | 0.873 [0.545;1.46] | * | 14.7 [-4.56;31.7] | -12.8 [-31.1;3.76] |
| CAV(VAL)-deaddGi SAL offline | Acquisition | 0.90 | 1.04 [0.79;1.28] | * | 18.2 [1.99;29.1] | -18.9 [-36.6;-2.79] |
| | Consolidation | 0.51 | 1.36 [0.99;2.06] | * | 23.9 [13.5;32.7] | -32.5 [-54.3;-15.4] |
| | Maintenance | 0.70 | 0.953 [0.707;1.39] | * | 28 [11.6;40.8] | -26.7 [-49.8;-11.5] |
| CAV(CL)-dreaddGi CNO during task | Acquisition | 0.52 | 1.17 [0.223;9.19] | * | 7.28 [-35.7;49.2] | -8.54 [-402;20.8] |
| | Consolidation | 0.43 | 0.958 [0.183;3.52] | * | 11.7 [-9.3;30.1] | -11.2 [-87.3;9.41] |
| | Maintenance | 0.23 | 0.752 [-12.7;14.5] | n.s. | 31.6 [13.6;75.7] | -23.8 [-424; 454] |
| CAV(VAL)-dreaddGi CNO offline | Acquisition | 0.78 | 1.2 [0.538;2.19] | * | 19.2 [1.87;35.4] | -23.1 [-57.2;-1.99] |
| | Consolidation | 0.26 | 0.604 [-0.433; 2] | n.s. | 22.7 [-354; 588] | -13.7 [-108;66.5] |
| | Maintenance | 0.29 | 0.687 [-6.37;5.78] | n.s. | 96.6 [-43.5; 184] | -66.4 [-316; 360] |

Supplementary Table 25: Statistics for Fig 6GHI (Deming regression).

| Group | Day | Phase | Factor | ANOVA | p-value | Sig. | Group 1 | Group 2 | Statistic | CI 2.5% | CI 97.5% | p-value | Cohen's D | Sig. |
|---|---|---|---|---|---|---|---|---|---|---|---|---|---|---|
| DCN | 1.00 | Acquisition | Treatment | F(1,30)=2.102 | 0.157 | n.s. | | | | | | | | |
| | | | Moment | F(1,30)=28.229 | <0.001 | *** | | | | | | | | |
| | | | Treatment:Moment | F(1,30)=0.87 | 0.358 | n.s. | | | | | | | | |
| | 4.00 | Consolidation | Treatment | F(1,30)=0.493 | 0.488 | n.s. | | | | | | | | |
| | | | Moment | F(1,30)=14.512 | <0.001 | *** | | | | | | | | |
| | | | Treatment:Moment | F(1,30)=0.595 | 0.446 | n.s. | | | | | | | | |
| | 7.00 | Maintenance | Treatment | F(1,30)=0.131 | 0.72 | n.s. | | | | | | | | |
| | | | Moment | F(1,30)=0.998 | 0.326 | n.s. | | | | | | | | |
| | | | Treatment:Moment | F(1,30)=8.61 | 0.006 | ** | Before-SAL | Before-CNO | 1.82 | -0.10 | 1.10 | 0.10 | 0.85 | n.s. |
| | | | | | | | After-SAL | After-CNO | -1.38 | -1.63 | 0.35 | 0.19 | -0.66 | n.s. |
| Sham | 1.00 | Acquisition | Treatment | F(1,32)=5.663 | 0.023 | * | | | | | | | | |
| | | | Moment | F(1,32)=251.757 | <0.001 | *** | | | | | | | | |
| | | | Treatment:Moment | F(1,32)=0.078 | 0.782 | n.s. | Before-SAL | Before-CNO | -0.66 | -1.91 | 1.01 | 0.52 | -0.31 | n.s. |
| | | | | | | | After-SAL | After-CNO | -0.54 | -1.76 | 1.04 | 0.59 | -0.26 | n.s. |
| | 4.00 | Consolidation | Treatment | F(1,32)=3.874 | 0.058 | n.s. | | | | | | | | |
| | | | Moment | F(1,32)=45.499 | <0.001 | *** | | | | | | | | |
| | | | Treatment:Moment | F(1,32)=0.153 | 0.698 | n.s. | | | | | | | | |
| | 7.00 | Maintenance | Treatment | F(1,32)=0.761 | 0.39 | n.s. | | | | | | | | |
| | | | Moment | F(1,32)=27.223 | <0.001 | *** | | | | | | | | |
| | | | Treatment:Moment | F(1,32)=1.4 | 0.245 | n.s. | | | | | | | | |

Supplementary Table 26: Statistics for Fig Sup 2 A.

| Group | Factor | ANOVA | p-value | Sig. | Group 1 | Group 2 | Statistic | CI 2.5% | CI 97.5% | p-value | Cohen's D | Sig. |
|---|---|---|---|---|---|---|---|---|---|---|---|---|
| DCN | Treatment | F(1,106)=7.96 | 0.006 | ** | | | | | | | | |
| | Speed | F(1,106)=201.619 | <0.001 | *** | | | | | | | | |
| | Treatment:Speed | F(1,106)=1.039 | 0.31 | n.s. | 5 rpm-SAL | 5 rpm-CNO | -0.449 | -12.68 | 8.25 | 0.66 | -0.18 | n.s. |
| | | | | | 10 rpm-SAL | 10 rpm-CNO | -1.369 | -35.51 | 7.91 | 0.19 | -0.54 | n.s. |
| | | | | | 15 rpm-SAL | 15 rpm-CNO | -1.413 | -84.23 | 16.33 | 0.17 | -0.58 | n.s. |
| | | | | | 20 pm-SAL | 20 rpm-CNO | -1.76 | -106.22 | 9.76 | 0.10 | -0.78 | n.s. |
| | | | | | 25 rpm-SAL | 25 rpm-CNO | Statistic | -30.56 | 3.46 | 0.11 | -0.83 | n.s. |
| Sham | Treatment | F(1,76)=1.836 | 0.179 | n.s. | | | | | | | | |
| | Speed | F(1,76)=152.296 | <0.001 | *** | | | | | | | | |
| | Treatment:Speed | F(1,76)=0.109 | 0.742 | n.s. | | | | | | | | |

Supplementary Table 27: Statistics for Fig Sup 2 B.

| Experiment | Value | ANOVA Group | p-value | Sig. |
|---|---|---|---|---|
| Horizontal Bar | Latency | F(3,28)=2.542 | 0.08 | n.s. |
| Vertical Pole | Latency | F(3,28)=1.717 | 0.19 | n.s. |
| Grid test | Latency | F(3,34)=0.197 | 0.90 | n.s. |
| Footprint | Linear movement | F(3,28)=0.392 | 0.76 | n.s. |
| | Sigma | F(3,28)=0.491 | 0.69 | n.s. |
| | Alternate coefficient | F(3,28)=1.022 | 0.40 | n.s. |

Supplementary Table 28: Statistics for Fig Sup 2 CDEF.

| Group | Day | Phase | Factor | ANOVA | p-value | Sig. | Group 1 | Group 2 | Statistic | CI 2.5% | CI 97.5% | p-value | Cohen's D | Sig. |
|---|---|---|---|---|---|---|---|---|---|---|---|---|---|---|
| Dentate-CL | 1.00 | Acquisition | Treatment | F(1,44)=1.443 | 0.236 | n.s. | | | | | | | | |
| | | | Moment | F(1,44)=169.334 | <0.001 | *** | | | | | | | | |
| | | | Treatment:Moment | F(1,44)=2.457 | 0.124 | n.s. | | | | | | | | |
| | 4.00 | Consolidation | Treatment | F(1,44)=15.56 | <0.001 | *** | | | | | | | | |
| | | | Moment | F(1,44)=74.842 | <0.001 | *** | | | | | | | | |
| | | | Treatment:Moment | F(1,44)=16.594 | <0.001 | *** | Before-SAL | Before-CNO | 2.73 | 0.42 | 3.08 | 0.01 | 1.10 | * |
| | | | | | | | After-SAL | After-CNO | -0.05 | -1.14 | 1.08 | 0.96 | -0.02 | n.s. |
| | 7.00 | Maintenance | Treatment | F(1,44)=2.483 | 0.122 | n.s. | | | | | | | | |
| | | | Moment | F(1,44)=0.033 | 0.857 | n.s. | | | | | | | | |
| | | | Treatment:Moment | F(1,44)=2.31 | 0.136 | n.s. | | | | | | | | |
| Interposed-VAL | 1.00 | Acquisition | Treatment | F(1,36)=16.74 | <0.001 | *** | | | | | | | | |
| | | | Moment | F(1,36)=65.574 | <0.001 | *** | | | | | | | | |
| | | | Treatment:Moment | F(1,36)=4.745 | 0.036 | * | Before-SAL | Before-CNO | -2.61 | -3.61 | -0.36 | 0.02 | -1.17 | * |
| | | | | | | | After-SAL | After-CNO | -0.98 | -1.95 | 0.74 | 0.35 | -0.44 | n.s. |
| | 4.00 | Consolidation | Treatment | F(1,36)=0.197 | 0.66 | n.s. | | | | | | | | |
| | | | Moment | F(1,36)=35.692 | <0.001 | *** | | | | | | | | |
| | | | Treatment:Moment | F(1,36)=10.455 | 0.003 | ** | Before-SAL | Before-CNO | -1.78 | -1.31 | 0.11 | 0.09 | -0.80 | n.s. |
| | | | | | | | After-SAL | After-CNO | 0.89 | -0.62 | 1.53 | 0.39 | 0.40 | n.s. |
| | 7.00 | Maintenance | Treatment | F(1,36)=20.015 | <0.001 | *** | | | | | | | | |
| | | | Moment | F(1,36)=89.27 | <0.001 | *** | | | | | | | | |
| | | | Treatment:Moment | F(1,36)=0.61 | 0.44 | n.s. | Before-SAL | Before-CNO | -1.79 | -1.79 | 0.14 | 0.09 | -0.80 | n.s. |
| | | | | | | | After-SAL | After-CNO | -1.58 | -1.36 | 0.20 | 0.13 | -0.71 | n.s. |

Supplementary Table 29: Statistics for Fig Sup 3 A.

| Group | Factor | ANOVA | p-value | Sig. | Group 1 | Group 2 | Statistic | CI 2.5% | CI 97.5% | p-value | Cohen's D | Sig. |
|---|---|---|---|---|---|---|---|---|---|---|---|---|
| Dentate-CL | Treatment | F(1,106)=0.416 | 0.52 | n.s. | | | | | | | | |
| | Speed | F(1,106)=116.649 | <0.001 | *** | | | | | | | | |
| | Treatment:Speed | F(1,106)=0.007 | 0.934 | n.s. | | | | | | | | |
| Interposed-VAL | Treatment | F(1,96)=17.495 | <0.001 | *** | | | | | | | | |
| | Speed | F(1,96)=132.095 | <0.001 | *** | | | | | | | | |
| | Treatment:Speed | F(1,96)=1.024 | 0.314 | n.s. | 5 rpm-SAL | 5 rpm-CNO | 0.88 | -23.27 | 53.27 | 0.40 | 0.39 | n.s. |
| | | | | | 10 rpm-SAL | 10 rpm-CNO | 1.21 | -17.12 | 56.52 | 0.26 | 0.54 | n.s. |
| | | | | | 15 rpm-SAL | 15 rpm-CNO | 3.81 | 22.59 | 82.81 | 0.00 | 1.70 | ** |
| | | | | | 20 pm-SAL | 20 rpm-CNO | 2.37 | 5.59 | 94.81 | 0.03 | 1.06 | * |
| | | | | | 25 rpm-SAL | 25 rpm-CNO | 2.04 | -2.58 | 58.78 | 0.07 | 0.91 | n.s. |

Supplementary Table 30: Statistics for Fig Sup 3 B.

| Experiment | Value | ANOVA Group | p-value | Sig. |
|---|---|---|---|---|
| Horizontal Bar | Latency | F(3,33)=0.826 | 0.49 | n.s. |
| Vertical Pole | Latency | F(3,33)=0.968 | 0.42 | n.s. |
| Grid test | Latency | F(3,33)=0.305 | 0.82 | n.s. |
| Footprint | Linear movement | F(3,31)=0.887 | 0.46 | n.s. |
| | Sigma | F(3,31)=0.103 | 0.96 | n.s. |
| | Alternate coefficient | F(3,31)=0.476 | 0.70 | n.s. |

Supplementary Table 31: Statistics for Fig Sup 3 CDEF.
